# Lost in a Large EEG Multiverse? Comparing Sampling Approaches for Representative Pipeline Selection

**DOI:** 10.1101/2025.04.08.647779

**Authors:** Cassie Ann Short, Andrea Hildebrandt, Robin Bosse, Stefan Debener, Metin Özyağcılar, Katharina Paul, Jan Wacker, Daniel Kristanto

**Affiliations:** Carl von Ossietzky Universität Oldenburg, Department of Psychology, Oldenburg, Germany; Universität Hildesheim, Institute of Psychology, Hildesheim, Germany; Universität Hamburg, Institute of Psychology, Hamburg, Germany

## Abstract

The multiplicity of defensible strategies for processing and analysing data has been implicated as a core contributor to the replicability crisis, creating uncertainty about the robustness of a result to variations in data processing choices. This issue is exacerbated where a large number of data processing pipelines are defensible, and where there is great heterogeneity in the pipelines applied in the literature, such as in processing and analysing electroencephalography (EEG) signals. In a multiverse analysis, equally defensible pipelines are computed and the robustness of the result to these variations is reported. However, a large number of defensible pipelines is sometimes infeasible to compute exhaustively, and researchers rely on sampling approaches. In these cases, pipelines are sampled from the full multiverse and the robustness is reported across these samples, assuming that they are representative for the entire multiverse. However, different sampling methods may yield different robustness results, introducing what we term multiverse sampling uncertainty. To illustrate, we computed a 528-pipeline multiverse analysis on EEG-recordings during an emotion classification task aiming to predict extraversion scores from the Late Positive Potential. We applied three sampling methods (random, stratified, and active learning) to sample 26 pipelines (5%), and evaluated the results in terms of the representativeness of the distribution of model fits to that of the full multiverse. Our results highlight variability in the representativeness of the distribution of model fits between samples. The active learning sample most closely represented the median model fit of the full multiverse. The need for representative pipeline sampling to mitigate bias in large multiverse analyses is discussed.

## Introduction

The flexibility that researchers have in selecting data processing and analysis options from multiple defensible alternatives, referred to as ‘researcher degrees of freedom’ (Simmons et al., 2011), has been identified as a key contributor to the replicability crisis (e.g., Ioannidis, 2005; Open Science Collaboration, 2015). Reporting only the results obtained using a single data processing pipeline creates a theoretical multiple comparisons problem for statistical tests that were not explicitly performed (Gelman & Loken, 2013), may increase false positive findings (Rubin, 2017), inflate effect sizes (Ioannidis, 2005; Simmons et al., 2011), and create uncertainty about the robustness of the single reported estimate (Hoffman et al., 2021). Researcher degrees of freedom is particularly exacerbated in neuroimaging studies, including electroencephalogram (EEG) signal, due to the many defensible combinations of options available along the data processing workflow to extract the signal of interest from the raw recording (e.g., Paul, Short et al., 2022). Different defensible EEG data processing pipelines can lead to heterogeneous results (e.g., Beauducel et al., 2024; Paul et al., 2024; Šoškić et al., 2024), enhance or diminish effects (e.g., Sadus et al., 2023; Schubert et al., 2022), suppress amplitude differences in transcranial magnetic stimulation evoked potentials (Rogasch et al., 2022), and influence data quality (Clayson et al., 2021). To minimise the risk of overestimating the precision and robustness of an estimate, it is necessary to explore the uncertainty introduced by researcher degrees of freedom in data processing and estimate its impact.

Multiverse analysis is a methodological approach to report uncertainty by systematically identifying and computing all pipelines that are equally defensible for answering a research question of interest (Steegen et al., 2016), and reporting the distribution of results across all pipelines (Simonsohn et al., 2020). This approach to communicate uncertainty is gaining popularity across many scientific fields (Short et al., 2025). In EEG research, various strategies have been implemented to identify the defensible multiverse of data processing pipelines for a given research question, including systematic literature reviews (e.g., Jacobsen et al., 2024), expert collaboration (e.g., Wacker, 2017), and by multiple experts independently selecting one pipeline each (e.g., Trübutschek et al., 2024). While such approaches have the potential to facilitate a transparent and systematic assessment of robustness to data processing uncertainty, the number of pipelines identified as defensible may be too large to compute exhaustively. For example, following the expert collaboration approach (Wacker, 2017), the CoScience Project reported that 18 million pipelines were agreed as defensible for EEG signal processing, prior to quantification and analysis, to answer the same research question from the same raw dataset (Paul, Short et al., 2022). Such huge multiverses require post-hoc reduction to minimise reduce the computational burden while preserving transparency. Pipeline sampling is the solution in such cases.

The post-hoc pipeline sampling approaches applied across the multiverse analysis literature are varied. The most prominent approaches include random sampling (e.g., Simonsohn et al., 2020), stratified random sampling (e.g., Beauducel et al., 2024; Paul et al., 2024), and active learning sampling (e.g., Dafflon et al., 2022; Kristanto et al., 2023). While each of these sampling approaches aims to represent the distribution of results across the entire population of defensible pipelines, each applies a functionally distinct procedure to achieve this goal. Random sampling selects pipelines independently and without replacement, with each pipeline having an equal chance of being selected based on the uniform probability distribution across all pipelines. While this approach is computationally efficient and requires no prior knowledge of the structural properties of pipelines, it may fail to capture the full distribution of potentially influential structural diversity, especially when sampling a small subset of a large multiverse. Stratified random sampling divides the pipelines into strata, which can ensure that every option within each decision node is represented in the sample of pipelines if the sample size allows. While this captures diversity at the level of decision nodes or options in isolation, it does not guarantee representation of potentially influential interactions between options across decision nodes. Active learning sampling (Dafflon et al., 2022) uses Bayesian optimisation with Gaussian process regression on a low-dimensional representation of the full multiverse created from pairwise similarities of the pipelines. This method iteratively selects pipelines, and uses the sample to estimate the full multiverse of pipelines that were not directly sampled. While this approach estimates the full distribution of results across all pipelines, it is more computationally intensive than the random and stratified sampling approaches, and may introduce bias if the initial randomly selected subset of burn-in pipelines is not diverse enough.

Given the functional differences between the multiverse sampling approaches, they may produce pipeline samples that differ in how accurately they represent the full defensible multiverse. Uncertainty about how well the distribution of estimates from a sampled set of pipelines reflects the population of pipelines raises uncertainty about the potential bias and the replicability of the multiverse analysis results. This source of uncertainty is in addition to those previously acknowledged (Hoffman et al., 2021), and we term this *multiverse sampling uncertainty*. Reporting multiverse sampling uncertainty is essential to strengthen the reliability and generalisability of multiverse analysis results where sampling is required.

Two approaches to report multiverse sampling uncertainty include (1) sample selection without explicit comparisons, with reference to previously published systematic comparisons of the representativeness of samples obtained using different sampling approaches, performed on data with similar characteristics and structure, and (2) sample selection with explicit comparisons, where multiple pipeline samples are computed using different sampling approaches, and they are compared for their representativeness of the full multiverse using a subset of cases for computational feasibility. The first approach may lack precision in the reported multiverse sampling uncertainty, especially given the possibility that influential decisions in multiverse analyses may not replicate across samples despite design similarities (Olsson-Collentine et al., 2023), but it requires little additional time or resources. The second approach provides data- and multiverse-specific precision in the reported multiverse sampling uncertainty, but requires substantial additional time and resources. However, the additional time and resources required can be reduced if software was available for sampling and output generation. To date, no published systematic comparisons exist to guide the first approach, nor does an open and easy-to-implement tool exist to ease the time and computational resources required for the second approach.

Firstly, we describe an open-source Multiverse Sampling Tool that enables comparison of pipeline samples obtained by random, stratified and active learning sampling in an automated workflow. The tool compares these samples based on their representation of model fit and spatial distribution of pipelines, providing a transparent means of reporting multiverse sampling uncertainty when exhaustive computation of the full multiverse is not feasible. Secondly, we use the Multiverse Sampling Tool on an EEG individual differences multiverse use case, offering the first systematic comparison of the representativeness of pipeline samples obtained through different sampling approaches. This comparison serves as a benchmark to guide the selection of a sampling approach in future large-scale EEG multiverse analyses with similar characteristics.

## Methods

The methods are organised into three sections. In section one, we describe the EEG multiverse use case in detail. In section two, we describe the three sampling approaches implemented in the Multiverse Sampling Tool. In section three, we describe the various elements of the Multiverse Sampling Tool and highlight modification options for different use cases.

### 1. Use Case Multiverse

#### Summary

The multiverse analysis use case aims to fit a model that predicts self-reported trait extraversion using mean amplitudes of the Late Positive Potential (LPP). The LPP is an event-related sustained positive amplitude deflection typically observed from approximately 250 ms post-stimulus at centro-parietal scalp sites, and the magnitude of the amplitude is sensitive to the emotional salience of the stimuli (e.g., Hajcak & Olvet, 2008; Speed et al., 2015). The LPP has previously been successfully quantified in the present dataset (Recio et al., 2017), demonstrating the feasibility of obtaining a theoretically relevant estimate of the LPP from this data. However, the broad morphology of the LPP, which may encompass overlapping cognitive processes, combined with potential individual and condition-specific variability (Gray, 1991), highlights the need for robustness analysis. The theoretical framework linking trait extraversion to heightened LPP responses to emotionally arousing stimuli (Eysenck, 1967; Speed et al., 2015) has led to a series of heterogeneous findings in terms of the specificity of emotional valence (e.g., Ku et al., 2020; Speed et al., 2015). This remains a substantive research interest in the field of EEG-related individual differences research. Given the psychometric relevance of the LPP in this context and the diverse methods for its quantification in the existing literature, the quantification of the LPP to emotionally salient stimuli is an ideal candidate for multiverse analysis.

#### Data

The data used were part of a larger dataset described in detail by Recio et al. (2017). Below, we describe the data that were used in the present analysis.

##### Participants

As reported by Recio et al. (2017), 269 young adults (52.4% female) with a mean age of 25.9 years (*SD* = 5.9) with normal or corrected-to-normal vision participated in the psychometric part of a larger study. Of these 269 individuals, 110 agreed to participate in the psychophysiological part of the experiment where EEG was recorded. Following exclusion from data analysis due to less than 25% viable trials within a condition following EEG data preprocessing (described below), a total of 98 participants aged between 18 and 38 years (*M* = 26.58, *SD* = 4.80, 46.51% female) were included in the final analyses. The original study was approved by a local ethics committee (Recio et al., 2017). Participants provided written informed consent prior to participation and were paid for their contribution.

##### Materials and procedure

The data analysed in the present study were taken from a larger dataset previously acquired using several face cognition tasks across two sessions: a psychometric and a psychophysiological session (Recio et al., 2017). Specifically, we analysed data from one questionnaire completed in the psychometric session, and data from one task from the psychophysiological session.

###### Questionnaire

Participants completed the German version of the extraversion and agreeableness items of the NEO Personality Inventory Revised (NEO-PI-R; Ostendorf & Angleitner, 2004; after Costa & McCrae, 1992), which is a self-report measure of the ‘Big Five’ personality traits as defined in the Five-Factor-Model of personality (FFM; Digman, 1990). The NEO-PI-R items require responses on a 5-point Likert scale ranging from strongly agree to strongly disagree. Here, we only analyse the extraversion score.

###### Task

Participants completed an emotion classification task previously described by Recio et al. (2017). The stimuli were dynamic images of facial expressions, obtained from the Radbound Faces Database (Langer et al., 2010) and morphed using the FantaMorph software (Abrosoft, 2010). Specifically, colour video clips of 38 models displaying dynamic facial expressions across six emotion conditions (anger, disgust, fear, happiness, sadness, and surprise) at two intensity levels (moderate and full) and an emotionally neutral condition (blinking, chewing) were presented. The intensity manipulation provided an elevation of task difficulty to prevent ceiling effects typically observed for recognising high intensity expressions (Recio et al., 2017). Dynamic expressions were used rather than static images to improve ecological validity (e.g., Recio et al., 2011). The task consisted of 798 trials, with 114 trials per condition (57 per intensity), with a short break after every 200 trials. Each video clip lasted 600 ms (200 ms dynamic followed by 400 ms static), at 30 frames per second, and each was preceded by a fixation cross for 700 ms. After each trial, the emotion categories displayed throughout the task were presented as text in German, and participants were asked to classify which emotion category the stimulus in the previous trial represented They selected the appropriate category with a mouse click. There was no time limit for the response.

###### EEG data acquisition

EEG data were recorded using 42 electrodes using a 64 channel Brain Products Amplifier (Brain Products GmbH). All electrodes were referenced online to the left mastoid, AFz served as the ground, the signal was filtered online with a 0.032-70 Hz bandpass filter, and sampled at 1000 Hz. Electrooculogram was recorded at the left and right external canthi and below the right eye. The electrode impedance was kept below 5kΩ.

#### Data Preprocessing

##### Extraversion

Extraversion was quantified as the average score of items within the extraversion subscale. The sample mean extraversion score was 2.26 (*SD* = 0.44).

##### EEG Data

All EEG data preprocessing and LPP quantification was performed offline in MATLAB R2022b (The MathWorks, Inc., 2022), using the EEGLab toolbox (version 2020.0; Delorme & Makeig, 2004) for part of the procedure. All scripts can be downloaded from [here](https://github.com/cassiesh/MultiverseSamplingTool). Initial preparation steps comprised of deleting irrelevant auxiliary channels, adding a channel for the left mastoid (which had served as an online reference), and correcting a 150 ms delay in event markers. Trials containing incorrect responses and trials containing correct responses given in under 200 ms post-stimulus were excluded from further processing.

The data were resampled to 500 Hz and re-referenced to Cz for cleaning. A high-pass linear (zero-phase) non-causal FIR filter with a cut-off frequency of 1 Hz was applied using pop_eegfiltnew(), which estimates the transition bandwidths and orders automatically. Following this, a standard deviation map of the voltage of each channel across the task duration in time windows of 5000 data points was computed for each participant. These maps were used to manually identify bad channels. Bad channels were classified as exhibiting sustained elevated standard deviation values (i.e., relative to the participant’s overall standard deviation range, involving clusters of time bins where standard deviation values exceeded the local baseline by a substantial margin (e.g., shifting from blue to yellow/red tones on the individual heatmap colour scale), across multiple time bins, with the exception of electrodes Fp1, FPz, Fp2, AF7, and AF8, as ocular artifacts were to remain in the data at this point in the procedure. Of the 102 participants screened at this stage, 66 had no bad channels identified, and 36 had bad channels identified (range = 1-5 channels, *M* = 0.54, *SD* = 1.02). The heatmaps for each participant can be viewed [here](https://github.com/cassiesh/MultiverseSamplingTool/tree/main/UseCaseMultiverse_MATLABScripts/StdDevImages). Following the removal of bad channels, Independent Component Analysis (ICA) was performed and artifacts were classified using ICLabel (Pion-Tonachini, 2019). Components classified as ocular artifacts that met a threshold of 0.85 probability were rejected. The data were low pass filtered using a linear (zero-phase) non-causal FIR filter with a cut-off frequency of 30 Hz, and previously rejected bad channels were interpolated using spherical spline interpolation. The data were epoched at −200 to 800 ms around the stimuli.

##### Multiverse Analysis

We quantified the mean LPP amplitude across 528 pipelines. The pipelines were defined by four forked decision nodes: baseline duration (two options), offline reference scheme (three options), time window (11 options), and channel clusters (eight options). The selected options were guided by the CoScience Project LPP quantification multiverse (Hennig et al., 2021), and previous knowledge of the current dataset, and are explained in Supplementary 6. Each option at each decision node was combined with each option from each of the other decision nodes, resulting in a Cartesian product of 528 combinations, illustrated in Figure 1.

**Figure 1.**
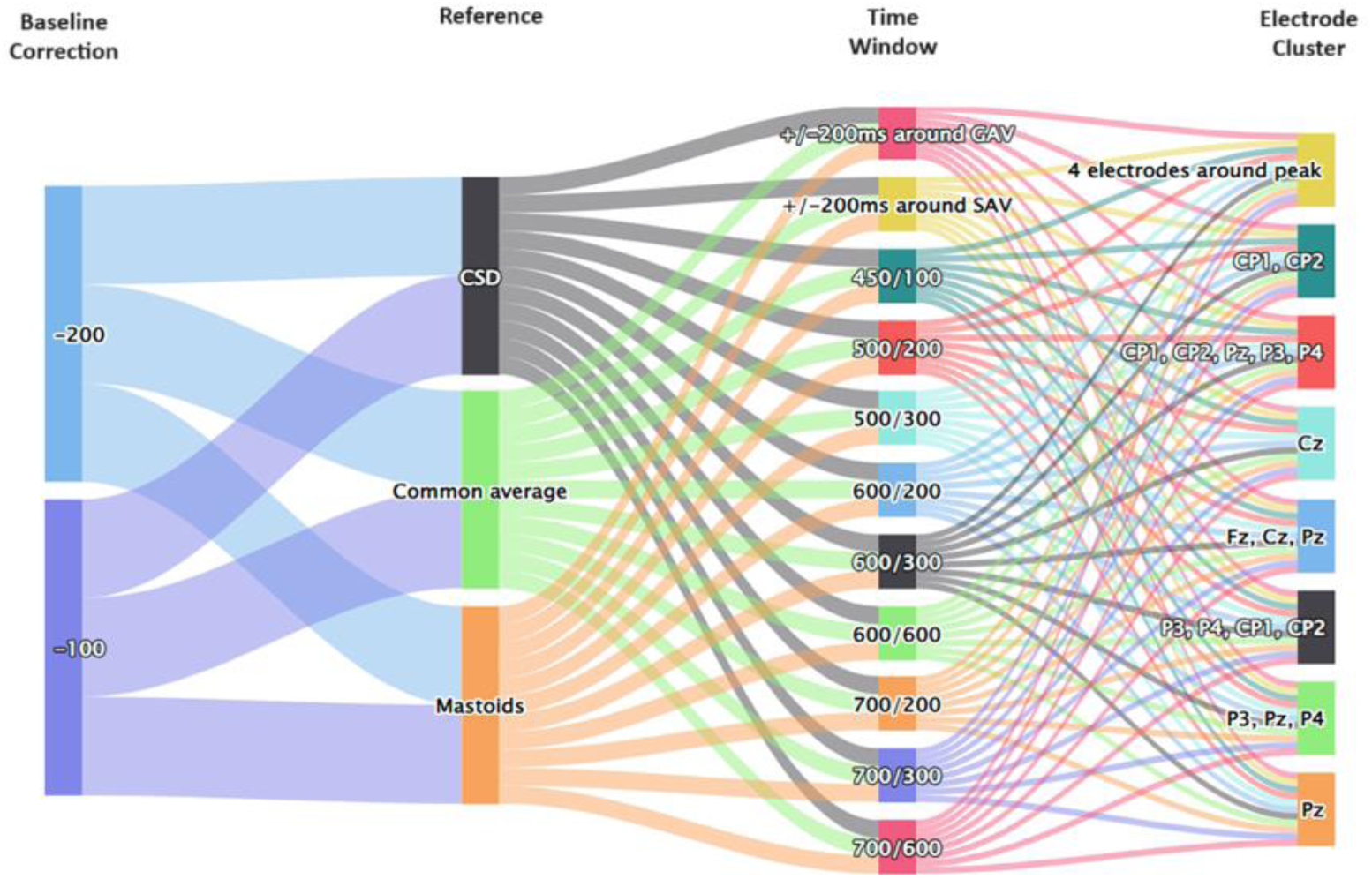
Sankey diagram of the four decision nodes and alternative options that created the 528 pipelines of the LPP quantification multiverse use case. Each option at each decision node was computed in combination with all other options at all other decision nodes, creating a fully factorial multiverse analysis across options. CSD = current source density, GAV = grand average peak, SAV = subject average peak. Time windows = the centre point of the window in post-stimulus ms / the width of the window in ms.

The two baseline correction options ((1) −100 ms to 0 ms and (2) −200 ms to 0 ms)) were applied to the preprocessed data. Bad epochs were identified following FASTER (Nolan et al., 2010), using a *z*-score threshold of ±3 on three EEG properties: deviation from channel means, epoch variance, and maximum amplitude difference. Epochs exceeding this threshold in any property were removed from further processing. Participants were excluded at this point if less than 25% of all trials for any condition remained in the data. The data were re-referenced to each of the three offline reference alternatives (linked mastoids, common average reference (CAV), and current source density (CSD). The CSD reference was applied using spherical spline surface Laplacian method (Perrin et al., 1989) with default Legendre polynomial order (20) and default smoothing parameter (1e-5).

The individual average peak with channel index and the grand average peak, which are required to define two of the alternative time window parameters later in the procedure, were stored for each of the six datasets per participant per condition. The midline peak, defined as the midline electrode with the maximum amplitude between 400 and 600 ms of the grand average, was identified and stored to define the electrode cluster of four electrodes around the peak. This was identified as channel Pz, therefore the four electrodes around the peak were CP1, CP2, P3, P4. The 11 time window options ((1) 400-500 ms, (2) 400-600 ms, (3) 350-650 ms, (4) 500-700 ms, (5) 450-750 ms, (6) 300-900 ms, (7) 600-800 ms, (8) 550-850 ms, (9) 400-1000 ms, (10) +/− 200ms around the grand average within 300-1000 ms, (11) +/− 200 ms around the single subject average within 300-1000 ms), and the eight electrode cluster options ((1) CP1, CP2, Pz, P3, P4; (2) P3, P4, CP1, CP2; (3) P3, Pz, P4; (4) Fz, Cz, Pz; (5) CP1, CP2; (6) Cz; (7) Pz; (8) four electrodes around the midline peak) were applied for LPP quantification. The mean LPP amplitude was saved across all quantification pipelines (528 pipelines illustrated in Figure 1), for each condition, for each participant.

### 2. Sampling Approaches

The Multiverse Sampling Tool implements three sampling approaches: random, stratified, and active learning. For each sample, model fit is calculated for each pipeline, and in the case of active learning, model fit is also predicted for unsampled pipelines.

#### Model Fit and Predicted Model Fit

The main criterion by which the sampling approaches are evaluated is the model fit obtained by each pipeline. Model fit refers to how well the applied regression model predicts individual differences in the outcome variable (extraversion score) using the LPP quantifications across six emotion categories as predictors. Fit is measured by *R^2^*, where a higher *R^2^* indicates a better fit. This criterion allows a model-specific evaluation of the sampled pipelines, which is beneficial for reporting uncertainty and achieving clear interpretability of the pipeline outcomes. Using model fit to compare pipeline samples is particularly relevant in EEG multiverse analyses, where data processing decisions can greatly influence the amount of noise and signal of interest remaining in the data for subsequent statistical modelling (e.g., Robbins et al., 2020; Rodrigues et al., 2024; Winkler et al., 2015). This criterion emphasises how different LPP quantification options affect model performance in estimating brain-personality relationships.

Importantly, in our multiverse analysis use case, the *R^2^*is used as an estimate of model fit in two different ways. First, for each sampling approach (random, stratified, and active learning), the *R^2^* reflects model fit as a measure of the proportion of variance in extraversion explained by LPP amplitudes to emotional stimuli across six conditions (see above) for each pipeline in the sample. In addition, unlike the other sampling approaches, the active learning sampling approach is able to estimate the prediction accuracy of the remaining pipelines that were not sampled. In this additional case, the *R^2^* is used to represent the predicted model fit of the unsampled pipelines trained on the directly sampled pipelines.

#### Random Sampling

The random sampling procedure selects a subset of pipelines of a specified sample size without replacement, where each pipeline has an equal probability of being selected. The pipeline configurations and the model fit following each sampled pipeline are stored, but these do not influence the sampling procedure. For example, if a sample size of 26 is specified within the sampling script, as in our use case described below, the script will select the random sample as:

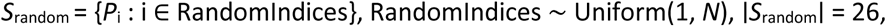

where *S*_random_ is the set of pipelines selected via random sampling; RandomIndices is a random selection of 26 indices, chosen uniformly from all possible pipelines; *N* is the total number of pipelines in the multiverse; and ∣*S*_random_∣ = 26 ensures that the random sample of pipelines contains exactly 26 pipelines.

#### Stratified Sampling

The stratified sampling procedure ensures a balanced representation of the options at each decision node. In our use case, there are 2 x 3 x 11 x 8 pipelines in the full multiverse (2 baselines, 3 references, 11 time windows, 8 electrode clusters). The function iterates over each decision node (category) to sample pipelines from within it. For each decision node, it groups the pipelines by the options within it, excluding previously selected pipelines, and determines how many pipelines to take from each group within the decision node for a balanced representation without exceeding the sample size. This means that the stratified sample will, as much as possible for the sample size, represent the diversity of the options within each decision node of the multiverse. This will return a sample that is balanced in the options across each decision node. For example, −100 ms and −200 ms baseline durations will feature equally in the sample. The pipeline configurations and model fit are stored, but the model fit does not influence the sampling procedure.

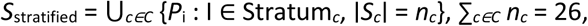

where *S*_stratified_ is the set of pipelines selected using stratified sampling; ⋃*_c∈C_* is the union of subsets (S*_c_*) sampled from each category (strata) *c* in *C; C* is the set of categories (strata); Stratum*_c_* is the subset of pipelines belonging to category *c* (a stratum); ∣*S_c_*∣ = *n_c_* is the number of pipelines sampled from stratum *c*; and ∑*_c∈C_ n_c_* = 26 is the total number of pipelines sampled across all strata.

#### Active Learning Sampling

The active learning procedure comprises two stages: low dimensional embedding using a subset of observations from the sample of participants, and active learning-based sampling of pipelines from the low dimensional space.

##### Low dimensional embedding

The first step in the active learning sampling procedure was to generate a low-dimensional embedding by projecting the high-dimensional data into two dimensions using a subsample of participants. This was performed using MATLAB, following the multiverse computation in this case, and prior to running the Multiverse Sampling Tool. The MATLAB scripts used for embedding and visualising the two-dimensional space are available [here](https://github.com/cassiesh/MultiverseSamplingTool/tree/main/UseCaseMultiverse_MATLABScripts/StdDevImages). For users preferring to compute the embedding using Python, an example is provided by Dafflon et al. (2022). Importantly, dimensional embedding must be performed prior to using the active learning sampling approach in the Multiverse Sampling Tool, as it helps to reduce overfitting to the training set while preserving the discriminative characteristics between pipelines (Dafflon et al., 2022).

In line with the statistical model of our multiverse analysis use case, the 528 pipelines were embedded into a two-dimensional space with respect to the Euclidean distances of pairwise LPP amplitudes across the vector of emotion conditions. The choice of similarity measure may depend on the statistical model. While Dafflon et al. (2022) used cosine similarity, Euclidean distance was more appropriate in our use case because the magnitude of the condition vectors was relevant for our subsequent regression model. The embedding process began by calculating a Euclidean distance matrix between all unique participant pairs within the training set of 20 participants. This produced a two-dimensional matrix of pairwise distances of size 190 (unique participant pairs) x 528 (pipelines).

Next, the low dimensional space was generated by applying *t*-distributed stochastic neighbour embedding (*t*-SNE; Van der Maaten & Hinton, 2008) with a perplexity value of 5 to this matrix. The *t*-SNE dimensionality reduction algorithm converts the Euclidean distances of between-person pairwise LPP amplitude difference vectors (condition minus neutral) into joint probabilities. It then minimises the Kullback-Leibler divergence between the joint probabilities to accurately represent them in two dimensions. The process maintains relative distances, such that similar pipelines in terms of pairwise similarity across the vector of experimental conditions are positioned in close proximity in the space, while dissimilar pipelines are positioned further apart, revealing underlying patterns of individual differences in the EEG pipeline multiverse. The low perplexity parameter of 5 prioritises local over global structure, emphasising the distances between closely clustered pipelines, potentially yielding a detailed insight into the small-scale variations. Among several explored dimension reduction algorithms (results in Supplementary 4), *t*-SNE with a perplexity of 5 produced the most even distribution of pipelines across the space, which is optimal for active learning to efficiently explore the space. The distribution of pipelines across the space is illustrated in Figure 2 in the results section.

**Figure 2.**
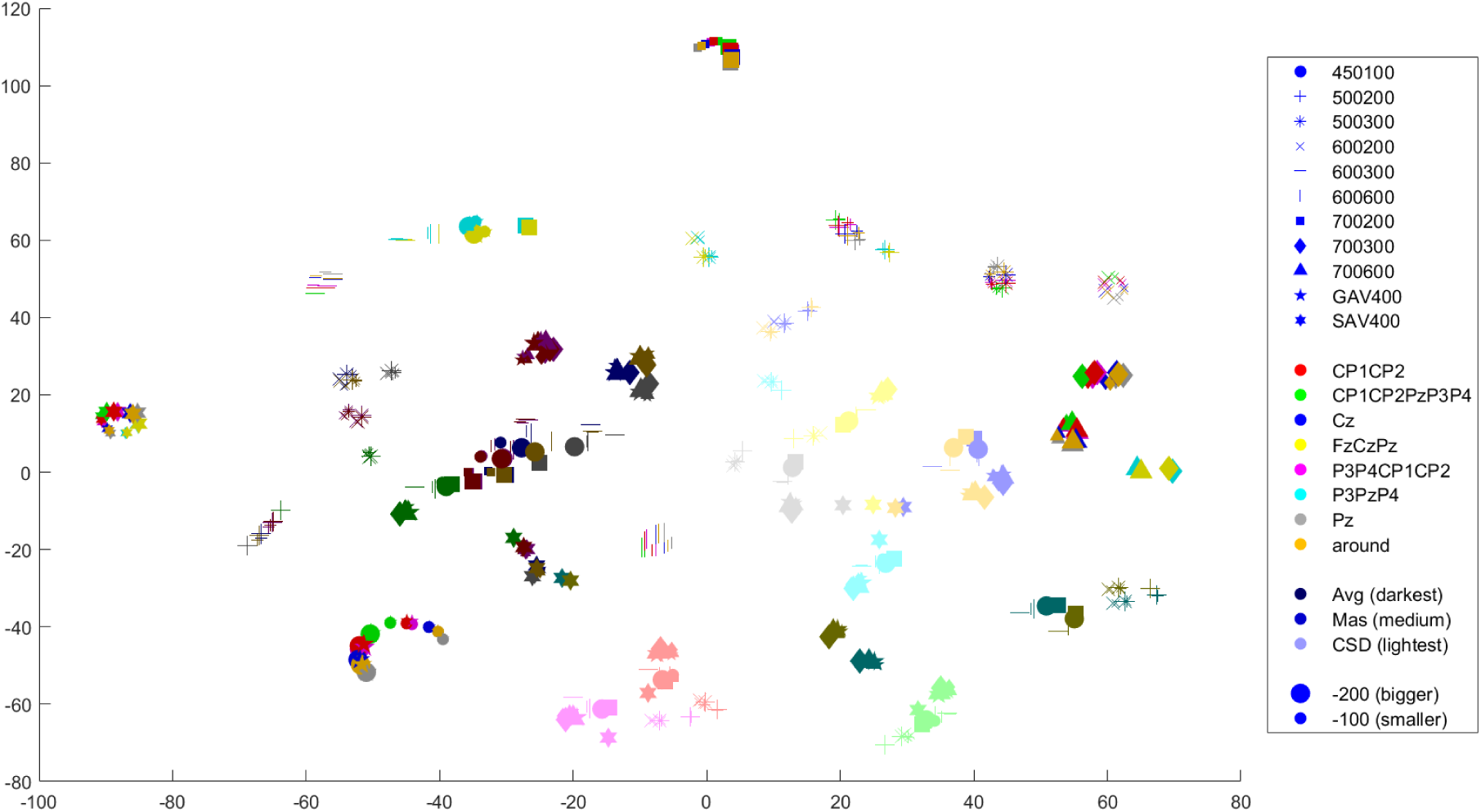
The distribution of all 528 pipelines with respect to pairwise Euclidean distances in the LPP difference scores across the condition vector, in the two-dimensional space after embedding with *t*-SNE. *N* = 20 participants.

##### Active learning

The active learning-based sampling procedure uses Bayesian optimisation to iteratively sample pipeline configurations with the aim of maximising predictive accuracy. The process begins by taking a ‘burn-in’ sample, which is a random sample of *n* pipelines. This burn-in sample is used to fit a Gaussian Process (GP) regression model, which provides probabilistic predictions of predictive accuracy across the low dimensional space. Once the burn-in sample is taken, the specified sample of pipelines is iteratively selected using Bayesian optimisation. At each iteration, an acquisition function (upper confidence bound) identifies the next point in the low dimensional space based on a user-defined balance (setting the *k* parameter) between exploration (sampling uncertain areas) and exploitation (sampling areas predicted to have high accuracy). The pipeline closest to the selected point in the space is evaluated and its accuracy is used to update the GP model. This procedure continues until the sample size is reached. Finally, the information from the sample is used to estimate the full multiverse of results across all defensible pipelines. The *R^2^* values stored for the 26 sampled pipelines represent actual model fit, whereas the *R^2^* values for the unsampled pipelines represent predicted model fit based on the GP model. The pipeline configurations do not influence the sampling procedure, whereas model fit does.

The selection of the next sampled pipelines is based on the Upper Confidence Bound (UCB) algorithm:

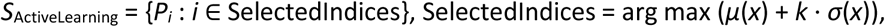

where *S*_ActiveLearning_ is the set of pipelines selected using active learning; *P_i_* is a pipeline in the multiverse of pipelines; SelectedIndices are indices of the pipelines selected iteratively based on the utility function; the arg max operation selects the index of *i* of the pipeline *P_i_* that maximises the utility function; *μ(x)* is the mean prediction from the GP model for pipeline *P_i_* at location *x* in the low-dimensional space; *σ(x)* is the uncertainty (standard deviation) of the GP model prediction at location *x; and k* is the parameter that controls the balance between exploration (*σ*(*x*)) and exploitation (*μ*(*x*)).

### 3. The Multiverse Sampling Tool

In this section, we outline how the sampling procedures are implemented for our multiverse analysis use case, and highlight elements that can be modified.

#### Sampling Methods for the Multiverse Analysis Use Case

The number of pipelines sampled by the three sampling approaches can be adjusted within the script. In our use case, various sample sizes were set based on small proportions of the full 528 pipeline multiverse. As previous work has demonstrated that ERP multiverse analyses can comprise quadrillions of pipelines (e.g., Paul, Short et al., 2022), the expectation that small proportions may be required in large EEG multiverse studies is plausible. Specifically, we sampled 26 pipelines (5%), 53 pipelines (10%), and 79 pipelines (15%). The results of 26 pipeline samples are reported within the main text, and the 53 and 79 pipeline samples are reported in Supplementary 5.

#### Sample Evaluation

The script provides several tabular and graphical outputs that report the representativeness of each sample with respect to the full multiverse and the model of interest. The outputs include (1) the median *R^2^*-values (model fit of each pipeline in each sample and the full multiverse, and predicted model fit of the estimated full multiverse in the case of active learning sample), (2) the Kolmogorov-Smirnov statistic of the distribution of *R^2^* between each sample and the full multiverse, (3) the pipelines in each sample and the full multiverse that yielded the highest *R^2^*-values, (4) a raincloud plot of *R^2^*-values for each sample and the full multiverse, (5) a scatter plot of the spatial distribution of sampled pipelines across the low dimensional space, and (6) specification curve plots of the *R^2^*-values of each sample and the full multiverse.

In our use case, the model of interest is a linear regression model that predicts extraversion scores using the vector of LPP amplitude differences across the six emotional conditions minus the neutral condition. The model fit is calculated for each pipeline in each sample by fitting the specified linear regression model and comparing the model predictions with the observed outcomes using the *R^2^*-metric. More generally, the metric can be modified according to the user’s needs to any appropriate and quantifiable measure of fit or prediction quality indices, such as *R^2^*, mean squared error, or otherwise. Therefore, the script is adaptable for a variety of brain-personality, -behaviour or -cognition relationship hypotheses.

#### Structure and Modifiability

The sampling script takes several data files as input: (1) a data frame containing the LPP values of interest across the full multiverse of pipelines (which can be computed on a sample of participants), where each pipeline for each participant for each condition is represented by a row, and the participant ID, condition label, pipeline label, and ERP value are organised across columns; (2) a dimensionally reduced format of these results from which a training subset of the participant data will be obtained; and (3) data frame(s) containing other relevant variables, such as behavioural or questionnaire data to be predicted by the brain data (here LPP). Participant IDs must match across all imported data files. While no one-size-fits-all rule exists for determining the optimal sample size for this purpose, the subsample should be large enough to capture individual differences and reduce overfitting relative to the dimensionality of the multiverse, yet small enough to remain computationally feasible. The subsample can be selected to preserve generalisability, for example, through random selection while avoiding strategies such as maximising variance in the outcome variable, which could bias the multiverse representation of model performance. In our use case, we selected the first 20 participants after data preprocessing (∼20% of the full sample), independent of extraversion scores, and thus we expect the subsample to reasonably generalise to the broader dataset.

Modifying and extending the active learning sampling script by Daflon et al. (2022), the Multiverse Sampling Tool is split into two components: Main.py, which executes the functions and commands, and Helper.py, which defines the functions used within Main.py. Both scripts include adjustable parameters that can be tailored to suit the specific characteristics of the imported data. A detailed overview of the structure and modifiability of each section of the Main.py and Helper.py scripts is provided in Supplementary 1 and Supplementary 2, respectively. We recommend sharing the final scripts, including all parameter settings and customisations, alongside the reported multiverse analysis to increase transparency of the multiverse sampling procedure.

## Tool Preparation and Results

This section describes the outputs generated by the Multiverse Sampling Tool. Rather than focusing on the substantive individual differences research question, we focus specifically on comparing the different multiverse pipeline sampling approaches, and the effectiveness of the tool to facilitate this comparison.

### Preliminary Data Preparation

#### Visualisation of the waveforms across the ERP multiverse

Visualising variation in waveforms across decision node options is a useful resource for interpreting EEG multiverse analysis results. In our use case, the grand average signal waveform for each condition was visualised across each combination of reference scheme and electrode cluster (Supplementary 3, using the MATLAB code available [here](https://github.com/cassiesh/MultiverseSamplingTool/tree/main/UseCaseMultiverse_MATLABScripts/StdDevImages)). To ensure computational feasibility, such visualisations can be limited to a subset of decision nodes or performed on a subset of participants. Visualising signal waveforms across a sample of decision nodes, such as in Supplementary 3, helps to identify how alternative processing options may impact waveform morphology. This can be informative for later interpretation of multiverse analysis results, as morphological variation may influence the strength and direction of the brain-personality associations of interest. Additionally, such visualisations can inform the tuning of modifiable parameters within the sampling tool. For example, if the visualisation reveals that options at a particular decision node lead to greater variation in waveform morphology between conditions than options at other decision nodes, users may wish to allocate more sampling resources toward capturing this variance to ensure it is not overlooked in small sample sizes. If such a data-driven targeted sampling is applied, both the accompanying visualisation and the sampling modifications should be reported. The sampling tool facilitates this transparency.

For example, Supplementary 3 shows that variability introduced by different reference schemes, particularly the CSD reference, appears more pronounced across electrode clusters than the variability introduced by electrode cluster selection within a given reference scheme. The CSD reference yields greater condition-specific waveform divergence and more variability in LPP onset latencies compared to the CAV and linked mastoids references, which may have implications for the selection of the time window of interest. Furthermore, within the CAV and linked mastoids reference schemes, the LPP in response to emotional conditions appears relatively stable across electrode clusters, whereas the neutral condition shows more pronounced variability under the CSD reference. This pattern highlights how regression coefficients in our multivariate model may be contingent on pipeline selection.

#### Low dimensional embedding

The *t*-SNE dimensionality reduction algorithm from the Statistics and Machine Learning Toolbox in MATLAB (R2022b Mathworks, 2022) was selected, as explained in the method section, and applied to the training set of participants (*N* = 20). The distribution of pipelines across the low dimensional space is visualised in Figure 2.

This visual representation highlights the decision nodes that yield more or less divergent LPP patterns between participants, relative to the conditions within the experimental design and the predictive model. These insights can guide parameter tuning within the sampling tool. For instance, the visualised distribution of the pipelines across the space can inform the choice between a more exploratory or exploitative strategy for the active learning sampling approach. Densely clustered regions suggest that a more exploratory approach (i.e., a large *k* parameter) is needed to avoid overfitting to localised clusters, whereas a more uniform pipeline distribution may be appropriate for a more exploitative approach (i.e., a small *k* parameter), depending on the study’s objectives. Additionally, the visualisation can help to determine an optimal burn-in sample size for the active learning sampling. Too few burn-in samples may limit the model’s representativeness, while too many may inefficiently expend computational resources. In cases where pipelines form dense clusters, a relatively larger random burn-in sample can help to provide more comprehensive coverage. In the use case (Figure 2), the between-person LPP amplitude condition vector appears particularly sensitive to the reference scheme options, corroborating patterns observed in the waveform plots (Supplementary 3). Given the dispersed distribution of pipelines across the space, we selected a modest random burn-in sample size of 10 pipelines and an exploratory *k* parameter of 10 for this multiverse.

### Data Imports into the Sampling Tool

Four data files are imported into the tool: (1) the LPP mean amplitude values across all participants, conditions and pipelines (4 columns for participant ID, condition, pipeline combination and LPP mean amplitude, and 241,472 rows of each participant x condition x pipeline combination), as a .mat file; (2) the difference scores (emotional condition minus the neutral condition) of the LPP mean amplitude across all conditions and pipelines (a 3 dimensional array of 98 subject IDs x 6 conditions x 528 pipelines), as a .mat file; (3) the low dimensional embedded data matrix of pairwise Euclidean distances for the first 20 participants (training sample) (528 pipelines x 2 participant IDs), as a .mat file; and (4) the extraversion scores of all 98 participants, identifiable by the ID, as a .dat file. Notably, the file format for running the tool is flexible. The tool will read the data as dataframes.

### Results Output

We describe the outputs generated by the Multiverse Sampling Tool, using our use case to illustrate their purpose and interpretability. In our multiverse analysis use case, we compute the full multiverse for the full sample of participants, to strengthen the benchmark comparison of the samples we provide for future multiverse analyses.

#### Output 1: Median Predictive Accuracy and Kolmogorov-Smirnov Statistic

The first output is a table summarising (a) the median *R^2^*-values of the pipelines sampled by each approach (random, stratified, active learning) and, for the present use case, the full multiverse, and (b) the Kolmogorov-Smirnov (K-S) statistic comparing the distribution of *R^2^*-values from each sample to the full multiverse. For all approaches, the *R^2^*-values of the 26 sampled pipelines represent model fit, indicating how well the model explains the variance in the outcome variable. In contrast, the active learning approach also predicts *the R^2^*-values for unsampled pipelines (predicted model fit), estimating generalisation across the unsampled multiverse. It should be considered when interpreting the results that the active learning approach is the only approach of the three that provides an extrapolated distribution beyond its sampled pipelines. Benchmarking active learning against random and stratified sampling approaches quantifies the added value gained by the additional computational cost.

##### Usefulness

The K-S statistic serves as a robust measure of distributional alignment in terms of model performance, quantifying the maximum distance between the cumulative distribution functions of the *R^2^*-values obtained by each sampling approach and the full multiverse. Lower K-S values indicate closer alignment and suggest that the sampled pipelines approximate the central tendency and variability of the full multiverse. This is crucial for generalising conclusions beyond the sampled pipelines to the full multiverse. When full multiverse computation is infeasible, the K-S statistic between samples remains informative. For instance, comparing the distributional alignment of the samples to one another can inform whether conclusions drawn from one sample are likely to generalise to others, providing information on the sampling uncertainty. However, the active learning sample will have the same number of data points/estimates (here *R*^2^) as the full multiverse, whereas the random and stratified samples will be limited to the specified sample size. This denser distribution may better approximate the structure of the full multiverse distribution, while sparser samples may miss key features.

##### Interpretation

The K-S statistic reveals that the active learning sample most closely approximates the distribution of *R*^2^-values from the full multiverse. This aligns with the exploratory parameter settings of the active learning approach, which specifically aims to capture the statistical properties of the full multiverse. The active learning approach has most effectively estimated the full multiverse of model fit based on the 26 sampled pipelines, suggesting that the 26 sampled pipelines effectively generalised to the broader multiverse. Therefore, active learning emerges as the optimal sampling approach for approximating the underlying central tendency of the full multiverse in terms of model fit. The sample with the smallest K-S statistic was the random sample at a larger sample size of 10% of the full multiverse, and the stratified sample at a sample size of 15% of the full multiverse (Supplementary 5). However, the K-S statistics at sample sizes of 10% and 15% did not exceed 0.154, generally showing a trend of closer approximation as sample size grows.

#### Output 2: Predictive Accuracy of the Pipelines

The second output provides a table listing the pipelines with the highest *R^2^*-values in the full multiverse and in each of the three sampling approaches. This does not imply a hierarchy of validity between pipelines or preprocessing decisions. Rather it provides insight into which combinations of options across decision nodes lead to LPP values with greater explanatory power for the specified outcome variable. However, this output alone does not confirm robustness, as high *R^2^*-values could be due to chance, especially without external validation. To address this, the Multiverse Sampling Tool includes a built-in option for a ‘lock box’ validation set, allowing users to cross-validate high-performing pipelines across independent subsamples. Additionally, it is important to recognise that in multivariate regression models, pipelines may exhibit similar predictive performance even if individual predictors vary in importance or change signs across pipelines. This complicates interpretation compared to simpler univariate models. Nevertheless, multivariate models are widely used in the literature. In such cases, users who wish to explore predictor-level contributions to the model fit are encouraged to examine the regression coefficients and the waveform visualisations exemplified in Supplementary 3 when interpreting the results of this output.

##### Usefulness

While a high *R^2^* does not inherently equate to superior data quality, it indicates which pipelines and, by extension, which sampling methods, capture data features that enhance the prediction accuracy of the model for the specified outcome. Therefore, while not synonymous with superior data quality, this output serves as a useful exploratory tool for identifying those pipelines.

##### Interpretation

Table 2 reveals a pattern where pipelines using the linked mastoids reference scheme, time window based on the subject average peak, and electrode clusters including CP1 and CP2 are overrepresented within the ten pipelines that yielded the highest model fit across the full multiverse. This suggests that these LPP quantification options ***may*** be relatively more effective at isolating the relevant LPP components that predict extraversion. Only the random sample included any of the ten pipelines with the highest *R^2^*-value from the full multiverse, the active learning sample with an exploratory acquisition function showed the highest average *R^2^*-values across the top ten, and the stratified sample, which aims to represent all preprocessing options individually, showed a wider range of *R^2^*-values. These findings highlight the influence that the chosen sampling strategy can have on the conclusions drawn from a sampled multiverse analysis. When the sample size was 10% and 15% of the full multiverse, the active learning did not sample any of the ten pipelines with the highest *R^2^*, but remained the sample with the greater average *R^2^* across the ten pipelines with the highest value (Supplementary 5). The random sample sampled one of these pipelines at 10% sample size, but none at 15% sample size, highlighting a lack of reliability from the random sampling approach. The stratified sample sampled three of these pipelines at 10% sample size, and one at 15% sample size, suggesting that, when the active learning acquisition function is biased towards exploration over exploitation, the stratified sample seems most likely of the three approaches to include within the sample pipelines with the highest *R^2^*-values.

**Table 1.** The median *R*^2^ of the model across the pipelines included in each sample and the full multiverse, and the K-S statistic of the *R^2^* between each sample and that of the full multiverse. (Sample size = 26 pipelines, 5% of the multiverse).

| Sampling Approach | Median $R^2$ | K-S Statistic |
| --- | --- | --- |
| Full multiverse | 0.064 | - |
| Active learning | 0.065 | 0.150 |
| Random | 0.057 | 0.158 |
| Stratified | 0.052 | 0.193 |

**Table 2.** The ten pipelines with the highest *R^2^* from the full multiverse and each of the sampling approaches. Each result includes the pipeline identifier based on the combination of preprocessing options selected across decision nodes and the corresponding *R^2^*-value for that pipeline, categorised by sample. (Sample size = 26 pipelines, 5% of the full multiverse).

| Best Full Sample | $R^2$ | Best Active Learning | $R^2$ | Best Random Sampling | $R^2$ | Best Stratified Sampling | $R^2$ |
| --- | --- | --- | --- | --- | --- | --- | --- |
| **b-200 ,rMas, 500200, CP1CP2** | **0.20**<br>**3** | b-100, rCSD, 500200, P3P4CP1CP2 | 0.163 | **b-100, rMas, 600200, CP1CP2** | **0.16**<br>**9** | b-200, rMas, SAV400, CP1CP2 | 0.147 |
| **b-100, rMas, 500200, CP1CP2** | **0.18**<br>**8** | b-100, rCSD, 500200, CP1CP2 | 0.163 | b-200, rAvg, 500200, CP1CP2 | 0.15<br>7 | b-200, rAvg, 700300, CP1CP2PzP3P4 | 0.137 |
| **b-200, rMas, SAV400, CP1CP2PzP3P4** | **0.18**<br>**4** | b-100, rCSD, 500200, Cz | 0.160 | b-100, rAvg, 600200, CP1CP2PzP3P4 | 0.11<br>5 | b-100, rCSD, 500200, FzCzPz | 0.121 |
| **b-100, rMas, SAV400, CP1CP2PzP3P4** | **0.18**<br>**2** | b-100, rCSD, 500200, CP1CP2PzP3P4 | 0.159 | b-200, rMas, 450100, CP1CP2PzP3P4, | 0.09<br>5 | b-100, rMas, 450100, CP1CP2 | 0.120 |
| **b-200, rAvg, 500200, CP1CP2PzP3P4** | **0.17**<br>**7** | b-200, rCSD, 500200, CP1CP2 | 0.159 | b-200, rCSD, 500200, FzCzPz | 0.09<br>1 | b-200, rMas, 450100, P3P4CP1CP2 | 0.103 |
| **b-200, rMas, 700200, CP1CP2** | **0.17**<br>**3** | b-200, rCSD, 500200, CP1CP2PzP3P4 | 0.158 | b-200, rAvg, 700300, P3P4CP1CP2 | 0.09<br>1 | b-200, rMas, GAV400, CP1CP2 | 0.086 |
| **b-200, rMas, 600200, CP1CP2** | **0.17**<br>**3** | b-200, rCSD, 500200, P3P4CP1CP2 | 0.158 | b-200, rAvg, 600600, P3P4CP1CP2 | 0.08<br>3 | b-200, rAvg, SAV400, P3PzP4 | 0.082 |
| **b-200, rAvg, SAV400, CP1CP2PzP3P4** | **0.17**<br>**1** | b-100, rCSD, 500200, around | 0.156 | b-100, rCSD, 600200, Cz | 0.08<br>1 | b-100, rMas, 700300, around | 0.080 |
| **b-100, rMas, 600200, CP1CP2** | **0.16**<br>**9** | b-200, rCSD, 500200, Cz | 0.151 | b-200, rAvg, SAV400, around | 0.06<br>8 | b-100, rMas, 600200, around | 0.072 |
| **b-100, rAvg, SAV400, CP1CP2PzP3P4** | **0.16**<br>**8** | b-200, rCSD, 500200, around | 0.148 | b-200, rCSD, 600300, CP1CP2 | 0.06<br>3 | b-200, rCSD, 450100, around | 0.062 |

#### Output 3: Raincloud Plots of Predictive Accuracies

The third output is a series of raincloud plots displaying central tendency and variability of the *R^2^*-values of the pipelines sampled by each approach and the full multiverse.

##### Usefulness

The raincloud plots (Figure 3) provide a clear visual comparison of the central tendency and dispersion of the *R^2^*-values of the specific model of interest across each sampling approach and the full multiverse. This plot is especially useful for quickly identifying sampling approaches that may skew or under-represent variability in model fit, a crucial factor for interpreting multiverse sampling uncertainty. It is also clearly illustrated in this plot that the active learning approach uses the selected sample size to estimate the full multiverse of pipelines, in contrast to the random and stratified sampling approaches.

**Figure 3.**
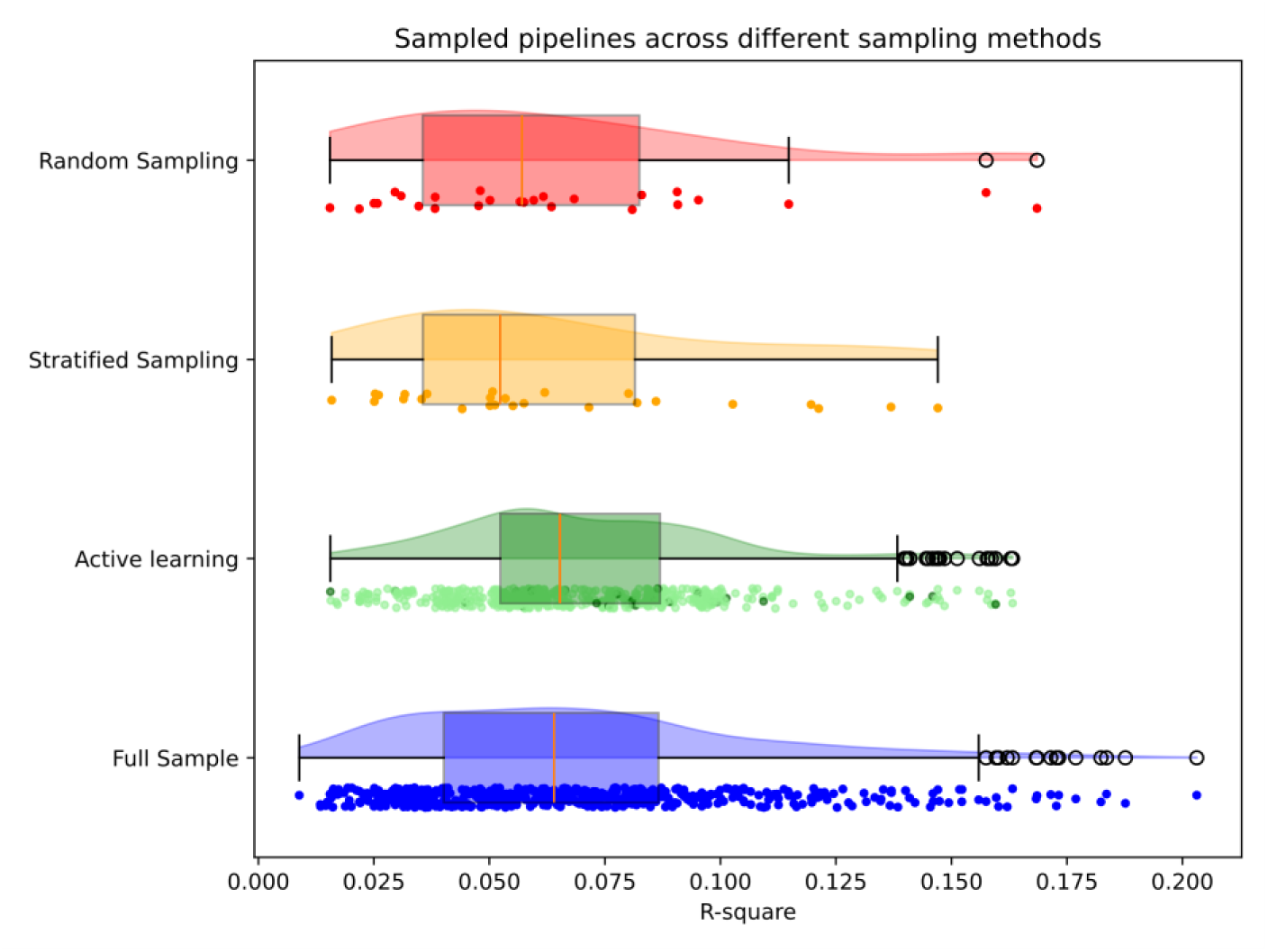
Raincloud plot of *R^2^*-values of the pipelines sampled by each of the three sampling approaches and the full multiverse. The pipelines directly sampled by the active learning algorithm are in dark green, whereas the pipelines estimated are in light green (sample size = 26 pipelines, 5% of the multiverse)

##### Interpretation

The broad dispersion of *R^2^* captured by the stratified sample relative to the other samples suggests that it may provide the most accurate report of variability in *R^2^* across the entire multiverse of pipelines. However, the active learning sample most accurately represents the median of the full multiverse. Ensuring that the sample median aligns with the full multiverse median is critical, particularly when the goal is to interpret the median model fit as a key inferential metric (e.g., Simonsohn et al., 2020). Such alignment reduces the risk of drawing biased conclusions from a multiverse analysis based on a non-representative sample. Therefore, based on this criterion, the active learning sample appears to introduce the least bias in terms of capturing the central trend of model fit across the multiverse. We recommend that Figure 3 be shared in a supplementary file alongside the reported sampled multiverse analysis for increased transparency of multiverse sampling uncertainty. At the 10% and 15% sample sizes, the stratified sample consistently presented the largest range of *R^2^*-values, while the median *R^2^* of the active learning sample consistently represented that of the full multiverse mostly closely (Supplementary 5).

#### Output 4: Scatter Plots of Spatial Distribution in the Low Dimensional Space

The fourth output is a set of scatter plots that map the spatial distribution of the pipelines sampled by each approach within the two-dimensional embedding of the full multiverse. This embedding is based on the Euclidean distances of pairwise LPP amplitudes across the vector of experimental conditions (emotion minus neutral). Each panel corresponds to one sampling approach. This provides a visual comparison of the spatial representativeness of each sample to the full multiverse. Additionally, the tool calculates the average nearest neighbour distance between each sample and the full multiverse. Lower distance values indicate a closer match to the full multiverse distribution.

##### Usefulness

The table quantifies and the scatter plots visualise the spatial representativeness achieved by each sampling approach in terms of the pairwise dissimilarity of the LPP vector across the low dimensional space. This is especially important for individual differences analyses, where sampling pipelines that span the full space of between-person variability is crucial for generalisability. A sample that is well-dispersed across this space indicates that the variability in the between-person LPP vector dissimilarity across the multiverse is captured. Conversely, if pipelines are clustered in certain regions, while large areas of the space are unsampled, key aspects of variability may be missed, potentially biasing conclusions toward those regions that were sampled. This table and set of scatterplots also allow users to evaluate how their choice of active learning parameters, such as the *k* parameter, may have affected the sample dispersion, and could thereby support data-driven refinement of sampling strategies based on representation of individual differences.

##### Interpretation

The scatter plots show that each sample was dispersed across the low dimensional space of pairwise Euclidean distances of LPP vectors across conditions. This suggests that each approach was effective in sampling pipelines that broadly represent the multiverse in terms of individual differences. This interpretation is supported by the average nearest neighbour distances presented in Table 3, which show relatively similar values across the three sampling methods. The stratified sample yielded the lowest average distance to the full multiverse, indicating slightly better spatial representativeness compared to the random and active learning samples. This suggests that stratified sampling may be particularly useful for studies prioritising diversity in preprocessing choices and between-person variability. Regarding the active learning sample, the combination of an exploratory *k* parameter and ten burn-in pipelines appears to have been effective in preventing clustering and promoting dispersion, however this may prompt further tuning of the algorithm to explore the potential for a lower nearest neighbour distance. At a sample size of 10%, the stratified sample presented a slightly larger distance than the random and active learning samples, while at a 15% sample size, the random sample presented a slightly smaller distance than the stratified an active learning samples (Supplementary 5). However, the range of the distances across all sampling approaches and all sample sizes was fairly small at 66.47 - 72.53, with the stratified sample at the 5% sample size producing the smallest distance, and the stratified sample at the 10% sample size producing the largest distance.

**Table 3.** The average nearest neighbour distance between each sampling method and the full multiverse.

| Sample Size | Full Multiverse vs. Random<br>Sample | Full Multiverse vs.<br>Stratified Sample | Full Multiverse vs. Active<br>Learning Sample |
| --- | --- | --- | --- |
| 26 pipelines | 70.06 | 66.47 | 70.80 |

#### Output 5: Specification Curves

The final output is a set of specification curves (Simonsohn et al., 2020), one for each sample and one for the full multiverse. Each specification curve plots the *R^2^*-values of pipelines within the sample in vertical alignment with the options at each decision node that were applied to the data for the respective pipeline.

##### Usefulness

The specification curves visualise how the *R^2^*-values vary across the preprocessing options, helping to identify whether particular options or combinations of options lead to systematically higher or lower *R^2^*-values. As such, these figures provide a summary of the robustness of model fit to specific analytical decisions that is captured by each sample, to improve transparency of multiverse sampling uncertainty. However, it is imperative to carefully distinguish between a probabilistic and a possibilistic interpretation of specification curves if a principled multiverse of equivalent pipelines was defined (Del Guidice & Gangestad, 2021). A probabilistic interpretation considers each result in terms of its likelihood, and is deemed inappropriate for a principled multiverse, while a possibilistic interpretation considers each result as a possibility in the data, as all results across the multiverse hold theorised equivalence in the validity or precision (Hall et al., 2022; Sarma et al., 2024). Therefore, in addition to highlighting potentially influential options and combinations of options, specification curves at the sample comparison stage also serve as a visual aid for selecting samples that respect the diversity of valid outcomes across the multiverse.

##### Interpretation

The specification curves from our use case multiverse show largely consistent distributions of *R^2^*across the samples. All three samples reveal that pipelines using the 500/200, 500/300, 600/200, and SAV400 (400 ms around the subject average peak) time windows, and those using the CP1 and CP2 electrodes, tend to produce higher *R^2^*-values than other options at the same decision nodes. This higher *R^2^* for pipelines quantifying the LPP around 500 ms may reflect latency alignment with the psychometrically relevant LPP in this task paradigm, whereas the higher *R^2^* for pipelines quantifying the LPP around the subject average peak may indicate individual differences in psychometrically relevant peak latency. The higher *R^2^*for pipelines incorporating CP1 and CP2 electrodes suggests that these sites may capture meaningful inter-individual variance in emotion processing that is associated with trait extraversion. While all sampling methods detect these trends, the additional pipeline data points provided by the active learning communicates the trends more comprehensively. A notable observation is that the active learning sample appears to amplify the influence of the CAV to produce both high and low *R^2^*-values, a pattern that is less apparent in the full multiverse. This discrepancy could result from overfitting to a trend seen in the initial burn-in or 26 pipeline sample. This highlights the importance of tuning the burn-in sample size and *k* parameter. When the full multiverse cannot be computed for comparison, cross-validation of the trends visualised by the samples can help to validate them. Overall, all three samples visualise trends that are present in the full multiverse, supporting their utility for effectively identifying influential options in the full multiverse in terms of *R^2^*-values. While the active learning can communicate these trends more clearly, care should be taken to tune the algorithm appropriately. At the sample size of 10%, the overfitted pattern regarding the average reference was no longer apparent (Supplementary 5).

## Discussion

We report the first systematic comparison of random, stratified, and active learning sampling approaches in representing an EEG individual differences multiverse. Representation was compared in terms of the distribution of model fit values and spatial distribution across a low-dimensional space for samples of 26 pipelines (5% of the full multiverse). By focusing on model fit for the sampled pipelines and predicted model fit for the estimated pipelines as the metric of comparison, the script enables a data- and model-specific evaluation of multiverse sampling uncertainty. This is particularly valuable in EEG multiverse analyses, where different data processing decisions can influence the psychometric properties of the estimated EEG components, given the heterogeneity in the amount of signal and noise that remains in the data following variations in data preprocessing (e.g., Robbins et al., 2020; Rodrigues et al., 2024; Winkler et al., 2015). We found variation between samples in representativeness to the full multiverse, emphasising that pipeline sampling choices can influence conclusions drawn from sampled multiverse analyses. This underscores the need for transparent sampling procedures for large multiverse analyses, and for reporting multiverse sampling uncertainty when full multiverse computation is not feasible.

The Multiverse Sampling Tool and use case application assists future EEG multiverse analyses in reporting multiverse sampling uncertainty in two ways. First, when the computation of multiple pipeline samples using different sampling approaches is not feasible, this study provides a benchmark of sample representativeness and variation across sampling approaches for multiverse analyses of similar design. This benchmark can be used to inform the sampling approach and sample size selected to increase and inform the representativeness of the sampled multiverse to the full multiverse. Second, the open source Multiverse Sampling Tool can be used in future large-scale EEG multiverse analyses to improve the feasibility of this comparison. This tool can be used to compare the representativeness of different samples in an automated workflow, reducing the time and computational resources required. We discuss each of these contributions.

### Benchmark Comparison of Sample Representativeness

Our comparison of sampling approaches was based on a psychometrically (Eysenck, 1967; Speed et al., 2015) and empirically (e.g., Ku et al., 2020; Speed et al., 2015) relevant 528 pipeline LPP quantification multiverse, in which mean LPP amplitudes in response to stimuli across six emotional expression conditions were used to predict extraversion scores. Within these lines, the results of the sample comparisons may be relevant to multiverse analyses that focus specifically on ERP quantification, where the ERP of interest is temporally sustained and distributed across more than one electrode site, such as the LPP, and where a multiple linear regression model is fitted.

There was variability in how accurately each sample captured the central tendency of model fit across the full multiverse. The active learning sample most accurately captured both the median and overall distribution of model fit, while the stratified sample most accurately captured the range of model fit (Table 1 and Figure 3). Accurately capturing the central trend of model performance is a particularly relevant criterion given that median effect size has been proposed as a metric for inferential analysis of multiverse results (e.g., Simonsohn et al., 2020) and the median effect has been reported descriptively as part of EEG multiverse analysis results (e.g., Beauducel et al., 2024; Paul et al., 2024). This aligns with the exploratory parameter settings selected for the active learning approach in the present use case multiverse, which prioritised exploration of pipelines across the low dimensional space rather than focusing on pipelines with the highest model fit (Dafflon et al., 2022). That the median model fit of the stratified sampling approach, when all options were equally represented, did not match that of the full multiverse suggests that some decision nodes or preprocessing options may contribute disproportionately to the variability in model fit, skewing the median of the sample away from that of the full multiverse. This can be examined more closely in the specification curves (Figure 5).

**Figure 4.**
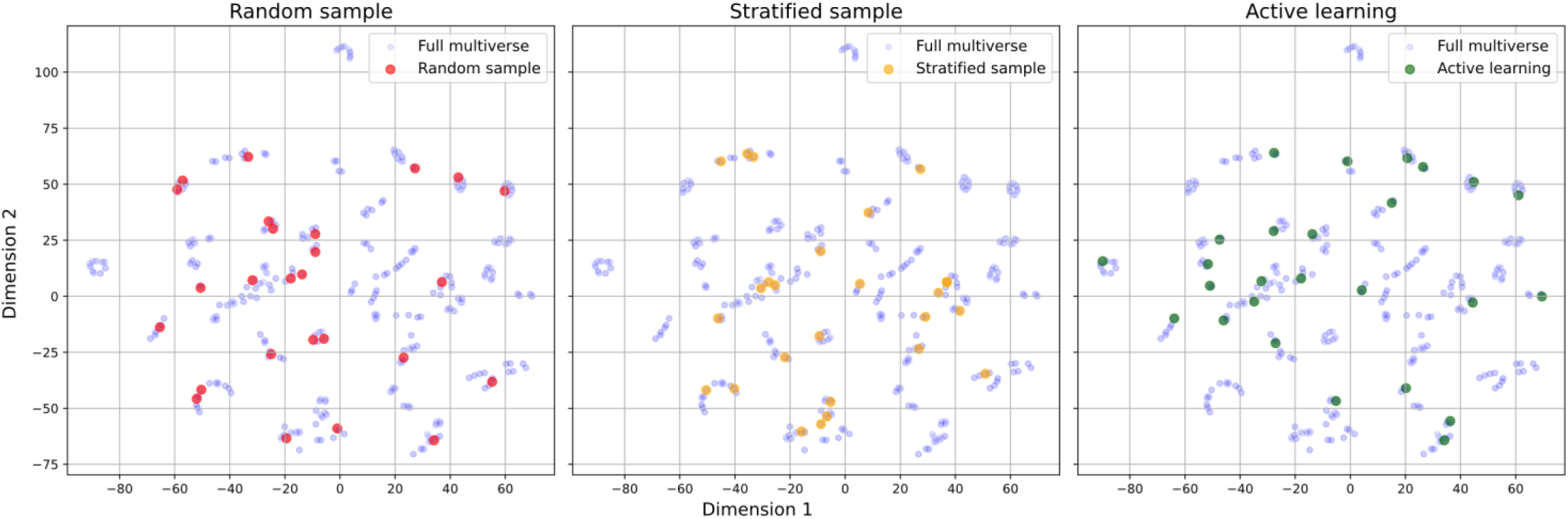
A set of scatter plots displaying the two-dimensional spatial distribution of pipelines, organised by pairwise Euclidean distances of the LPP vector of emotion conditions, for each sampling approach. Green = active learning; red = random; orange = stratified, blue = unsampled pipelines. (Pipeline sample size = 26, 5% of the full multiverse).

**Figure 5.**
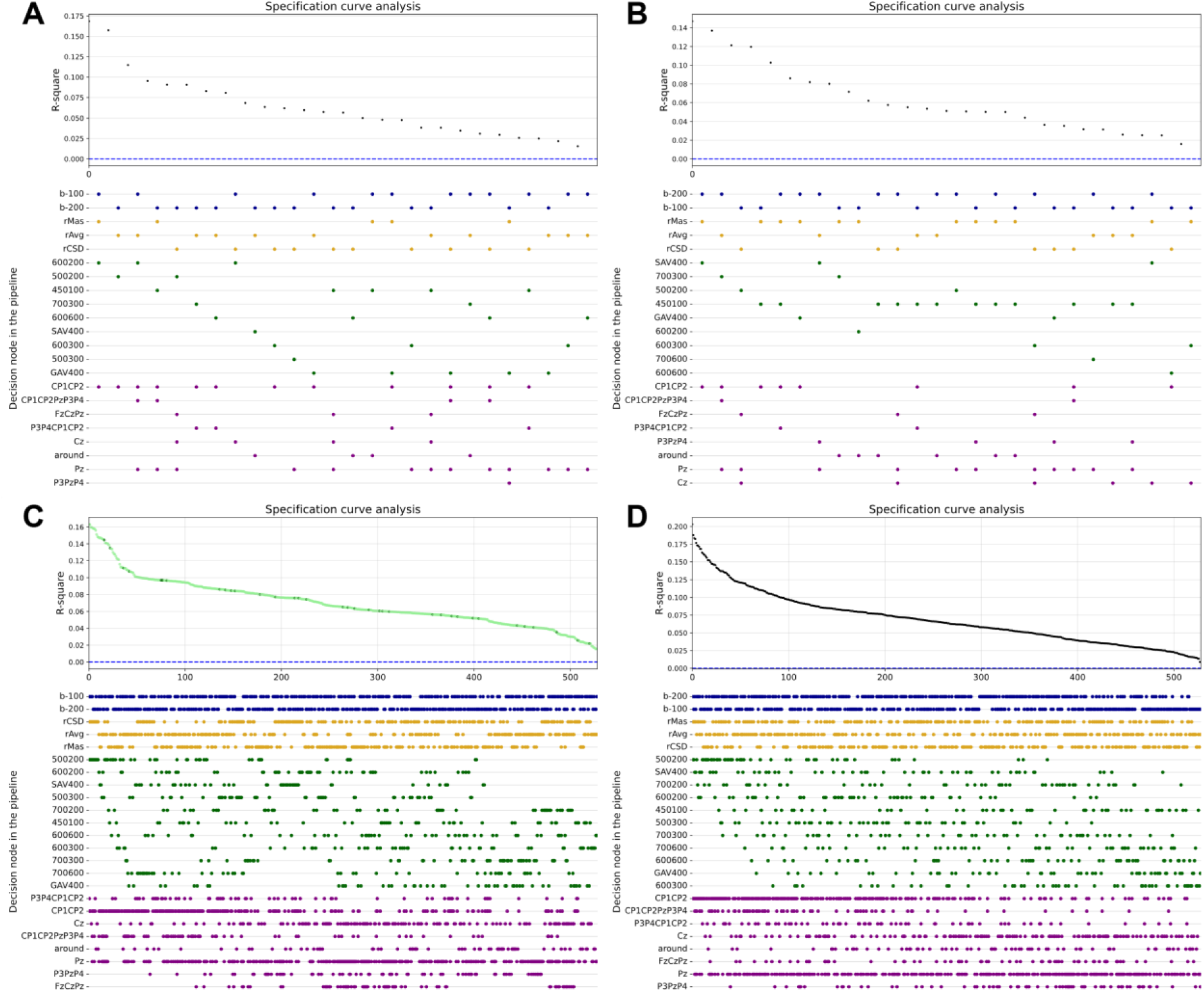
Specification curves displaying the variability in the *R^2^* across each sample, in vertical alignment with the respective pipeline options. Panel A = random sample, Panel B = stratified sample, Panel C = active learning sample, Panel D = full multiverse. Each colour in the lower specification panel corresponds to one decision node. Blue = baseline duration, yellow = reference scheme, green = time window, purple = electrode cluster. Each row within a colour represents a different option at that decision node. In the top panel of the active learning plot, dark green points denote pipelines that were directly sampled and light green points denote pipelines that were estimated. (Sample size = 26 pipelines, 5% of the full multiverse).

The specification curves (Simonsohn et al., 2020) plot the model fit for each pipeline in each sample and the full multiverse (Figure 5), alongside the parameterisation options implemented to produce them. This visualisation can be used to descriptively identify the most influential options and combinations of options in the multiverse, and the extent to which each sample communicates these influences. In the present comparison, all samples communicated the pattern observed in the full multiverse that variation in the time window and electrode cluster may be most influential on the model fit. However, the active learning sample provided the most comprehensive and easily recognisable representation, suggesting that this sampling approach may be particularly useful for multiverse analyses of these characteristics when the full multiverse cannot be computed. On the other hand, the active learning sample overemphasised the tendency of the average reference to yield both the lowest and highest model fit values, a pattern that was less pronounced in the full multiverse. This discrepancy can arise if there is an overrepresentation of a pattern in the burn-in sample and the 26 pipeline (5%) sample, leading to overfitting. While this highlights a potential artefact of active learning in small samples, it also underscores the importance of carefully selecting burn-in parameters and sample sizes to avoid misleading interpretations. Increasing the burn-in sample size or sampling a larger percentage of the multiverse can mitigate this risk, as evidenced by the reduced emphasis on this pattern when 10% of the pipelines were sampled (Supplementary 5). Future applications of active learning in ERP multiverse analyses should consider how sample size and initialisation influence generalisability.

The final comparison based on model-fit is the list of pipelines with the highest values in each sample (Table 2). While a high model fit does not necessarily indicate superior data quality, it does indicate which pipelines and which sampling approaches capture the data features that most strongly improve model performance. The random sample was the only sample to contain any of the top ten performing pipelines of the full multiverse, while the top ten performing pipelines in the active learning sample had the highest model fit on average of the three samples. While it can be expected that tuning the *k*-parameter of the active learning approach to prioritise model performance rather than exploration of the space may yield a sample of pipelines that includes pipelines with relatively high model fit values, this would potentially risk missing critical sources of variability of the full multiverse. In our use case, we set the k-parameter to prioritise exploration. This trade-off raises important considerations for researchers selecting a sampling approach. Prioritizing model performance by tuning the k-parameter of the active learning approach toward exploitative sampling could increase the likelihood of capturing the pipelines with the highest model fit, but at the cost of missing critical sources of variability in the full multiverse. Future research should consider this trade-off in line with the aim of their multiverse analysis, particularly when evaluating effects across a heterogeneous space of analytical choices. The outputs of the Multiverse Sampling Tool can help calibrate the acquisition function based on priority.

Comparing the samples on their distribution across the low-dimensional space provides insight into the diversity captured by each sample in terms of pairwise between-person differences. This is particularly useful in individual differences multiverse analyses, where it is critical to represent the variability in between-person differences across the multiverse. All three sampling approaches achieved a well-distributed selection of pipelines across the low-dimensional space (Figure 4 and Table 3), and this remained consistent across different sample sizes (Supplementary 5). This suggests that all three sampling approaches capture a representative distribution of the psychometric properties of the LPP vector for predicting between-person differences in extraversion. This finding supports the robustness of multiverse sampling to capture between-person variability in the context of the model tested in the use case multiverse, indicating that researchers may, in this context, prioritise other considerations, such as the representation of model fit.

Taken together, our results highlight that while all three sampling approaches captured to some extent the variability in model fit and between-person differences in LPP vector amplitudes of the full multiverse, the active learning sample excelled in representing the central tendency of model fit and more comprehensively conveyed the most influential LPP quantification options. However, the variability between samples in the central tendency of model fit reinforces that sampled multiverse analyses are subject to multiverse sampling uncertainty, meaning that different sampling strategies may lead to different conclusions about the robustness of results. This underscores the need for increased transparency of sampling in multiverse analyses to improve interpretability of the robustness of brain-personality relationships. To mitigate sampling uncertainty when the computation of multiple samples is not feasible, sample selection could be informed by prior comparisons, such as those presented here, to improve the generalisability of results to the full multiverse.

### Open-Source Tool

While multiverse analyses are increasingly recognised for their ability to reveal the robustness of results to analytical decisions (e.g., Short et al., 2025), the uncertainty introduced by pipeline sampling remains overlooked for large multiverse analyses where full computation is not feasible. Several tools and packages have been developed to facilitate multiverse computation (e.g., Liu et al., 2020; Sarma et al., 2024), but no tool currently exists to compare sampling approaches to quantify multiverse sampling uncertainty. The Multiverse Sampling Tool fills this gap by providing a structured framework for evaluating and reporting multiverse sampling uncertainty that is transparent and tailored to the characteristics of the given multiverse analysis. Therefore, this tool can increase the methodological transparency of multiverse analyses and reveal potential biases introduced by pipeline selection.

The Multiverse Sampling Tool executes three functionally distinct sampling procedures: random, stratified, and active learning, without requiring the full multiverse to be computed for all participants, thereby reducing the computational burden. Specifically, the random sampling approach requires only pipeline indices, the stratified sampling approach additionally incorporates option labels, and the active learning sampling approach estimates the full multiverse using a training subset of participants. In addition to generating three pipeline samples, the tool generates a series of tabular and graphical outputs that describe the model fit distributions for each sample and, for active learning, the predicted model fit for the unsampled pipelines and the dispersion of pipelines across a low-dimensional space of pairwise distances between individuals. These outputs provide users with a transparent report of the multiverse sampling uncertainty specific to their analysis, facilitating more informed and methodologically rigorous sample selection.

While the active learning approach in the tool is adapted from an existing script (Dafflon et al., 2022), it extends beyond the original implementation, which was primarily designed to optimise pipeline selection in active learning rather than evaluate sampling uncertainty across approaches. The original active learning script focused on tuning the acquisition function and sample size of the active learning approach to sample pipelines by reporting the mean average error, the distribution across the low dimensional space, and Bayesian optimisation performance across different random starting conditions. In contrast, the Multiverse Sampling Tool focuses on comparing the active learning sample with alternative sampling approaches, to evaluate the variability in the central tendency of the model fit and spatial distribution across samples. Thus, the original study (Dafflon et al., 2022) can be used to inform the parameter tuning of the active learning, and our tool can be used to execute the active learning sample and compare it to other samples to report multiverse sampling uncertainty.

The Multiverse Sampling Tool is designed to systematise and bring transparency to the multiverse sampling process, providing a framework for evaluating and reporting multiverse sampling uncertainty and supporting the primary goal of multiverse analysis, which is to make uncertainty explicit (Hoffmann et al., 2021). Specifically, when the script and outputs are shared, the tool makes transparent the sampling parameter settings, and the uncertainty that exists in the robustness of the multiverse results across different pipeline samples. By providing an automated and structured workflow, the tool ensures that sample selection aligns with study-specific data characteristics, experimental design, the EEG metric of interest, and the statistical model being fitted. Additionally, it can be integrated into preregistrations, allowing researchers to define sampling thresholds and selection criteria in advance with direct reference to the tool. This structured approach enhances methodological rigour and reproducibility, while fostering greater confidence in the generalisability of sampled multiverse analysis results. By making multiverse sampling uncertainty explicit, this tool advances the field toward more transparent, interpretable, and robust EEG multiverse analyses.

### Limitations

While the Multiverse Sampling Tool provides a valuable framework for evaluating *multiverse sampling uncertainty*, it is not without limitations. First, the script relies on *R^2^*-values as the primary metric for evaluation, interpreting them as either model fit (for sampled pipelines) or predicted model fit (for unsampled pipelines in active learning). This dual use can complicate direct comparisons between sampling approaches and requires careful interpretation to avoid conflating these metrics. Further, the larger the pipeline sample size, the more certain the estimates for comparison. Second, while the tool provides rich outputs for evaluating and visualising multiverse sampling uncertainty, it does not automate decision-making, leaving the selection of parameters and selection of a final sample for reporting to the user. While this flexibility allows for sampling procedures tailored to the study design, it also introduces subjectivity. However, this subjectivity can be made transparent by sharing the implemented script within the tool alongside the reported multiverse analysis results. Third, to compute the active learning-based sample, we rely on dimensionality reduction techniques, which may influence the convergence of active learning given that is highly dependent on the distribution of the pipelines across the space. However, different dimensionality reduction methods can be explored, as shown in Supplementary 4, to ensure the selection of an approach that best preserves the structure of between-person pipeline dissimilarities, with a useful broad distribution across the space.

The sampling script is open to the community and can be modified, allowing the functionality to evolve with the field, and ensuring that the script has the potential to remain a useful resource, and meet the diverse needs of the research community.

## Conclusion

The Multiverse Sampling Tool provides an easy-to-implement framework to compare three functionally distinct sampling approaches, enhancing transparency in large-scale EEG multiverse analyses that require sampling and are subject to the typically unreported *multiverse sampling uncertainty*. The tool’s adjustable parameters to provide versatility for a variety of EEG multiverse analyses, allowing researchers to tailor sampling strategies to their specific study design and computational restraints. Furthermore, the reported use case multiverse provides a reference point for future multiverse analyses with similar characteristics, helping researchers to select an appropriate sampling approach when computation of multiple samples is not feasible. By systematically evaluating multiverse sampling uncertainty, the Multiverse Sampling Tool provides a means of improving transparency and reproducibility of large-scale EEG multiverse analyses, strengthening the reliability of conclusions drawn from samples of complex analytical spaces.

## Supplementary 1: The Structure and Modifiability of the Main.py Script

It is organized into five sections: preliminary installations, preparation, functions, sampling, and results.

### Section A: Preliminary Installations

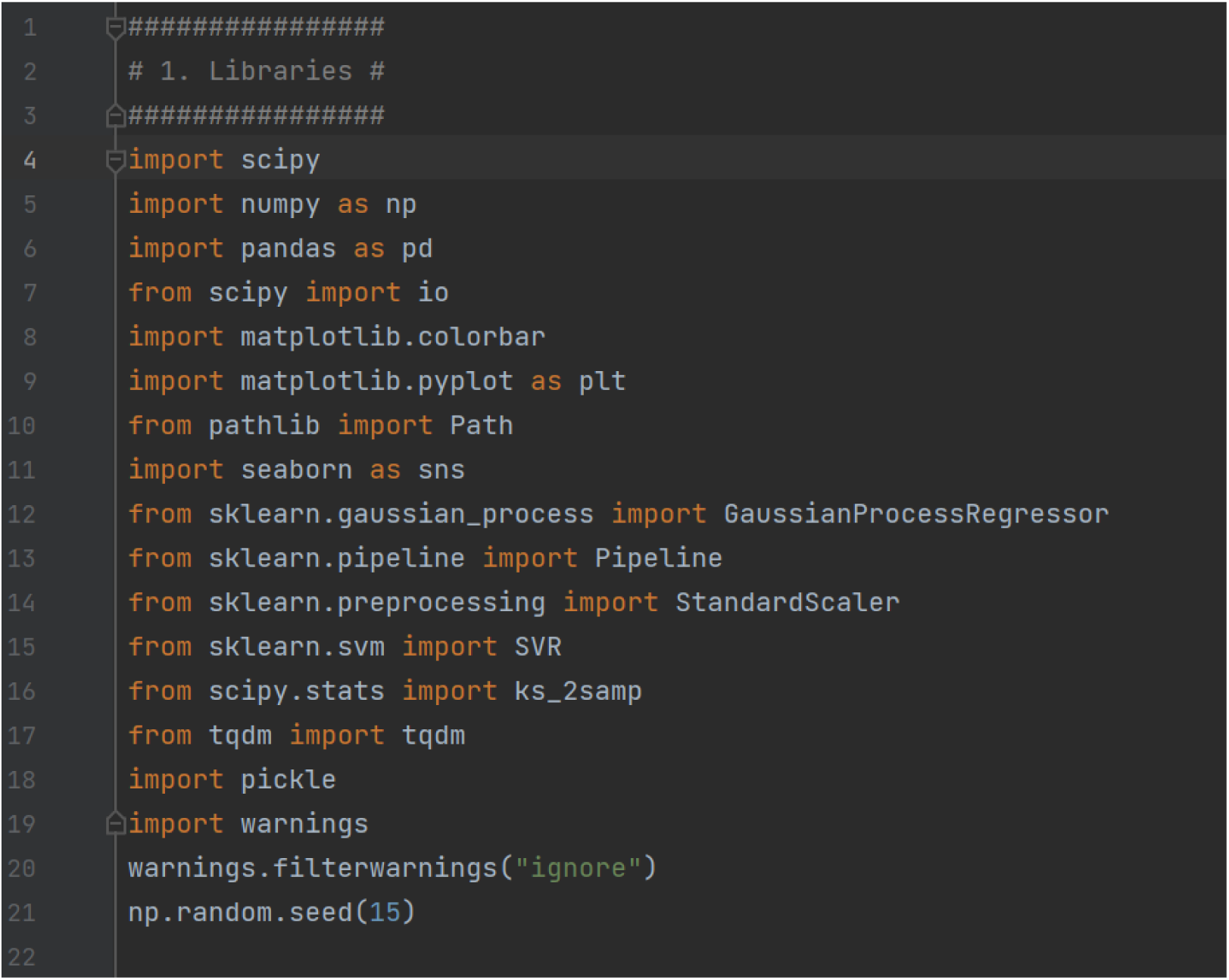

a. scipy, scipy import io, scipy-stats import ks_2samp. This is a library that provides statistical functions used in the script. The ks_2amp function from the scipy.stats module is used in the script to perform the Kolmogorov-Smirnov test.
b. numpy as np. In this script, NumPy is used for setting a seed for random number generation to ensure reproducibility and for performing random sampling.
c. pandas as pd. The panda library of data structures and analysis tools is used in the script to create and manipulate data frames, which are used to store and handle the configurations for different decision nodes and the results of the sampling and statistical tests.
d. matlabplot.colorbar, matplotlib.pyplot as plt. Matlabplot is a plotting library for creating visualisations. It’s plotting functions are used in this script, for example for the scatter plots.
e. pathlib import Path. Path is used in the script to construct paths for data and output directories that is platform independent.
f. seaborn as sns. Seaborn is a statistical data visualisation library based on matplotlib. In this Python script, seaborn is used to create the raincloud plots.
g. sklearn.gaussian_process import GaussianProcessRegressor, sklearn.pipeline import Pipeline, sklearn.preprocessing import StandardScaler, sklearn.svm import SVR. Th sklearn library is used for machine learning and contains simple and efficient tools for predictive data analysis. In this script, it’s used for the Gaussian process regressor and the support vector regressor, which are part of the active learning sampling procedure.
h. tqdm. This is a library for creating progress bars for loops, and is used in this script to provide a visual reference for the progress of the sampling loops.
i. pickle. This Python module for serialising and deserialising object structures is used to save load previously saved data, for example. ‘PredictedAcc’.
j. Warnings, warnings.filterwarnings(“ignore”). This module controls the display of warnings, and this script uses it to suppress warnings that might clutter the output.
k. Np.random.seed(15) sets the seed for NumPy’s random number generator to 15. This ensures that any subsequent random operations produce reproducible results.

### Section B: Preparing data

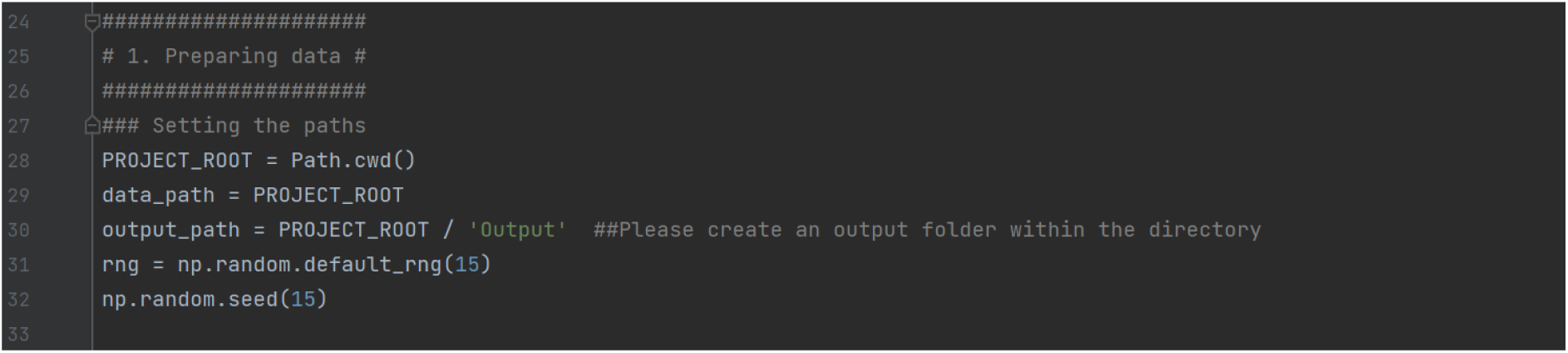

B1: Setting the paths

a. PROJECT_ROOT = Path.cwd(). This sets the PROJECT_ROOT variable to the current working directory using the cwd() function. The path entered should include the data files required for the script to run.
b. Data_path = PROJECT_ROOT. This assigns the PROJECT_ROOT directory to the data_path, as all data files should be located within the PROJECT_ROOT directory that was set above.
c. Output_path = PROJECT_ROOT / ‘Output’. This creates an output path by appending an ‘Output’ directory to the PROJECT_ROOT directory.
d. rng = np.random.default_rng(11). This initializes a new random number generator object with a fixed seed (15) for reproducibility.
e. np.random.seed(15). Sets the seed for the random number generator to 15, ensuring reproducibility.

B2: Import data

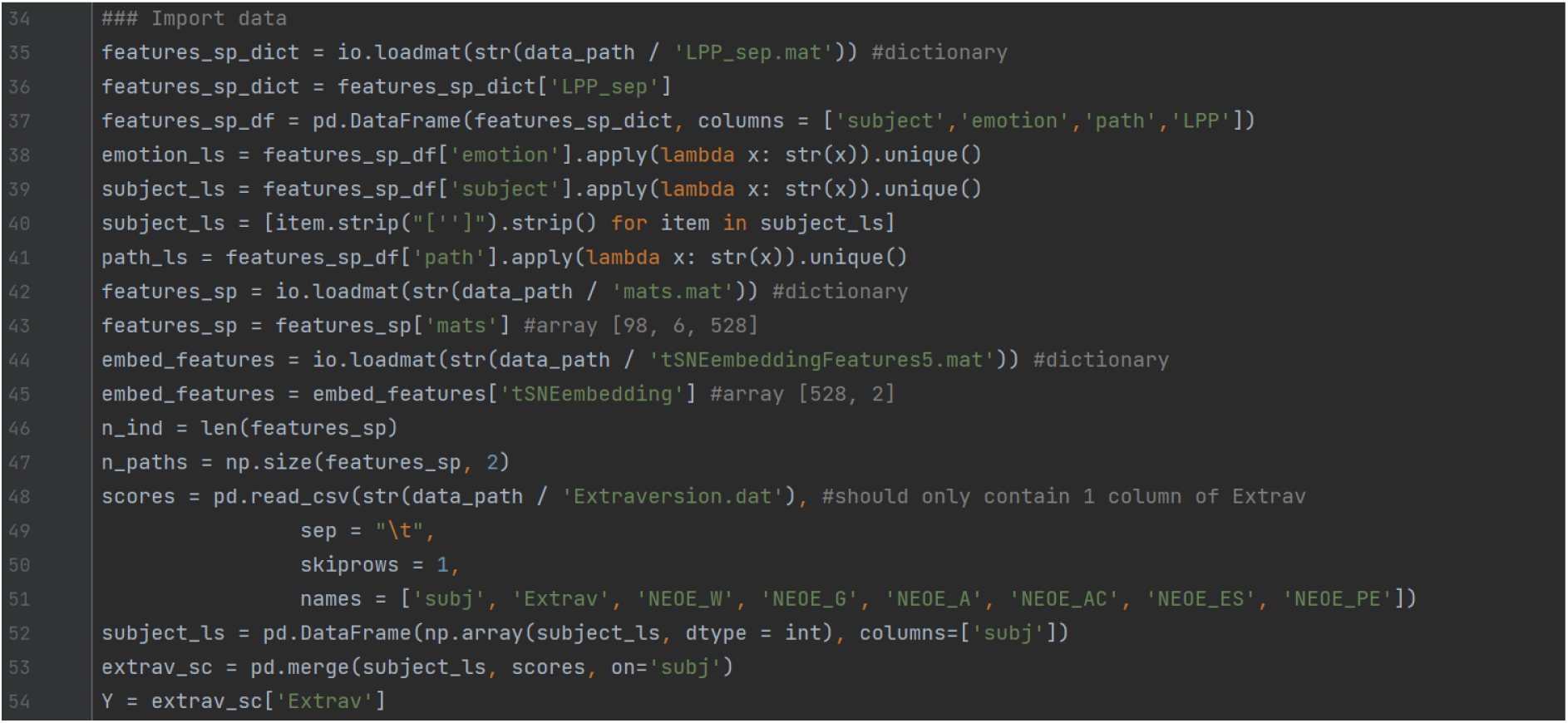

This subsection handles data importation, preparation and preliminary transformations:

a. Imports the LPP_sep.mat file and specifies a dataframe (features_sp_df) with specific column names (subject, emotion, path, LPP);
b. Extracts unique values from the columns: emotion_ls is a list of unique emotions, subject_ls is a list of unique subjects (with formatting adjustments to remove extra characters), and path_ls is a list of unique paths:
c. Loads mats.mat and specifies that it is an array (features_sp) with shape [98, 6, 528] (98 participants, 6 conditions, 528 pipelines);
d. Loads tSNEembeddingFeatures5.mat and specifies it as an array with shape [528, 2] (528 data points embedded in a 2 dimensional space);
e. Stores dimensional information of features_sp;
f. Loads extraversion scores from Extraversion.dat, which contains columns for participant ID, extraversion score, and additional personality scores that are not used in the current script, merges the data based on participant ID and extracts extraversion as the target variable.B3: Forked paths

B3: Setting the forking paths

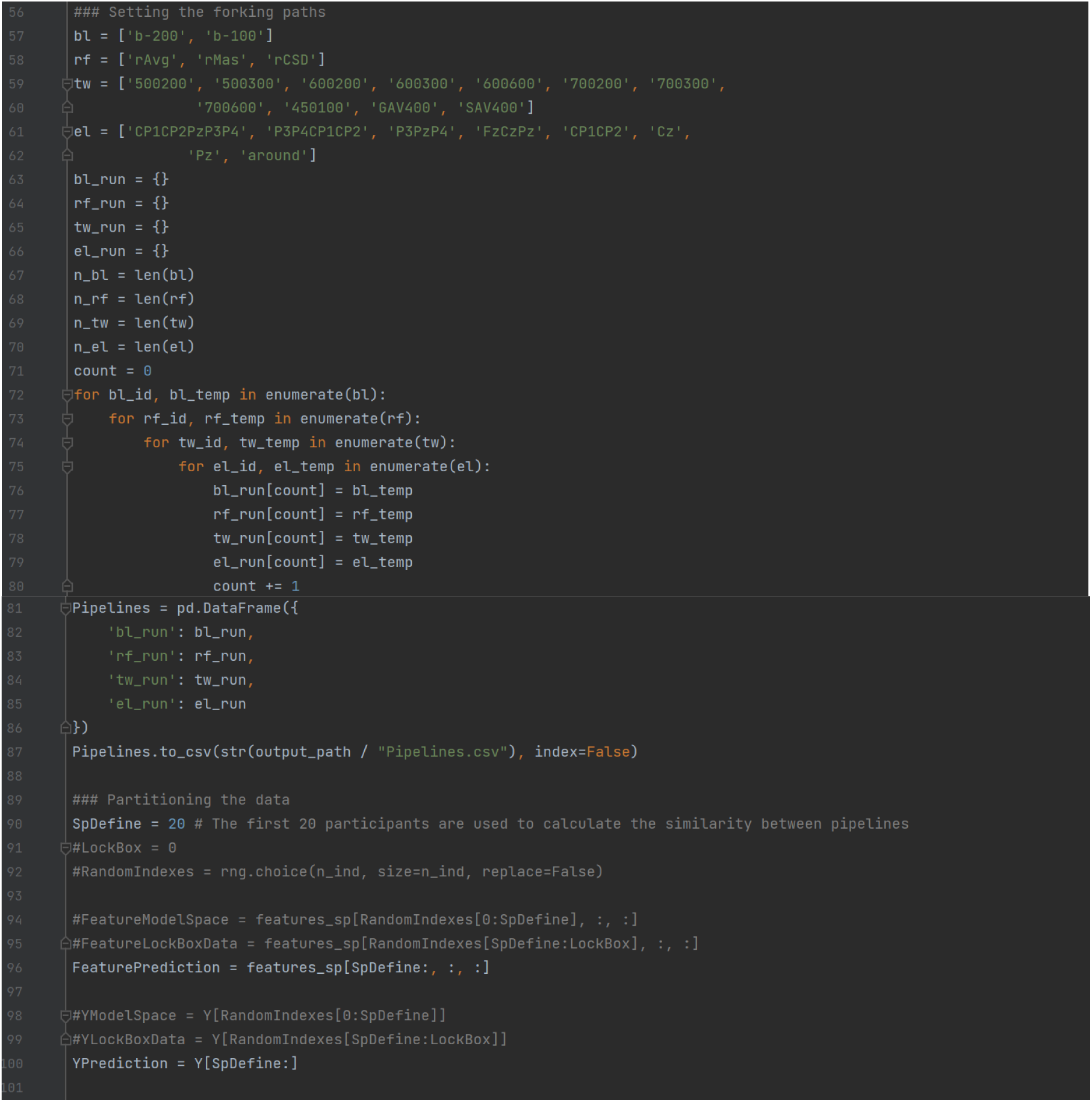

a. Defines lists for different decision nodes of the multiverse analysis: electrodes, reference, time window and electrodes (bl, rf, tw, el). These should be replaced with names that represent the decision nodes of the user’s multiverse.
b. Initialises empty dictionaries for each of the decision nodes to store the generated configurations (bl_run, rf_run, tw_run, el_run). The prefix of these labels should be consistent with the labels you assigned in a for ease of understanding;
c. Counts and stores the number of options available for each decision node (n_bl, n_rf, n_tw, n_el; how many baselines, how many reference schemes, how many time windows and how many electrodes). The suffix of these labels should be consistent with the labels you assigned in sub-step ‘a’.
d. Nested for loop that iterates over each list (assigned in ‘a’), creating a Cartesian product of all configurations. Inside the loop, each configuration combination is assigned a unique count as its key and the corresponding values from each list are stored in their respective dictionaries initialized in ‘b’). The enumerate function is used to iterate over each list, providing the index (bl_id, rf_id, tw_id, el_id) and the value (bl_temp, rf_temp, tw_temp, el_temp). Again, the prefixes should be consistent with the labels assigned in ‘a’.
e. Converts four dictionaries (bl_run, rf_run, tw_run, el_run) into a dataframe, whre each row represents a unique pipeline configuration and saves the dataframe as a CSV file (Pipelines.csv).
f. Defines a cutoff point (20 participants in this case) for splitting the data (SpDefine). The data is partitioned into features for prediction (FeaturePrediction), which contains the data for participants after the first 20, and target for prediction (YPrediction), which contains the corresponding Y (extraversion scores) for these participants.
g. Optional: LockBox could define an additional subset for cross validation, RandomIndexes could randomly shuffle participant indices.

### Section C: Exhaustive analysis

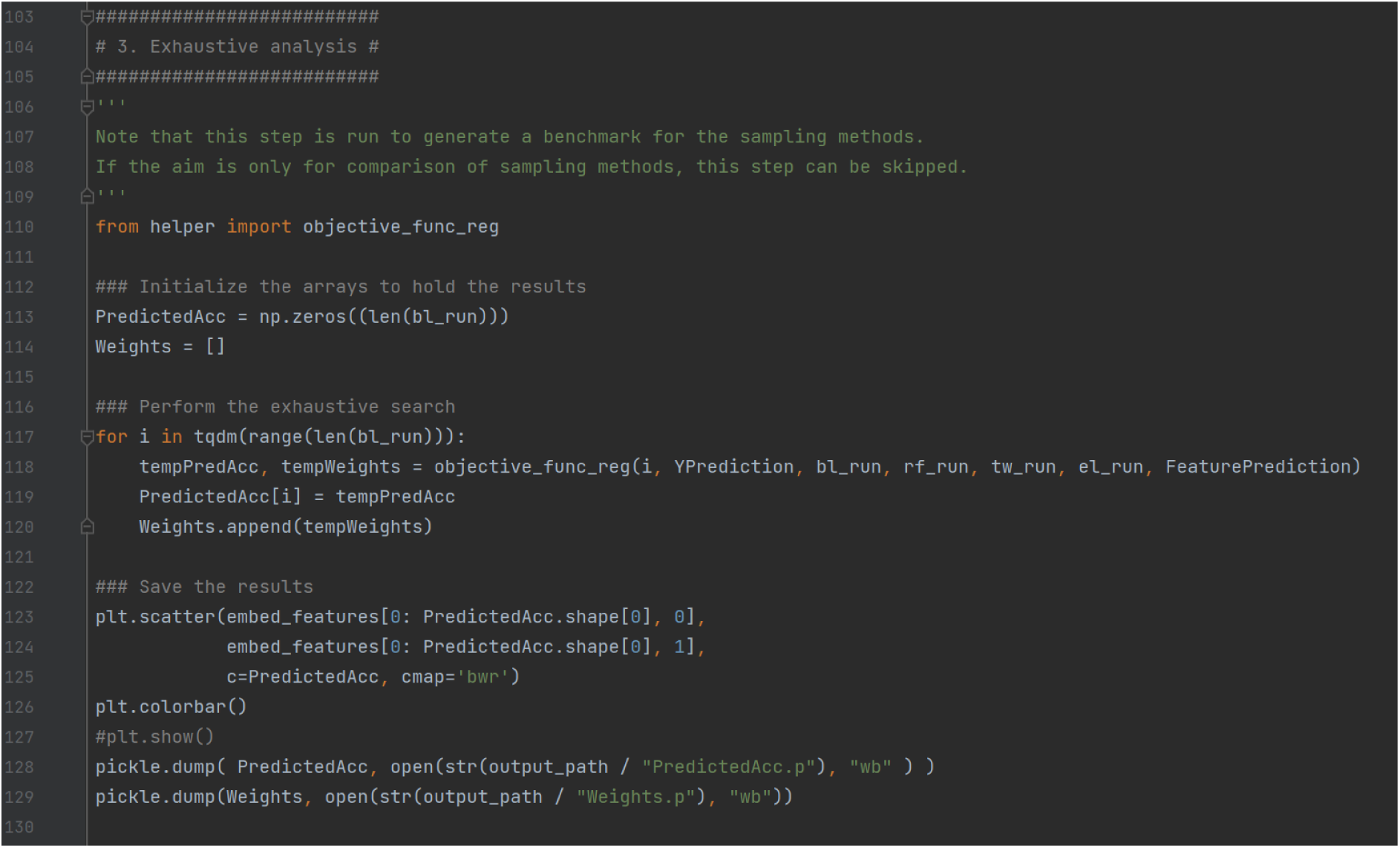

This section of the main script performs an exhaustive analysis to evaluate all pipeline configurations and generates a benchmark for later comparison with samples. This is not necessary and can be skipped when the multiverse is too large to compute exhaustively. This section calculates the performance (*R^2^*) for each pipeline and saves the results for later analysis.

a. Initializes arrays for results: PredictedAcc to store the performance (*R^2^*) for each pipeline, and Weights to store feature weights for each pipeline;
b. Performs the exhaustive search by iterating through all pipelines, calling the helper function objective_func_reg and imputing the pipeline index (i), the extraversion scores for prediction (YPrediction), the pipeline parameters (bl_run, rf_run, tw_run, el_run), and the features corresponding o the prediction subset (FeaturesPrediction). It outputs the *R^2^* score of the pipeline (tempPredAcc) and the feature weights from the linear regression model (tempWeights), and stores the results in PredictedAcc and Weights;
c. Creates a scatterplot of the distribution of pipeline performance across the two-dimensional space;
d. Saves the results.

### Section D: Comparison of sampling methods

Section D is a large code chunk that computes and compares the active learning, stratified and random samples. This section is explained in smaller sub-chunks.

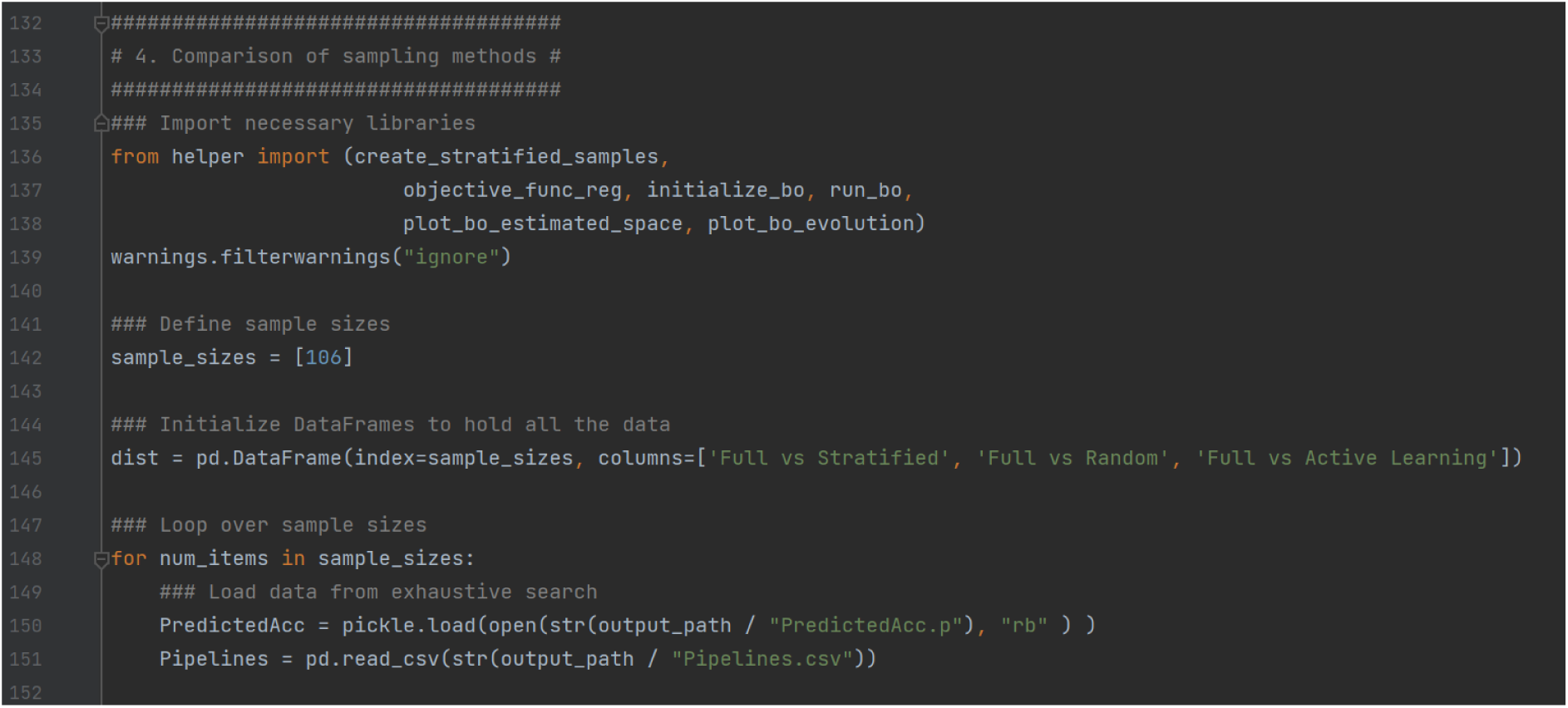

a. Imports key functions from the helper script;
b. Defines sample sizes (the example in the code image is 106, which means that each sampling method will select 106 pipelines - 20% of the 528 - for evaluation);
c. Creates an empty dataframe (dist) to store similarity metrics between the full exhaustive benchmark and the sampled subsets. The columns represent comparisons of the exhaustive multiverse to the stratified sample, the random sample and the active learning sample;
d. Initiates a loop to iterate over the defined sample sizes;
e. Loads the previously computed benchmark data from the exhaustive computation, where PredictedAcc contains the *R^2^* for all pipelines, and Pipelines is a dataframe listing all pipeline configurations;

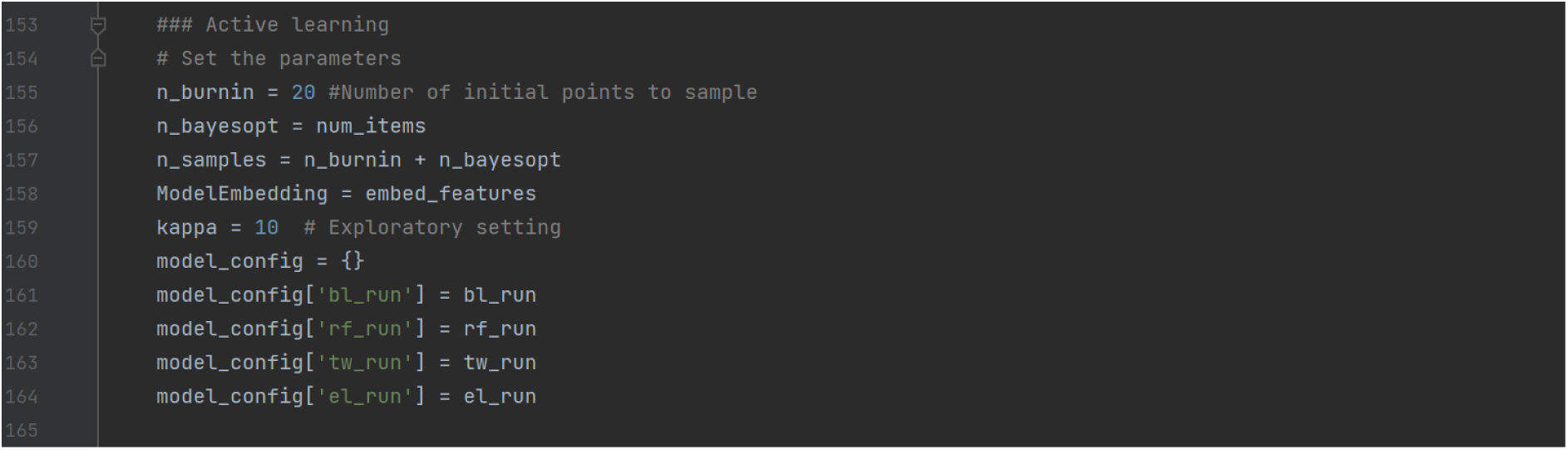

This sub-chunk prepares the parameters and configurations for implementing active learning.

a. Sets the active learning parameters: n_burnin (the number of randomly selected pipelines to evaluate before starting Bayesian optimization), n_bayes_opt (the number of additional pipeline evaluations to perform using Bayesian optimization), n_samples (total number of pipelines to evaluate, which is n_burnin + n_bayesopt);
b. Assigns the precomputed t-SNE embeddings (embed_features) of pipelines to ModelEmbedding;
c. Sets the exploration vs. exploitation trade-off parameter for Bayesian optimization. A higher kappa value, 10 as cited by Dafflon et al., (2022) provides a more explorative sample, favoring the selection of data points with greater uncertainty, whereas a lower kappa value, such as 0.1 as cited by Dafflon et al., (2022) provides a more exploitative sample, favoring data points that are predicted to improve the model performance based on current knowledge of the sample so far;
d. Creates a dictionary (model_config) to map pipeline components to their respective configurations (bl_run, rf_run, tw_run, el_run), which provides a structured way to pass all pipeline configurations to the Bayesian optimization process.

As with the other parameters set in the script, the user can adjust these parameters before running the script based on their data and multiverse analysis goals. We would like to reiterate that the active learning sampling feature in the Python script is a modification of that published by Dafflon et al. (2022), available [here](https://github.com/Mind-the-Pineapple/into-the-multiverse).

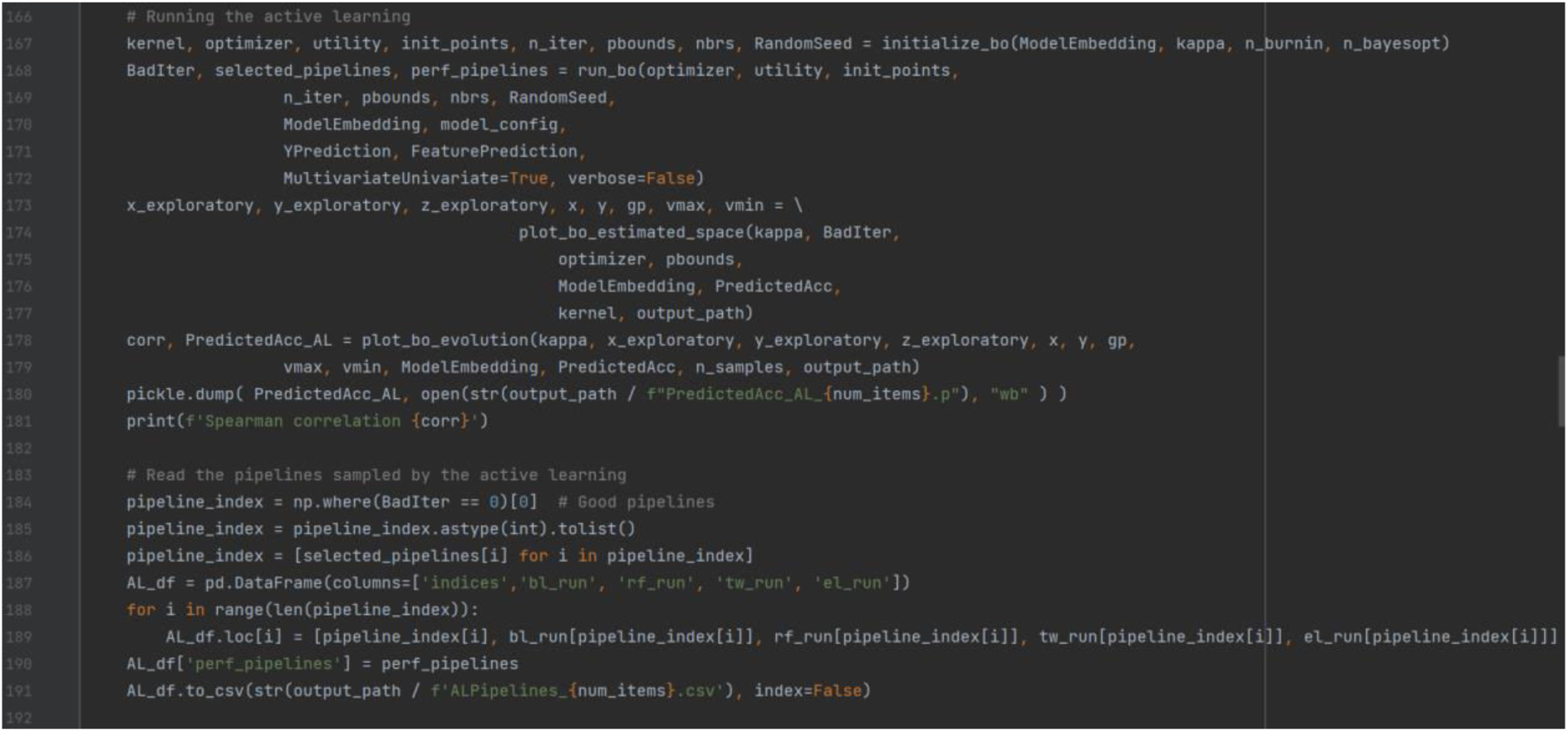

This sub-chunk executes the active learning sampling. It includes steps for initializing the Bayesian optimization, running the optimization, visualizing the results, and saving the sampled pipelines and their performance:

a. Calls the initialize_bo function from the helper script to set up the Bayesian optimization with the following outputs:

a. Gaussian process regressor kernel: In the use case, a combination between a matern and a white noise kernel is used, which was default in the script as downloaded from Dafflon et al., (2022). The default can be adapted if the user wishes to modify the smoothness and the noise variance;
b. Optimizer object;
c. Utility function for selecting the next point in the space;
d. Number of random initialization points (burn in sample);
e. Number of iterations for the Bayesian optimization;
f. Parameter bounds for the optimization process;
g. Nearest neighbor model for pipeline embeddings;
h. Seed for reproducibility.
b. Executes the Bayesian optimization process using the run_bo function, which selects pipelines iteratively based on the utility function and evaluates selected pipelines using the objective_func_reg function;
c. Visualizes the search space using the plot_bo_estimated_space function;
d. Visualises the Bayesian optimization evolution using the plot_bo_evolution function;
e. Saves the results and the sampled pipelines (identified pipelines selected with acceptable performance - BadIter == 0, creates a dataframe (AL_df) to store the indices of selected pipelines, pipeline configurations, and performance scores).

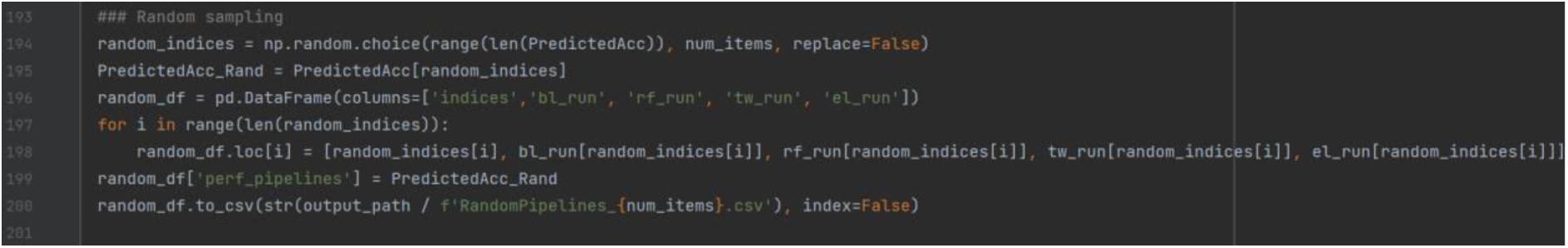

This sub-chunk implements random sampling, where pipelines are selected randomly, and evaluates the performance of the randomly sampled pipelines:

a. Randomly selects num_items unique pipeline indices from the full set of pipelines, ensuring that no index is selected more than once;
b. Extracts the performance scores (PredictedAcc) of the randomly selected pipelines (random_indices);
c. Creates a dataframe (random_df) for the randomly selected pipelines with columns to store indices and corresponding pipeline cofigurations (bl_run, rf_run, tw_run, el_run), and fills the dataframe with the information of the sampled pipelines;
d. Adds a column to the dataframe that contains the performance scores (PredictedAcc_Rand);
e. Saves the dataframe as a CSV file.

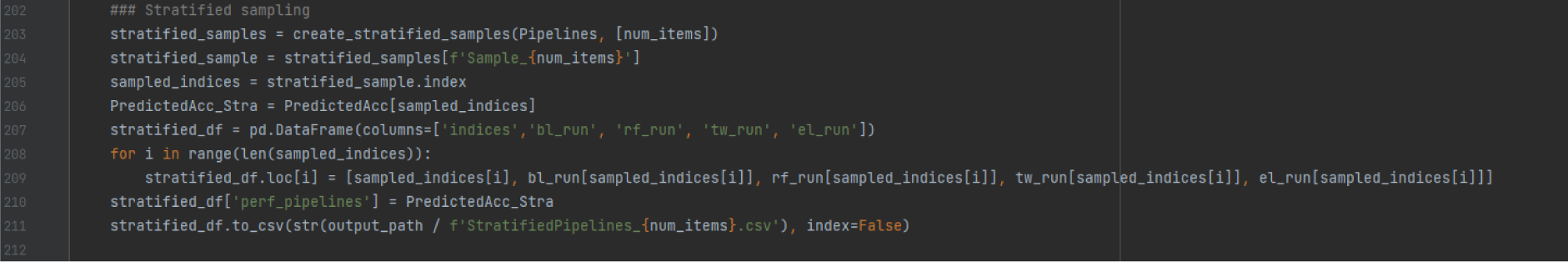

This sub-chunk implements stratified sampling by selecting pipelines proportionally to represent the diversity of the entire identified multiverse, and evaluates their performance:

a. Uses the create_stratified_samples function from the helper script with inputs of the dataframe that contains all pipelines (Pipelines), and the list of desired sample sizes (num_items). It outputs a dictionary (stratified_samples) where each key corresponds to a sample size;
b. Collects the stratified sample corresponding to num_items, and extracts the indices of the sampled pipelines (sampled_indices);
c. Retrieves the performance scores (PredictedAcc) of the pipelines in the stratified sample using their indices;
d. Initializes an empty dataframe (stratified_df) with columns for the indices of sampled pipelines and the pipeline configurations, and fills the dataframe with the information about the sampled pipelines;
e. Adds a column (perf_pipelines) to the dataframe containing he performance scores (PredictedAcc_Stra) of the pipelines in the sample;
f. Saves the dataframe of the sampled pipelines.

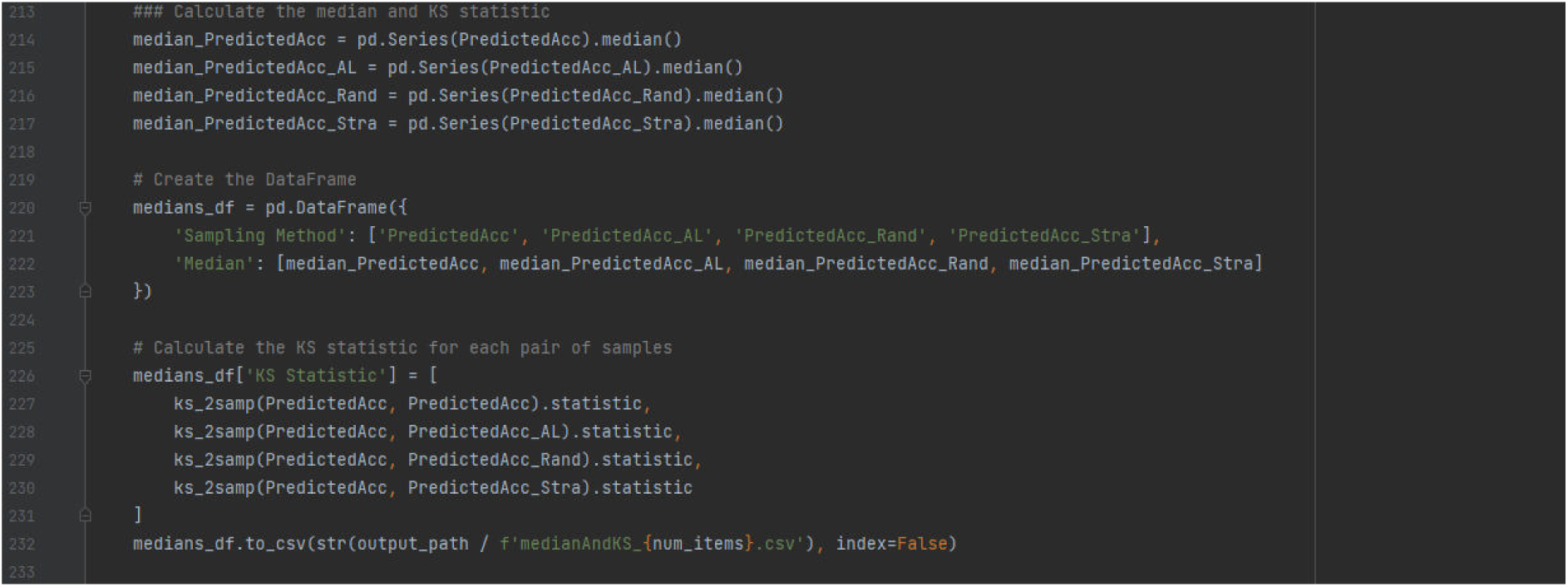

This sub-chunk computes and compares the median performance and the Kolmogorov-Smirnov (K-S) statistic for pipeline performance across different samples:

a. Calculates the median performance for the full multiverse if this is computed, the active learning sample, random sample, and stratified sample;
b. Creates a dataframe to store the median performance for each sample;
c. Calculates the K-S statistic for each sample vs. the full multiverse;
d. Saves the results.

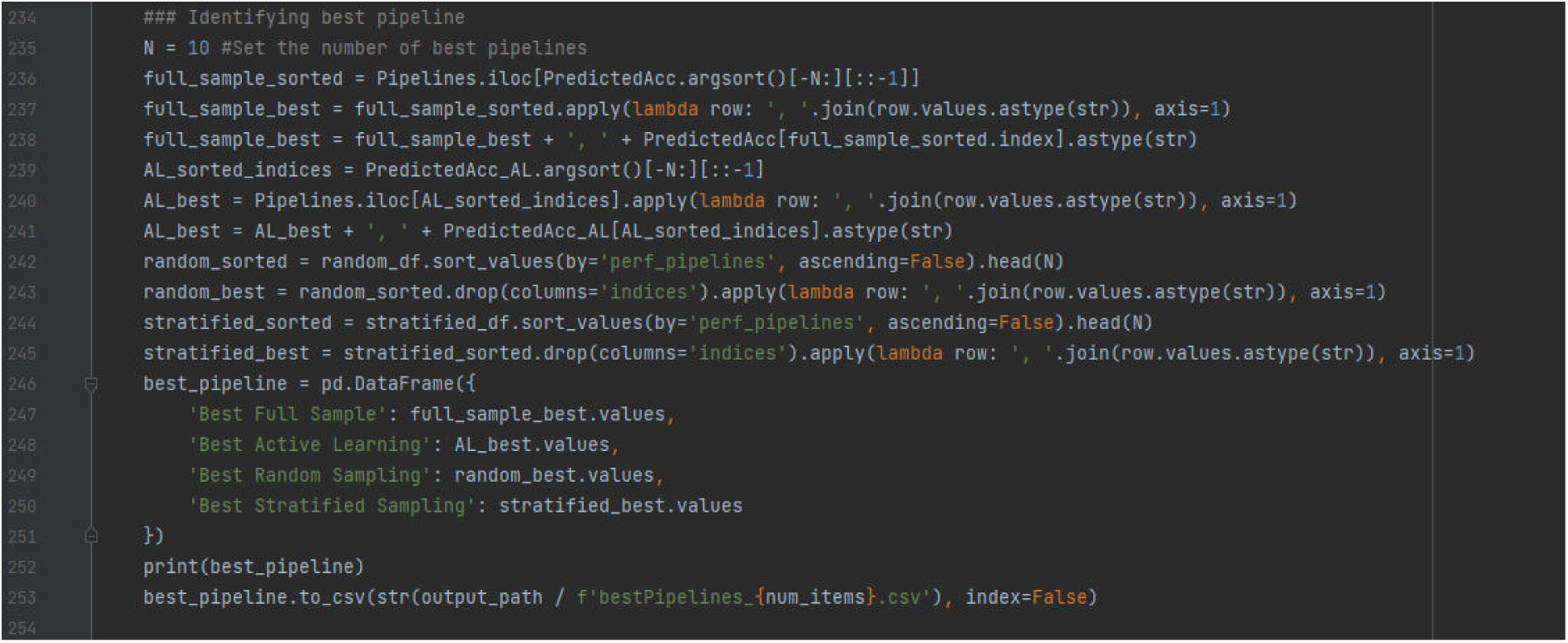

This sub-chunk identifies and compares the top performing pipelines for the full multiverse and each of the three samples:

a. Specifies the number of pipelines to list (e.g., top 10 performing pipelines for each);
b. Extracts the top performing pipelines from the full multiverse and each sample by sorting the indices of the PreditedAcc array in ascending order, selecting the top N indices in descending order of performance, retrieving the pipeline configurations corresponding to these, converting each pipeline into a comma-separated string, and appending the performance scores to the pipeline strings;
c. Creates a dataframe of the top performing pipelines for the full multiverse and each sample with columns for the full multiverse and each sample;
d. Saves the results.

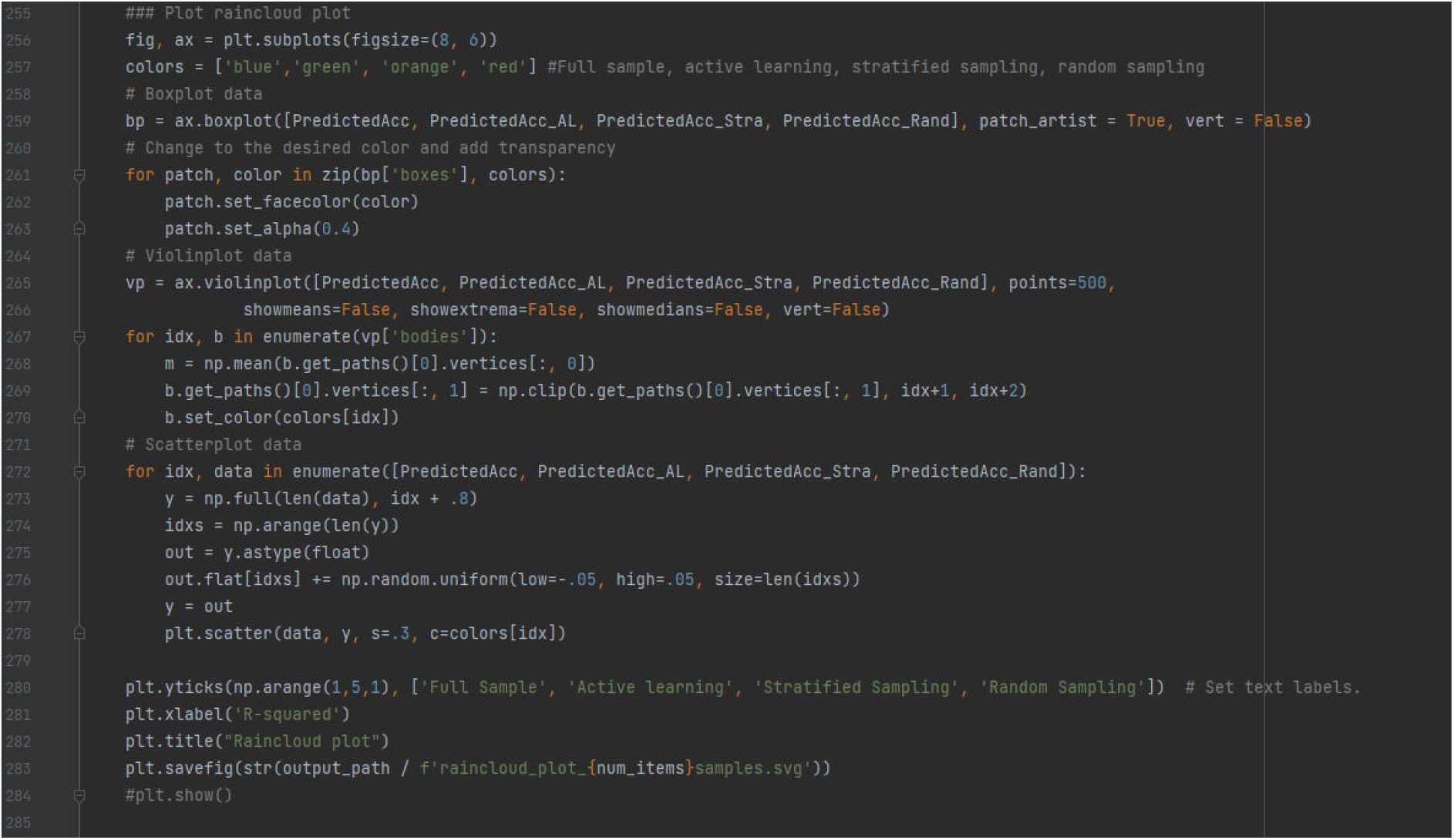

This sub-chunk visualizes the performance distribution of pipelines sampled by the different methods using a raincloud plot:

a. Initializes a plot with specified figure size and single axis and defines the color palette for the sampling methods;
b. Creates a horizontal boxplot for the distributions of the PredictedAcc for the full multiverse and each of the samples;
c. Loops through the boxplot patches and assigns color and transparency;
d. Creates a horizontal violin plot for the same distributions as the boxplot and iterates through the violin plot to assign color and alignment adjustment;
e. Adds jittered scatter points to represent individual data points in each distribution;
f. Sets the axis labels;
g. Saves the plot as an SVG file.

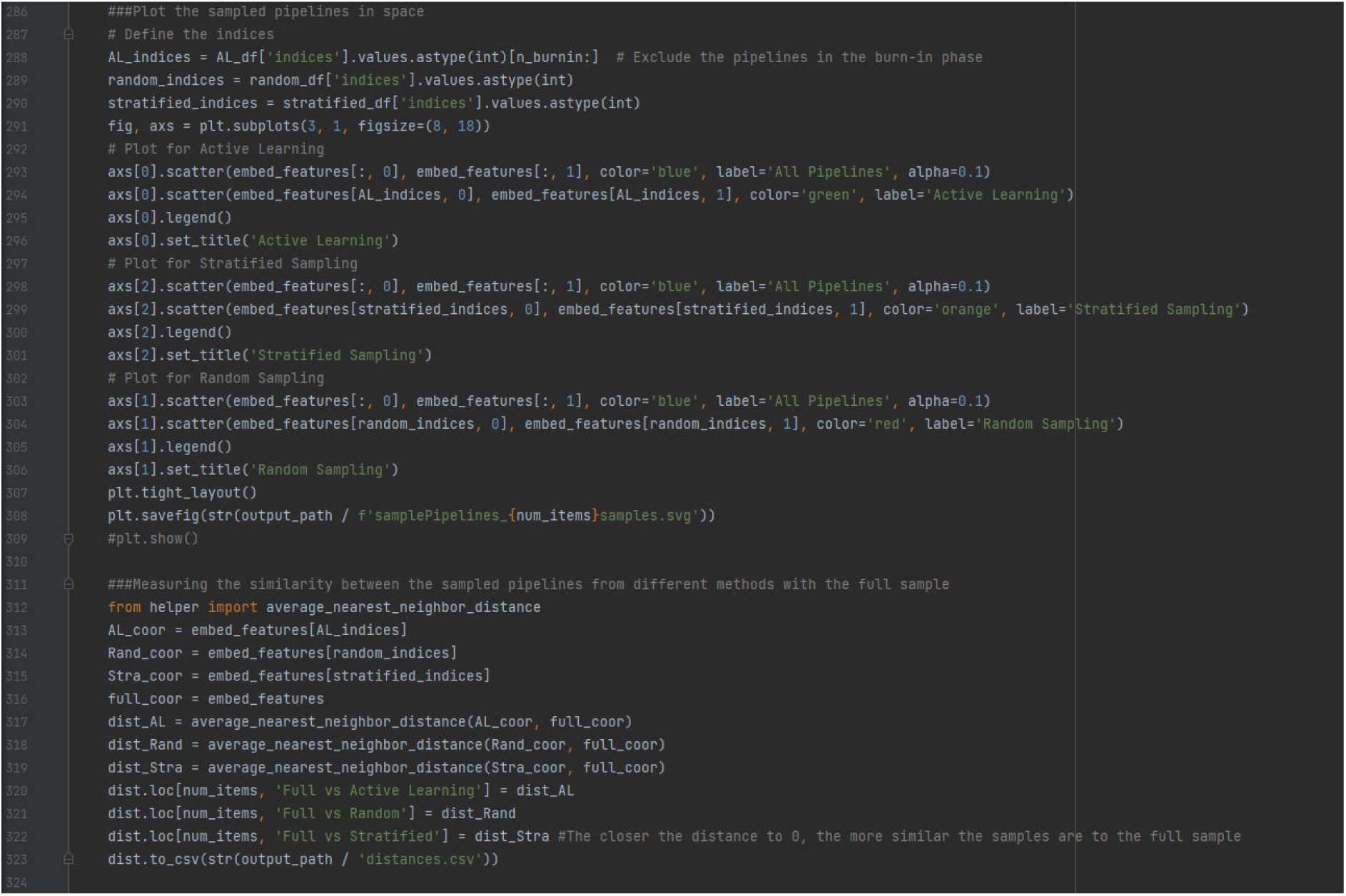

This sub-chunk visualizes how pipelines selected by the different sampling methods are distributed in the two-dimensional space relative to the full multiverse, and calculates similarity metrics between the sampled pipelines and the full multiverse:

a. Extracts indices of pipelines sampled by each method (excluding the burn-in pipelines for the active learning sample);
b. Creates a figure with three subplots stacked vertically (one subplot for each sample), specifying the figure size;
c. For each subplot, all pipelines are plotted in the two-dimensional space as faint blue points, and the sampled pipelines are colored as green for the active learning sample, orange for the stratified sample, and red for the random sample;
d. Saves the plot.
e. Measures the similarity of each ample with the full sample by extracting coordinates of sampled pipelines and all pipelines from the embedding space, uses the average_nearest_neighbor_distance function from the helper script to compute the similarity of each sample of pipelines to the full multiverse;
f. Saves the results.

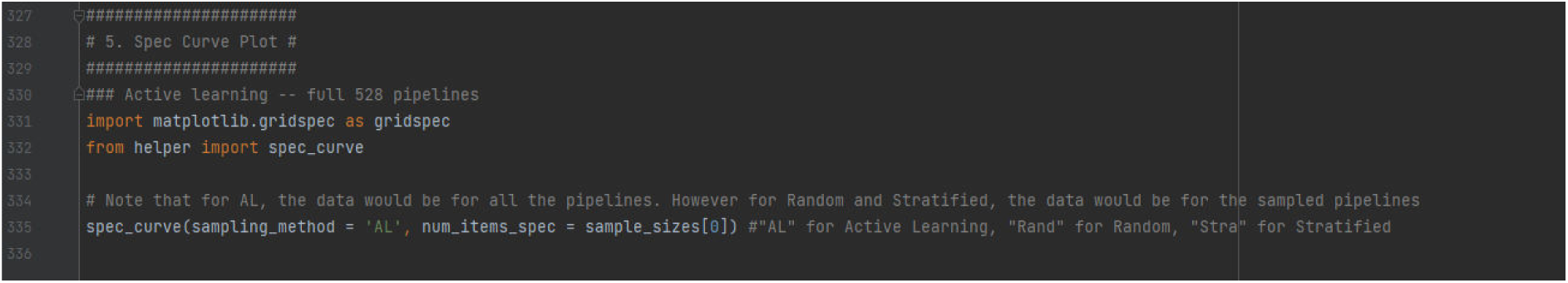

This sub-chunk generates a specification curve plot to visualize the performance of pipelines across different sampling methods. Uses the spec_curve function to create the specification curve, with AL entered for active learning, Rand entered for random, and Stra entered for stratified.

## Supplementary 2: The Structure and Functionality of the MultiverseSampling_Helper.py Script

### Preliminary Installations

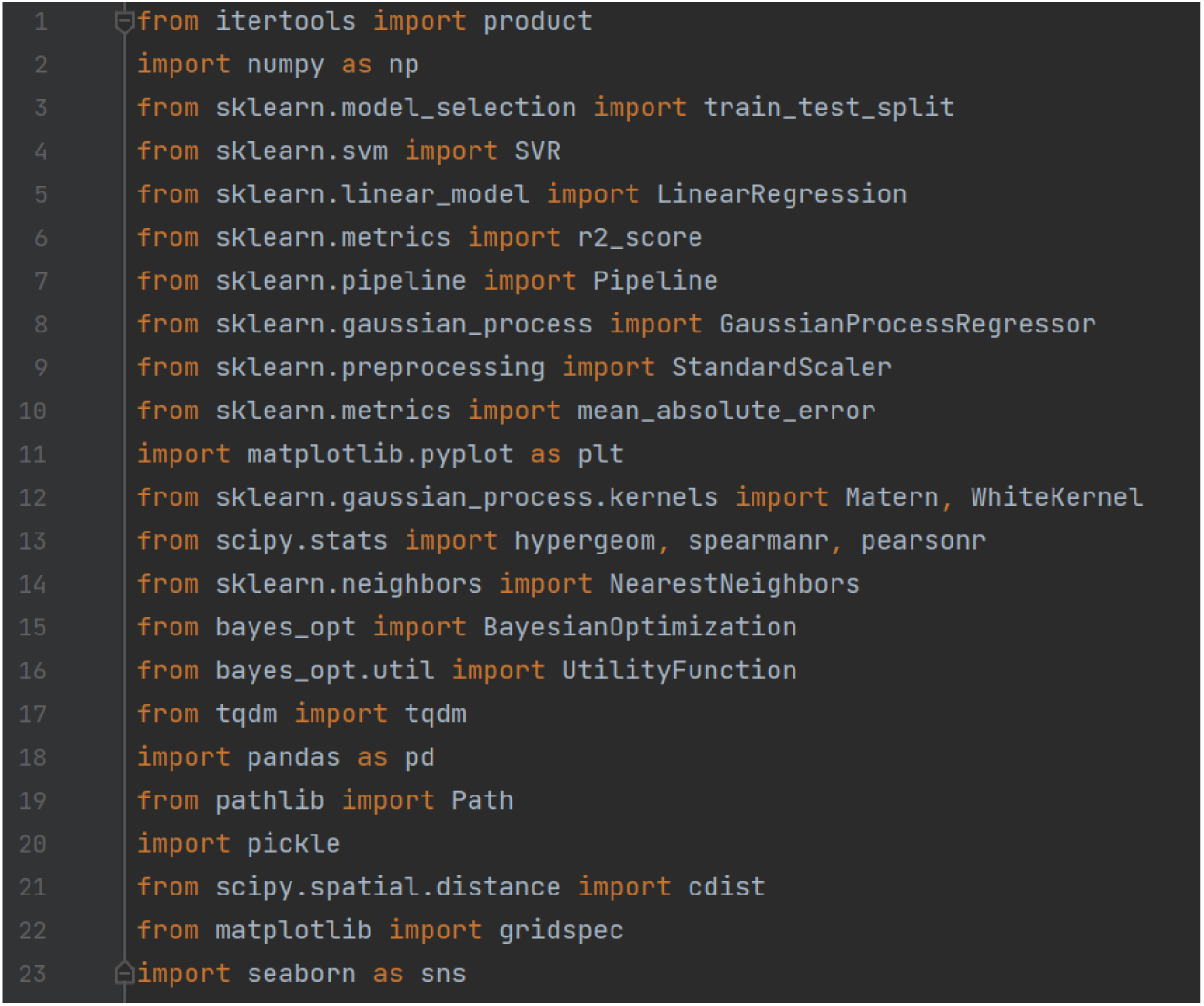

a. sklearn is used for machine learning tasks, including data splitting, egression models, and evaluation metrics; the kernel definitions for Gaussian Process regression, and for statistical tests and correlation metrics;
b. Sklearn.neighbors and scipy.spatial are used for identifying the nearest points in the model embedding space;
c. bayesopt libraries implements the core Bayesian Optimization process functions;
d. numpy, pandas and pickle libraries assist in organizing, reading and manipulating data;
e. pathlib manages file paths;
f. matplotlib and seaborn libraries are used to create visual plots and graphs;
g. tqdm creates progress bars for loops;
h. itertools generates the Cartesian product of multiple input sequences for grid construction.

### Preparation

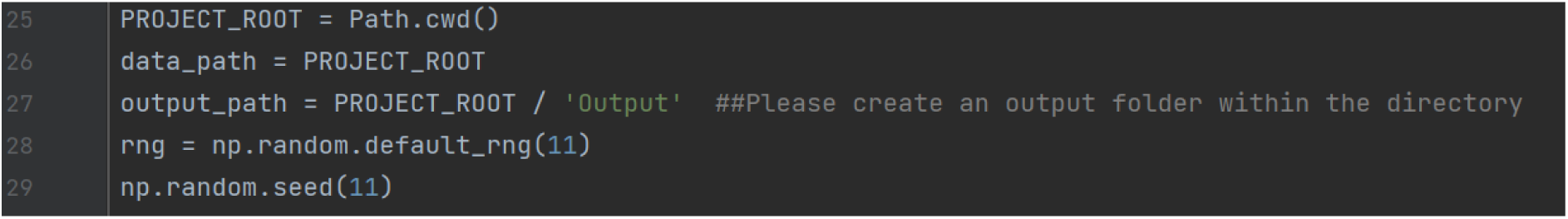

a. PROJECT_ROOT defines the root directory for the project;
b. data_path and output_path are set for organizing input data and outputs;
c. The random seed ensures reproducibility in random number generation.

### Function objective_func_reg

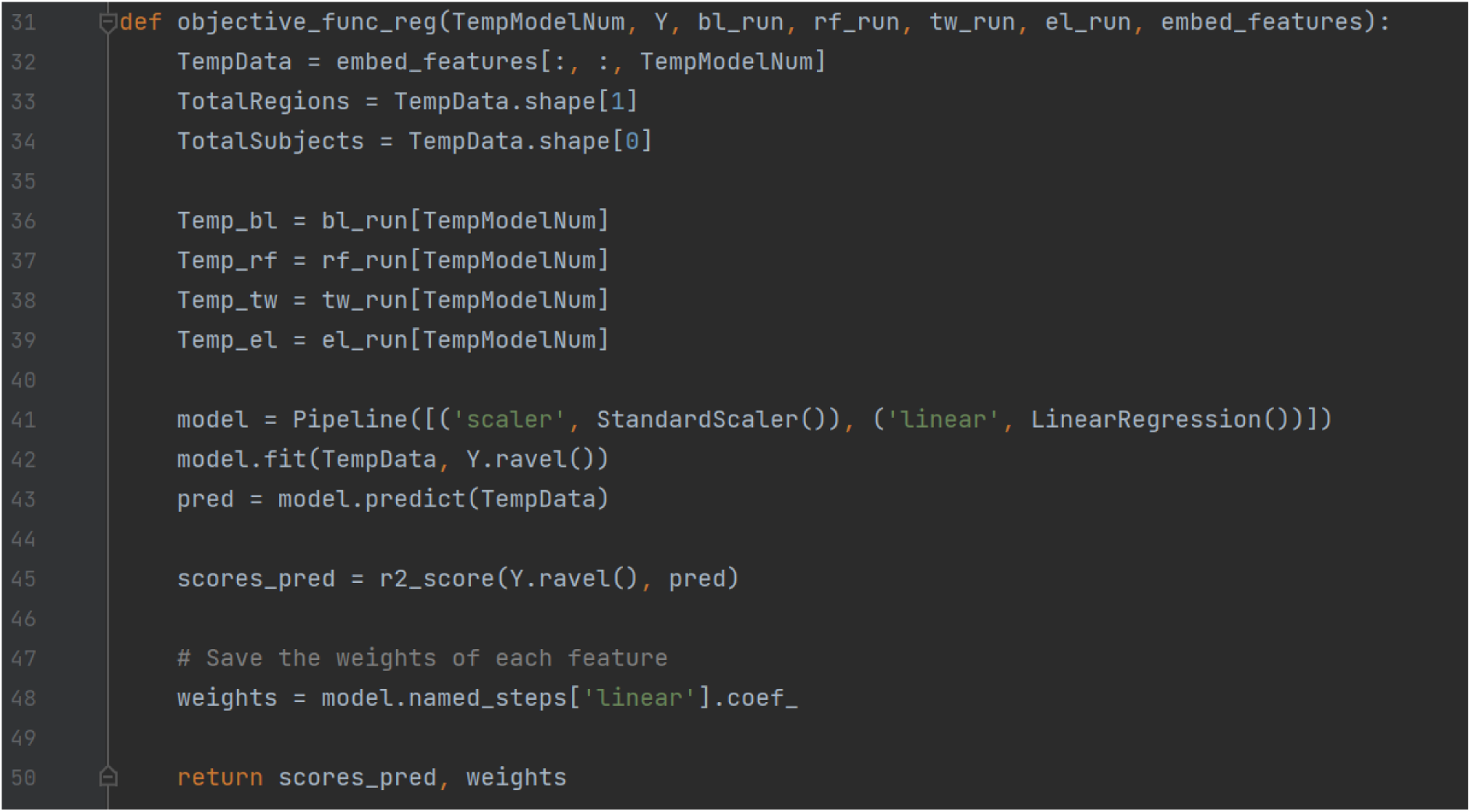

This function evaluates the performance of a pipeline model by fitting a linear regression to the feature data and computing the *R*^2^.

<u>Inputs:</u>

a. TempModelNum – index of the model to evaluate;
b. Y – target values;
c. bl_run, rf_run, tw_run, el_run – parameters corresponding to decision nodes;
d. embed_features – feature embedding data for models.

<u>Steps:</u>

a. Extracts features for a specific model;
b. Constructs and trains a linear regression model;
c. Computes the *R*^2^ to evaluate the model’s performance;
d. Returns the *R*^2^ and feature weights.

### Function posterior

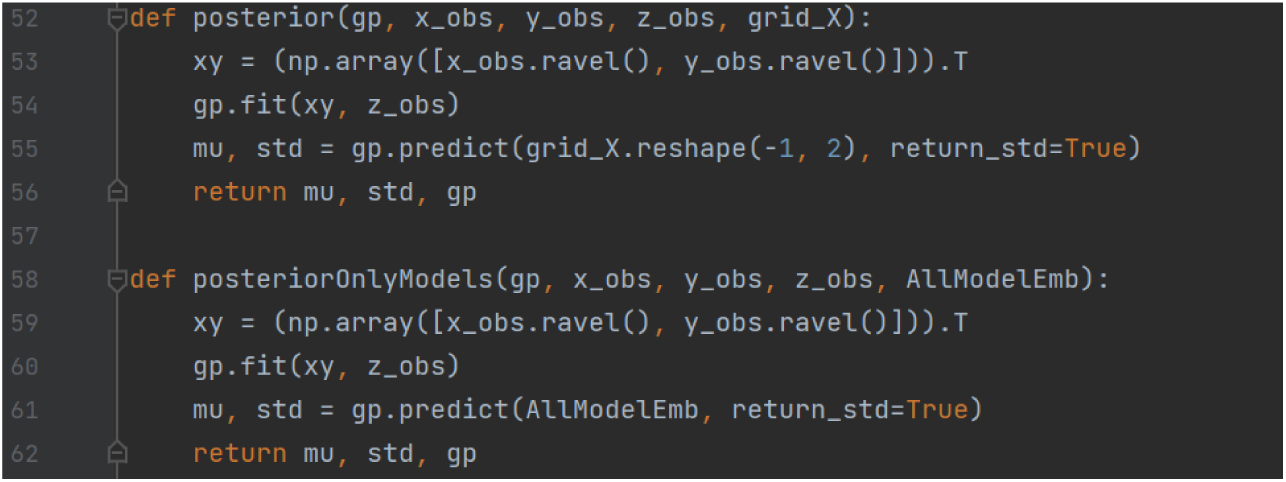

Computes the posterior mean and standard deviation for Gaussian Process regression.

Fits the Gaussian Process to observed data points (x_obs, y_obs, z_obs), which dynamically expands as more points are sampled, and predicts the posterior distribution for new points (grid_x).

### Function initialize_bo

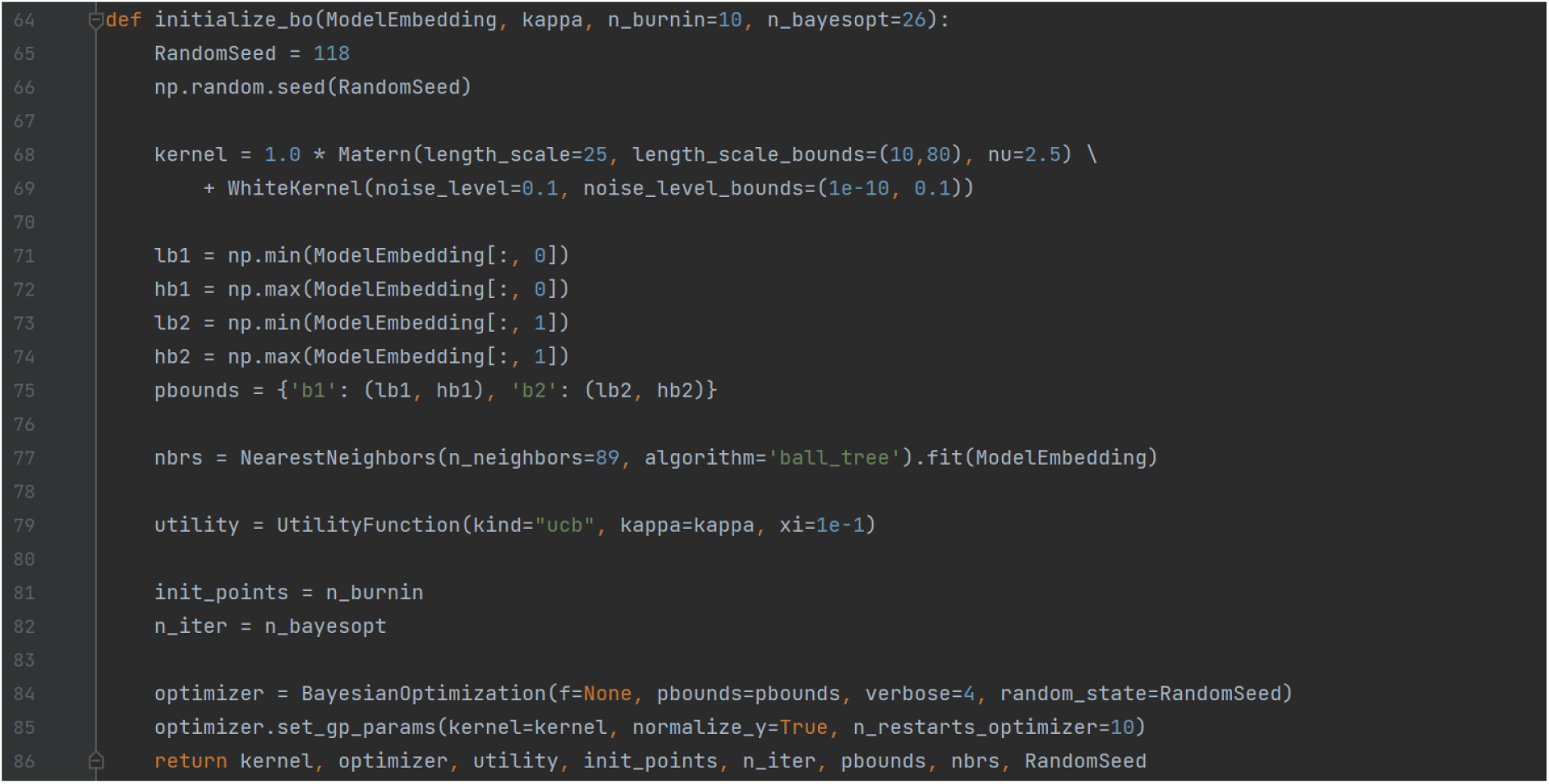

This function sets up the Bayesian optimization:

a. Defines a kernel combining a Matern kernel (spatial covariance) and noise;
b. Configures bounds for optimization and initializes the optimizer and utility function;
c. Uses NearestNeighbors for model selection based on proximity.

### Function run_bo

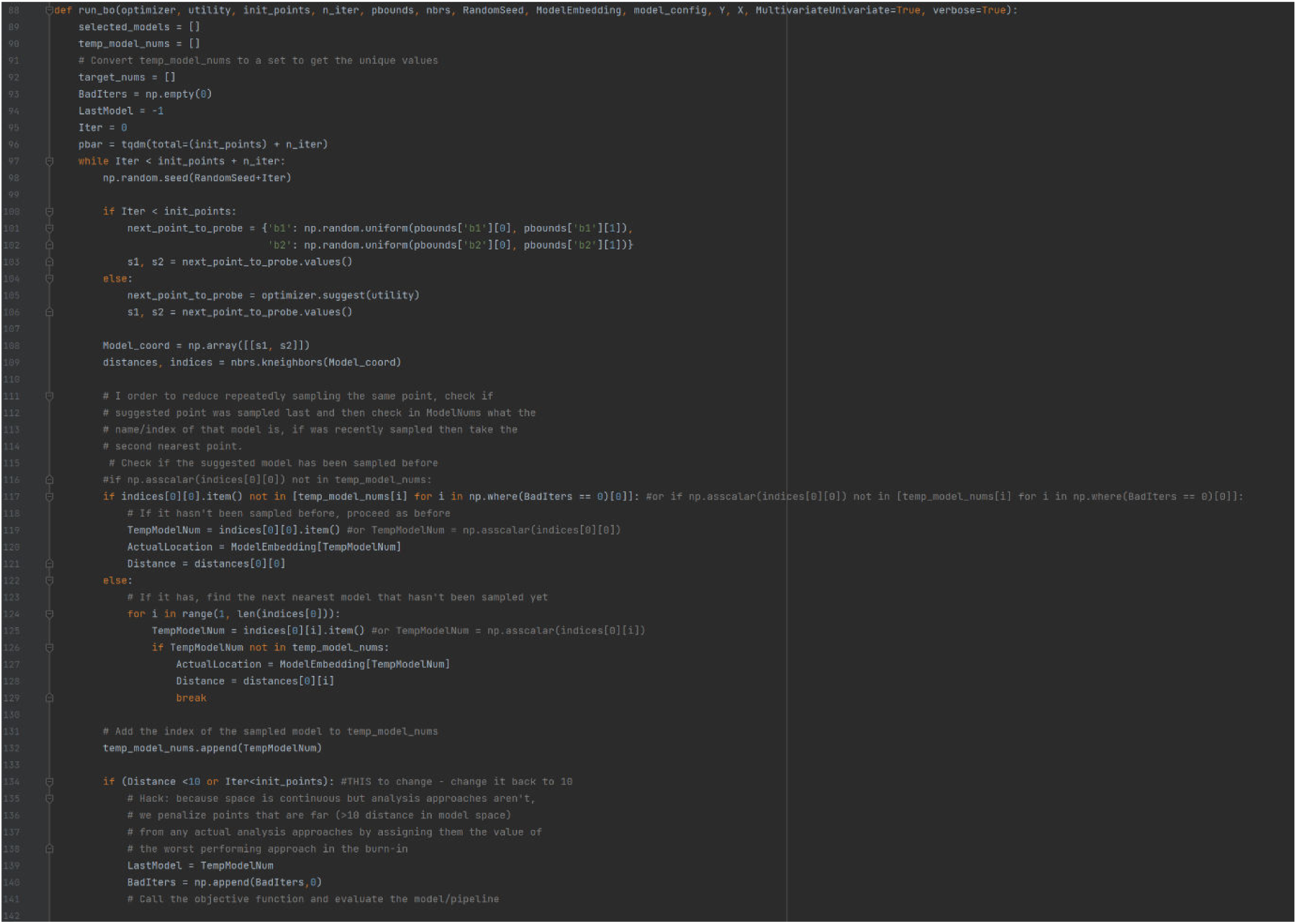

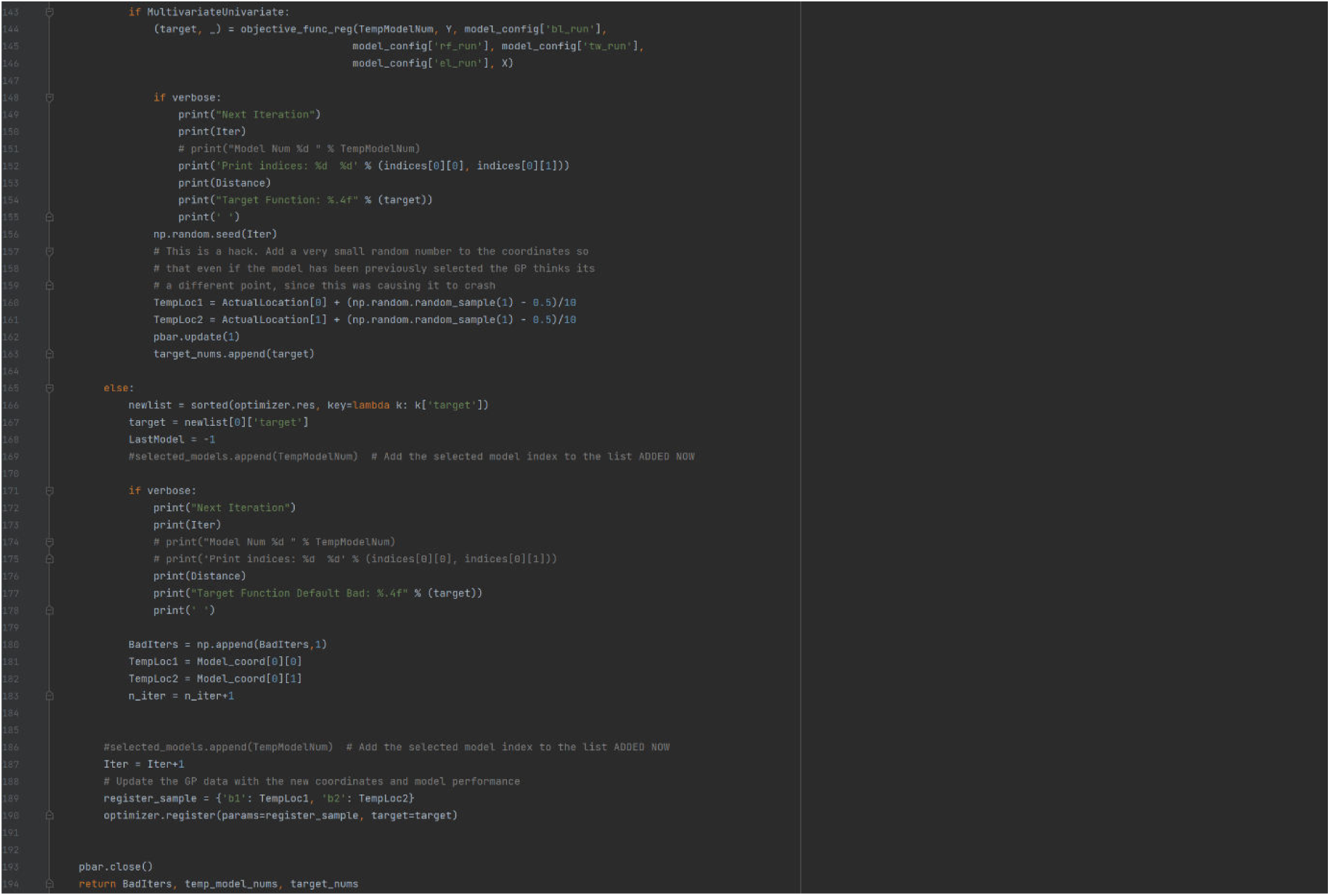

Implements Bayesian Optimization for active learning:

a. Randomly samples points (init_points) during a “burn-in” phase;
b. Suggests new points using the utility function;
c. Evaluates the objective function for the sampled models and updates the optimizer;
d. Penalizes models that were recently sampled or far from existing points to avoid repeated sampling and maintain exploration;
e. Tracks bad iterations, selected models, and target scores;
f. The use of np.random.seed(RandomSeed + Iter) ensures reproducibility for each iteration.

### Function plot_bo_estimated_space

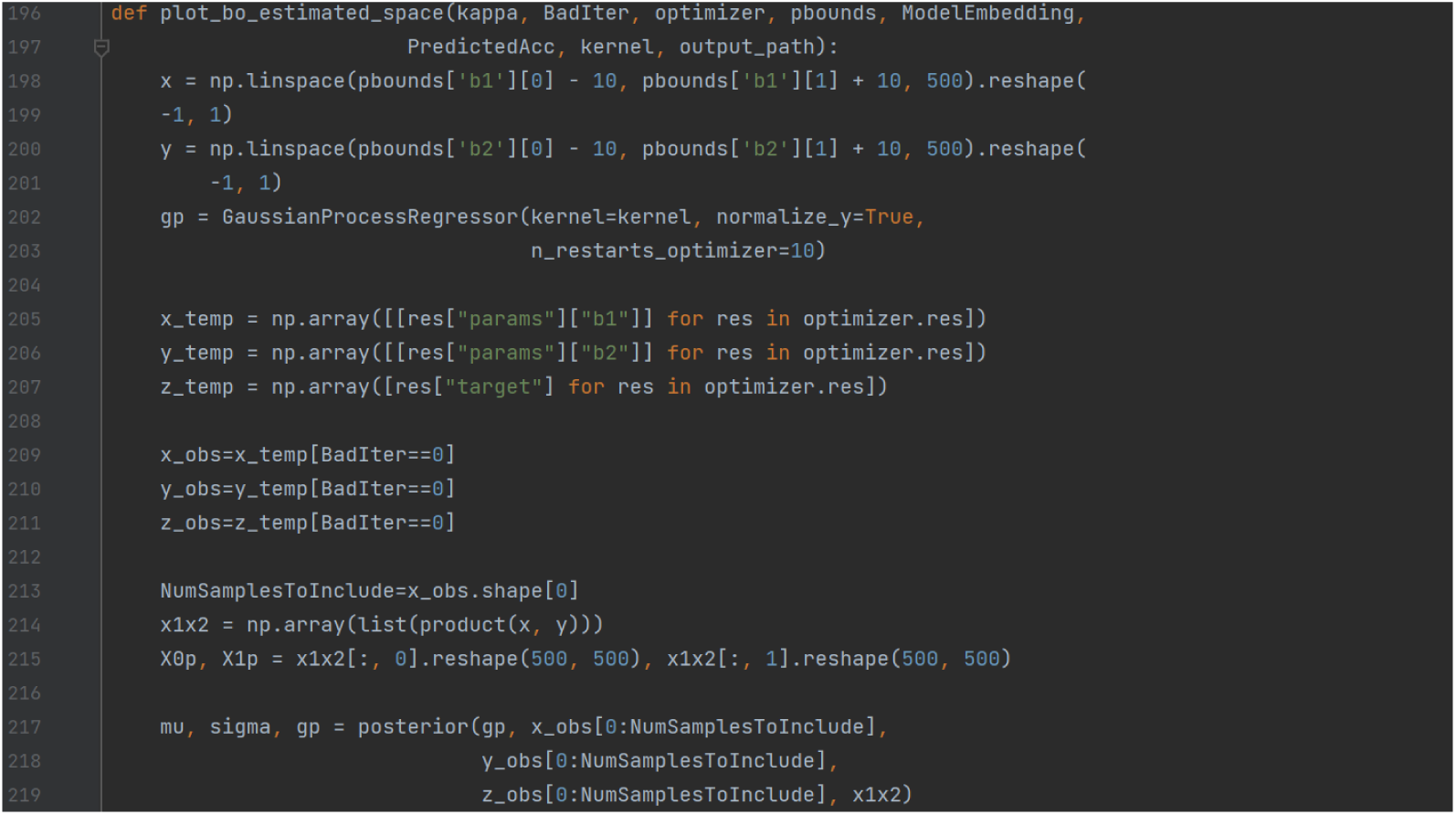

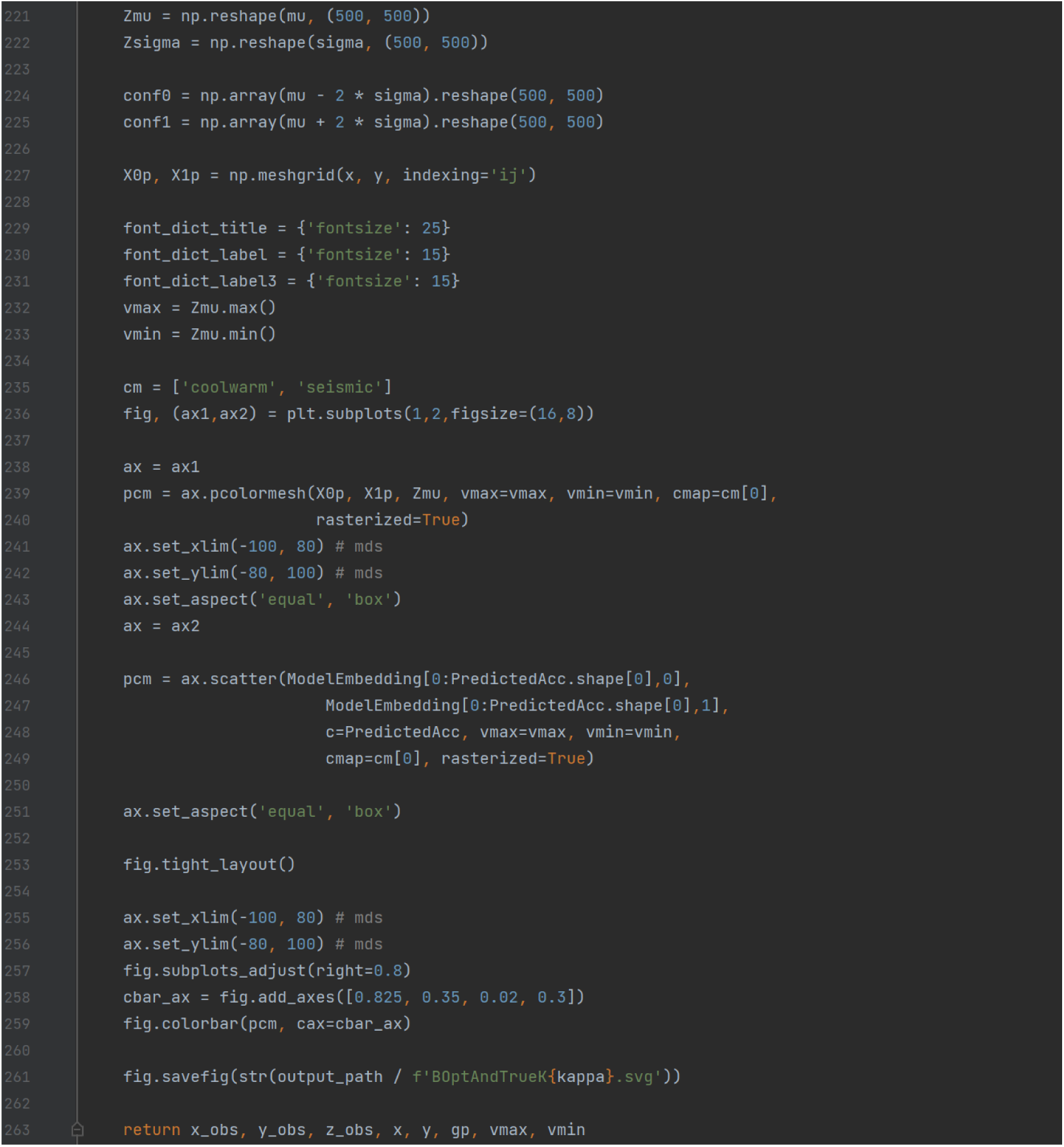

Visualizes the Bayesian Optimization process:

a. Plots the posterior mean and uncertainty of the search space;
b. Highlights the locations of sampled models.

Specifically,

- The np.linspace funtion creates evenly spaced grid points in the parameter space, defined by pbounds (the boundaries for b1 and b2). The bounds are expanded slightly (−10, +10) to provide better visualization at the edges;
- The GuassianProcessRegressor() initialized the Gaussian Process using the provided kernel (which includes spatial covariance and noise). The Gaussian Process will fit the observed data and predict the posterior mean and uncertainty across the grid;
- The x/y/z_temp and x/y/z_obs extract observed points (b1, b2) and their corresponding target values from the Bayesian optimizer’s history. BadIter == 0 excludes points flagged as bad iterations as defined above;
- x1×2 and X0p, X1p construct all combinations of grid points (x,y) using a Cartesian product, resulting in a 2D grid of shape (500,500);
- mu, sigma, gp calls the posterior function to fit the Gaussian Process to observed data, predicts the posterior mean and uncertainty over the grid, and reshapes predictions (Zmu, Zsigma) into 2D arrays for plotting;
- conf0 and conf1 compute 95% confidence intervals using +/− 2 SDs from the mean;
- Then plot parameters like font size, color map and value range are configured;
- Then the posterior mean and the predictive accuracies are plotted.

### Function plot_bo_evolution

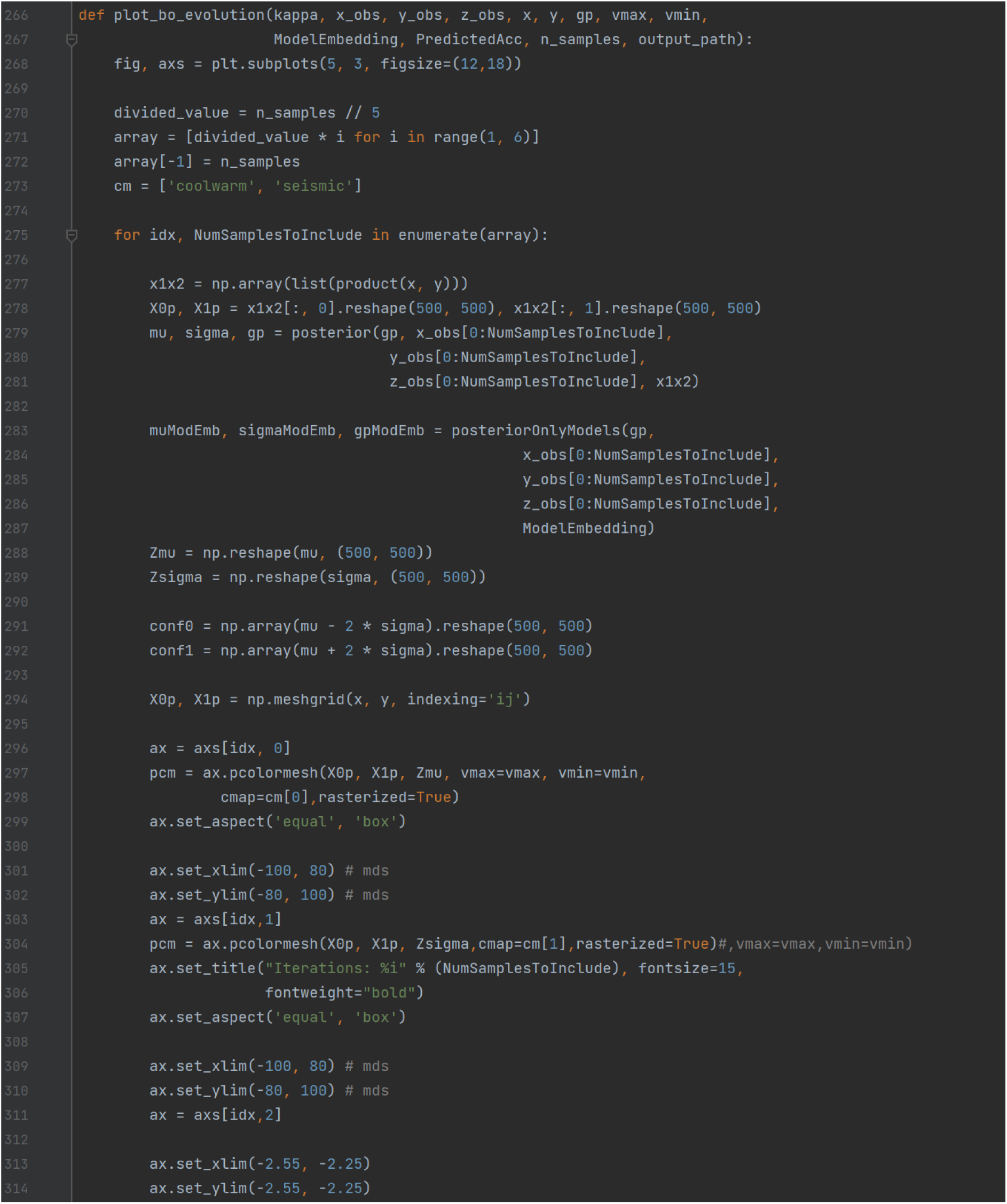

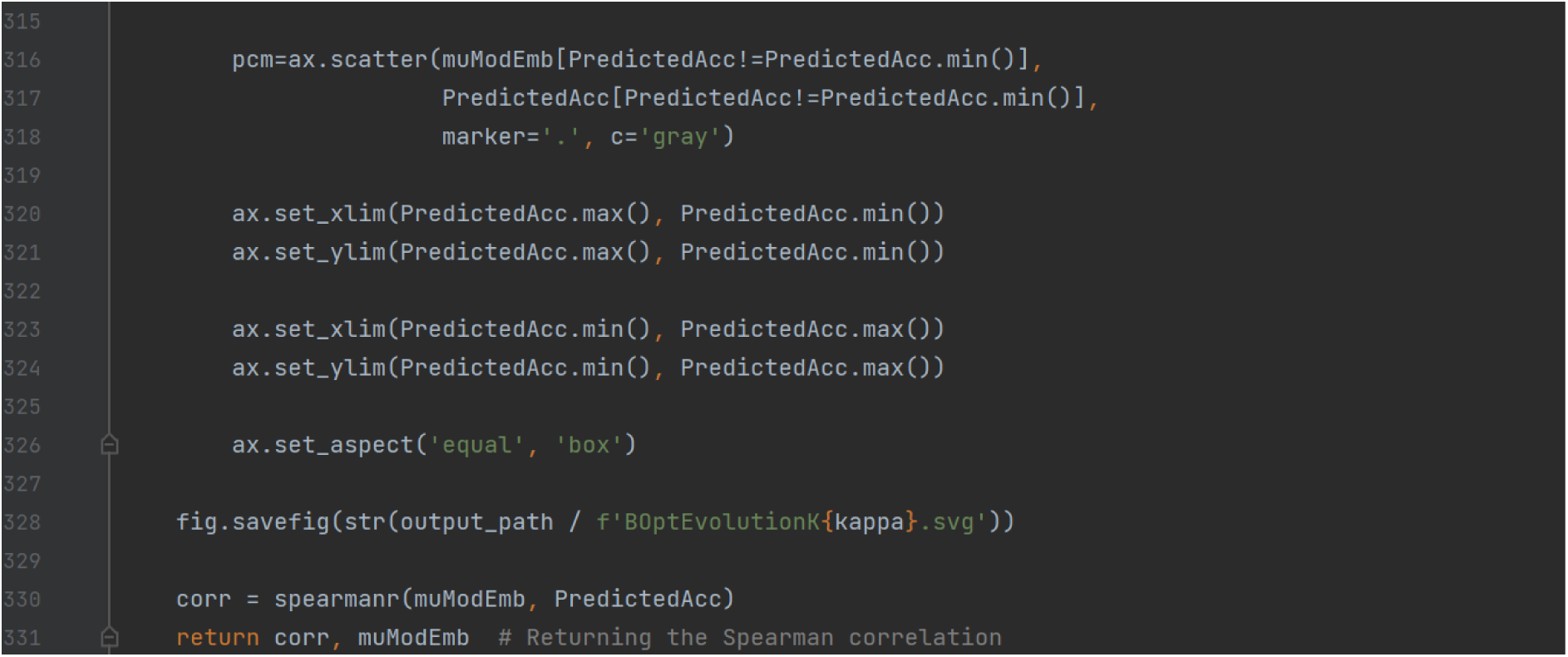

This function visualizes the evolution of the Bayesian optimization across iterations by:

1. Showing the Gaussian Process posterior predictions (mean and uncertainty) change as more observations are added.

a. Creates a grid of 5 rows and 3 columns for plotting (3 columns are for the posterior mean, the posterior uncertainty, and the scatterplot for the predicted vs actual performance of the models;
b. Divides the total number of samples (n_samples) into 5 equal segments, to represent snapshots of the optimization process at different stages, and loops through these defined intervals to visualize the optimization at each stage;
c. Creates a grid of points over the search space using the Cartesian product (x, y), which are used for the Gaussian Process predictions;
d. Fits the Gaussian Process on the observed data (x_obs, y_obs, z_obs) up to the current iteration, and predicts the posterior mean and uncertainty over the grid;
e. Fits a Gaussian Process for the model embeddings (ModelEmbedding), and predicts the mean (muModEmb) and uncertainty (sigmaModEmb) for these embeddings;
f. Plots the posterior mean (Zmu) and the posterior uncertainty (Zsigma) as heatmaps over the search space;
g. Plots the predicted means (muModEmb) against the accuracies (PredictedAcc) for the model embeddings to compare the Gaussian Processes’ predictions with the actual performance of sampled models, and saves as an SVG file.
2. Calculating the Spearman correlation between predicted performance and actual performance of models.

a. Computes the Spearman correlation between the Gaussian Process’ predicted means (muModEmb) and the actual accuracies (PredictedAcc);
b. Returns the correlation and predicted means for further analysis.

In summary, this function tracks the Bayesian optimization progress through visualization, provides three plots per stage (the posterior mean, the posterior uncertainty, and the predicted vs. actual performance), saves the evolution plot which visualizes the optimization process, and evaluates the Gaussian Process accuracy by returning the Spearman correlation between the predictions and actual performance.

### Function create_unique_balanced_stratified_sample

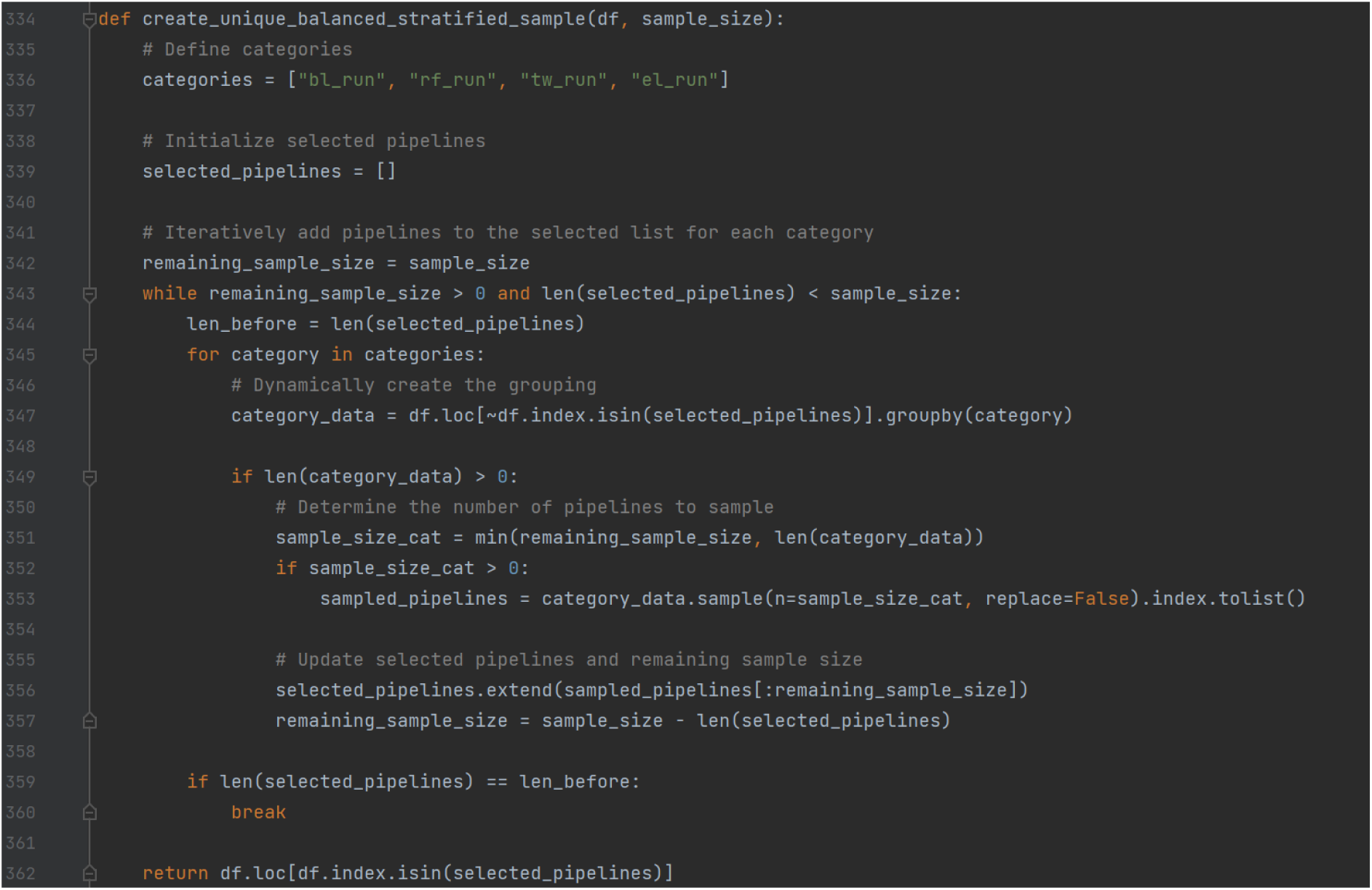

This function generates a stratified sample that balances representation of options within each decision node.

a. Defines categories (columns in the dataframe, which are the decision node categories);
b. Initializes the selected_pipelines variable, which keeps track of the indices of rows that have already been selected, and the remaining_sample_size variable, which tracks how many more samples are needed to reach the target sample_size;
c. Engages in a while loop until the target sample_size is reached, by iterating until remaining_sample_size becomes 0, or the selected_pipelines list reaches the desired sample_size;
d. Groups the remaining data by each category by filtering the dataframe to exclude rows that are already selected and grouping the remaining rows by category;
e. Samples from each group by determining how many rows to sample within each group and randomly sampling within that group without replacement;
f. Updates the selected pipelines by adding the sampled rows to the selected_pipelines list and updating the remaining_sample_size based on the number of rows added;
g. Considering an exit condition, which exits the loop early if no new rows are added during the current iteration to prevent an infinite loop in cases where there is insufficient data;
h. Returns the stratified sample containing only the rows corresponding to the selected pipelines.

If you do not wish for a balanced representation of options within a specific decision node, for example if this is not representative of your multiverse because there is a bias within the larger multiverse of defensible pipelines, the function should be amended accordingly.

### Function create_stratified_samples

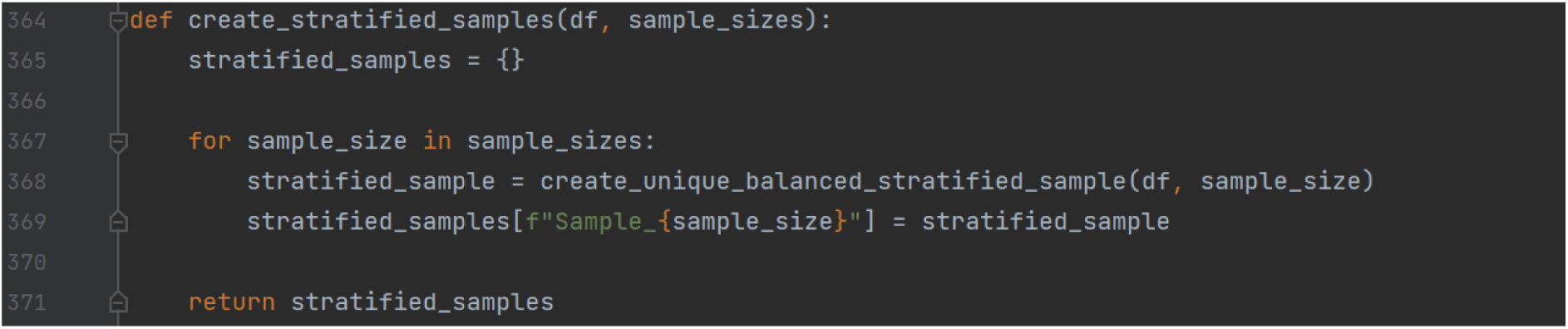

This function uses the create_unique_balanced_stratified_sample to generate multiple stratified samples of specific sizes from the dataset.

a. Initializes a dictionary (stratified_samples) to store the stratified samples. Each key in this dictionary will represent a sample size, and each key corresponds to the stratified sample of that size;
b. Loops through the list of desired sample sizes;
c. For each sample size, generates a stratified sample using the create_unique_balanced_stratified_sample function, the current sample_size, and the dataframe, and stores it in the stratified_samples dictionary where all samples are organized and accessible by their size;
d. Returns the dictionary containing all of the stratified samples.

### Function average_nearest_neighbor_distance

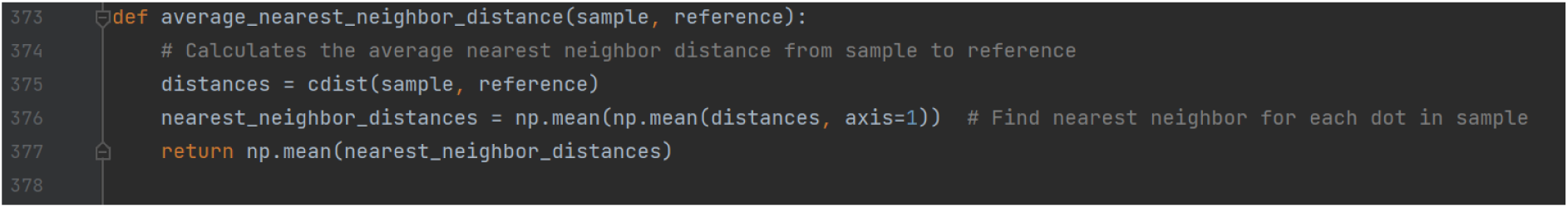

This function calculates the average nearest neighbor distance between points in the sample and points in the reference:

a. Computes pairwise distances – the Euclidean distance between every point in the sample and every point in the reference. Distances is a two-dimensional array of shape (n_sample, n_reference), where each element represents the distance from a point in the sample to a point in the reference;
b. Calculates the nearest neighbor distance by computing the mean of the distances for each point in the sample across all points in the reference and taking the overall mean of these distances to get a single scalar value;
c. Returns the computed average nearest neighbor distance as a scalar to evaluate the proximity between the two sets of points.

### Function spec_curve

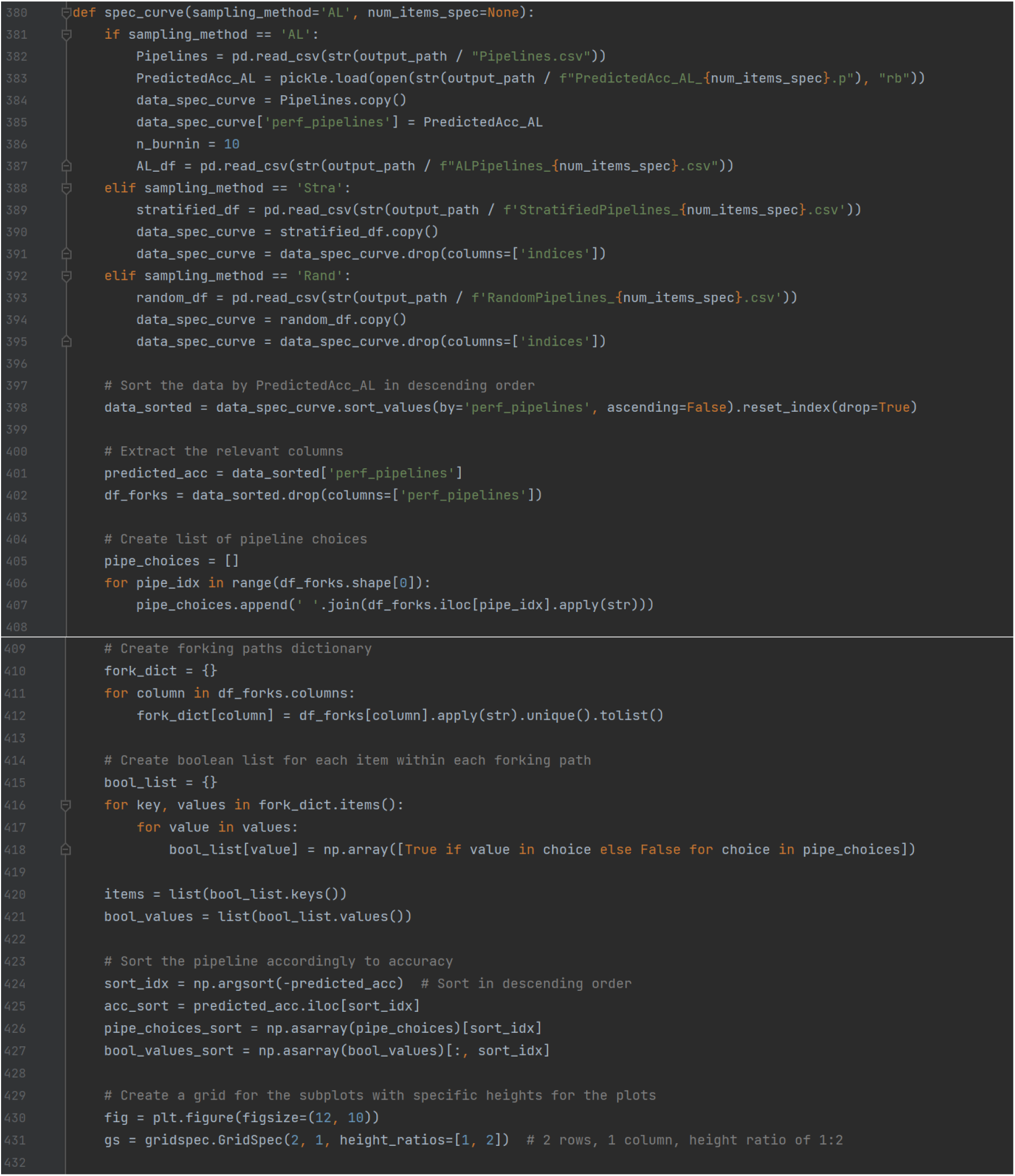

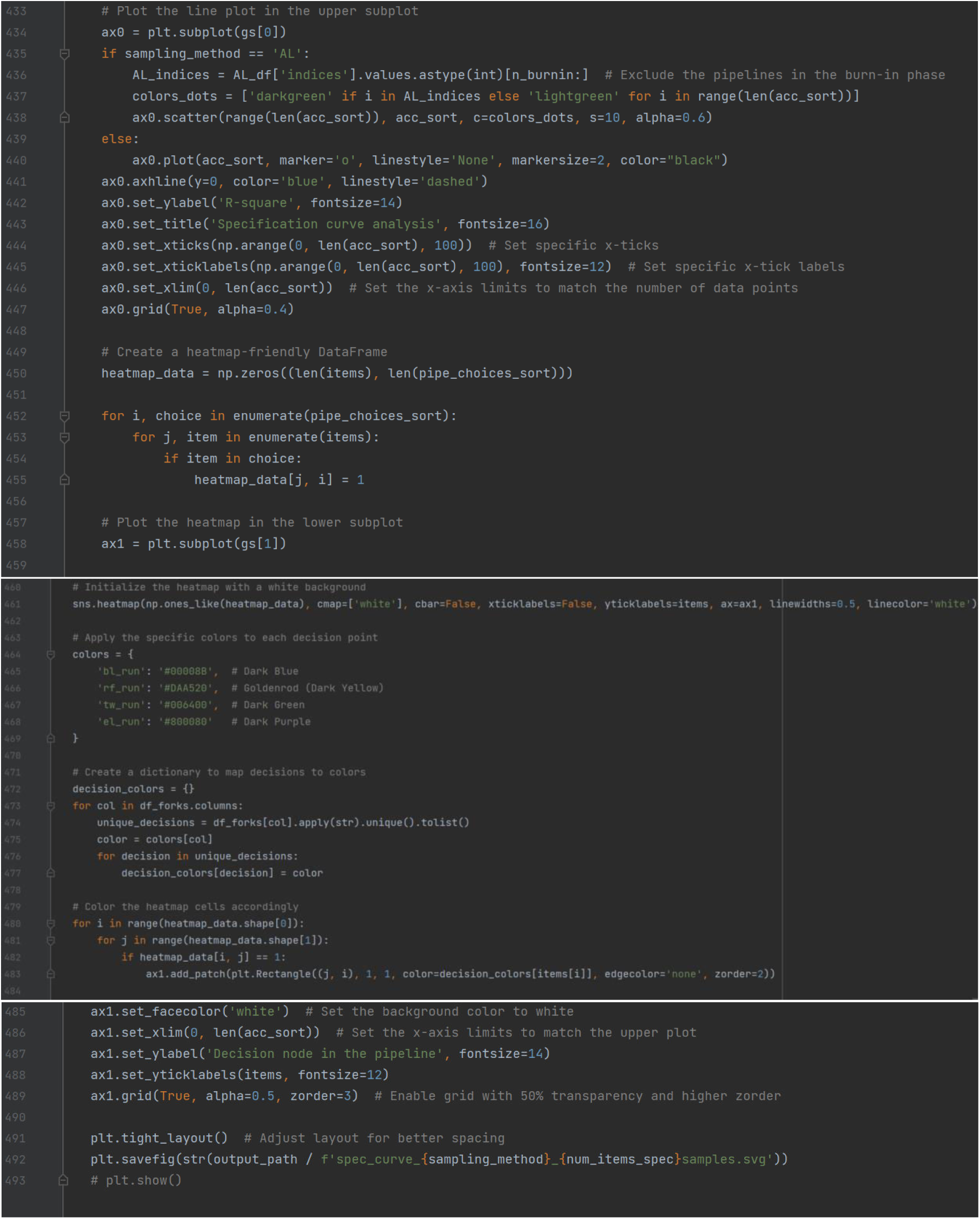

This function creates a specification curve figure, which consists of the upper panel that visualizes the performance of each pipeline and the lower panel that visualizes the options at each decision node that contribute to the respective pipelines of each result plotted in the upper panel.

a. Reads and processes pipeline performance data depending on the sampling_method (AL, Stra, or Rand), and combines the pipeline decisions with performance metrics (perf_pipelines) into a single dataframe. For AL (the active learning sample), it excludes the burn-in iterations (n_burnin);
b. Sorts pipelines based on performance (descending order), and extracts decision paths and their unique combinations;
c. Creates a dictionary where each key is a decision node and the values are a list of unique options for that decision node;
d. Converts each option value into a Boolean array indicating whether that decision is present in each pipeline configuration;
e. Visualizes the results by plotting the upper and lower panels as described above. For AL (the active learning sample), the points in the upper panel are color coded to distinguish the pipelines that are sampled and the rest that are estimated;
f. Saves the visualization as an SVG file.

## Supplementary 3 Grand Average Waveforms

Visualising the grand average waveforms for each experimental condition across the alternative reference schemes and electrodes of interest. Baseline = −200ms. *N* = 98 participants.

The default amplitude range on the y axis is −6 to 10 µV. This differs only for panels marked by ‘**\***’. In these instances, the y axis range exceeded the default, so it is determined by the minimum and maximum µV value of that waveform.

The electrode option of 4 electrodes around the midline peak was not included in the figure as the resulting electrodes were identical to the electrode cluster CP1, CP2, P3, P4, Pz that is already included.

The figure is also uploaded as an SVG file with high resolution, titled: GrandAvergeWaveforms.svg

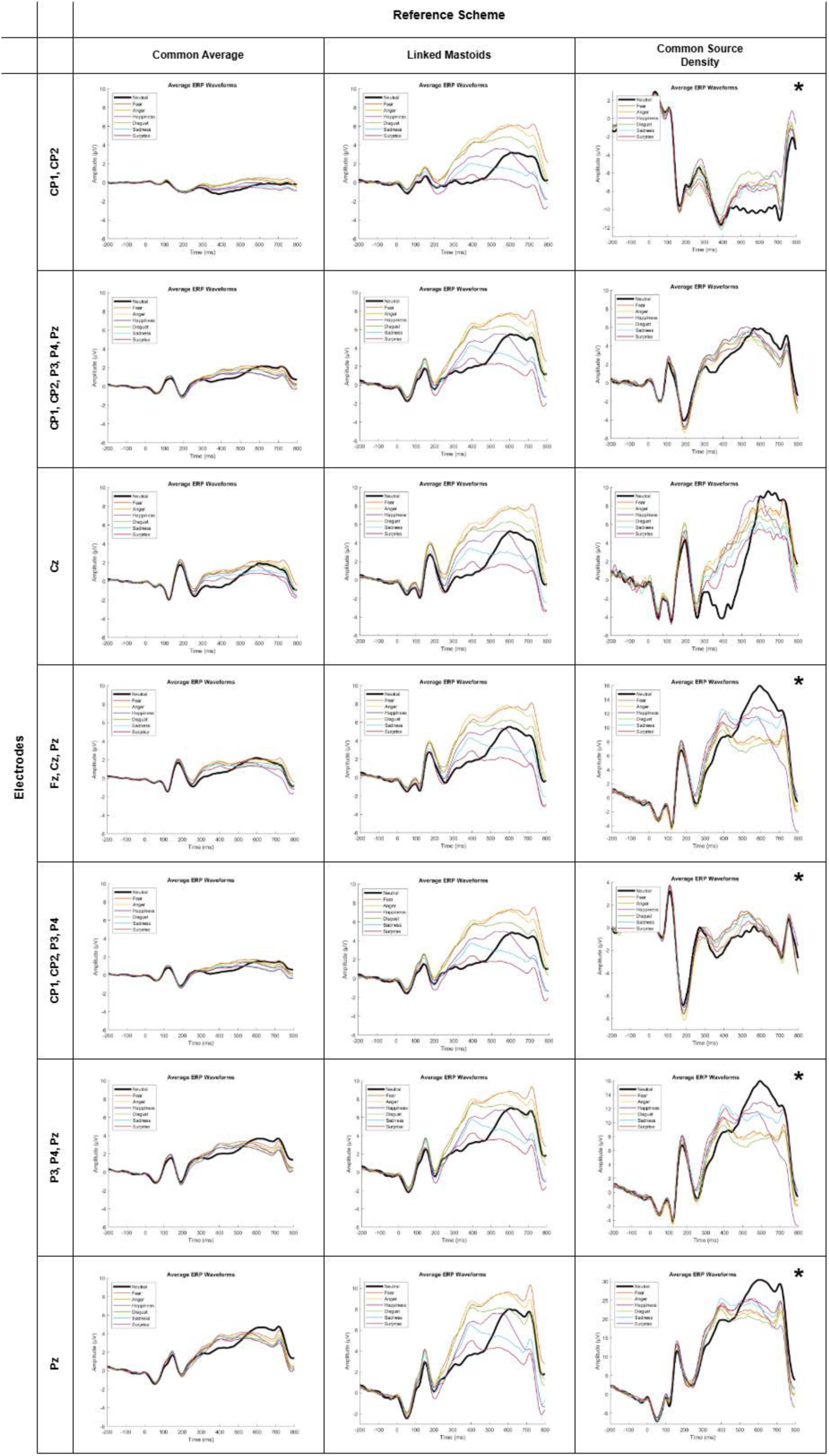

## Supplementary 4: The alternative embedding methods tested

**Figure 1:**
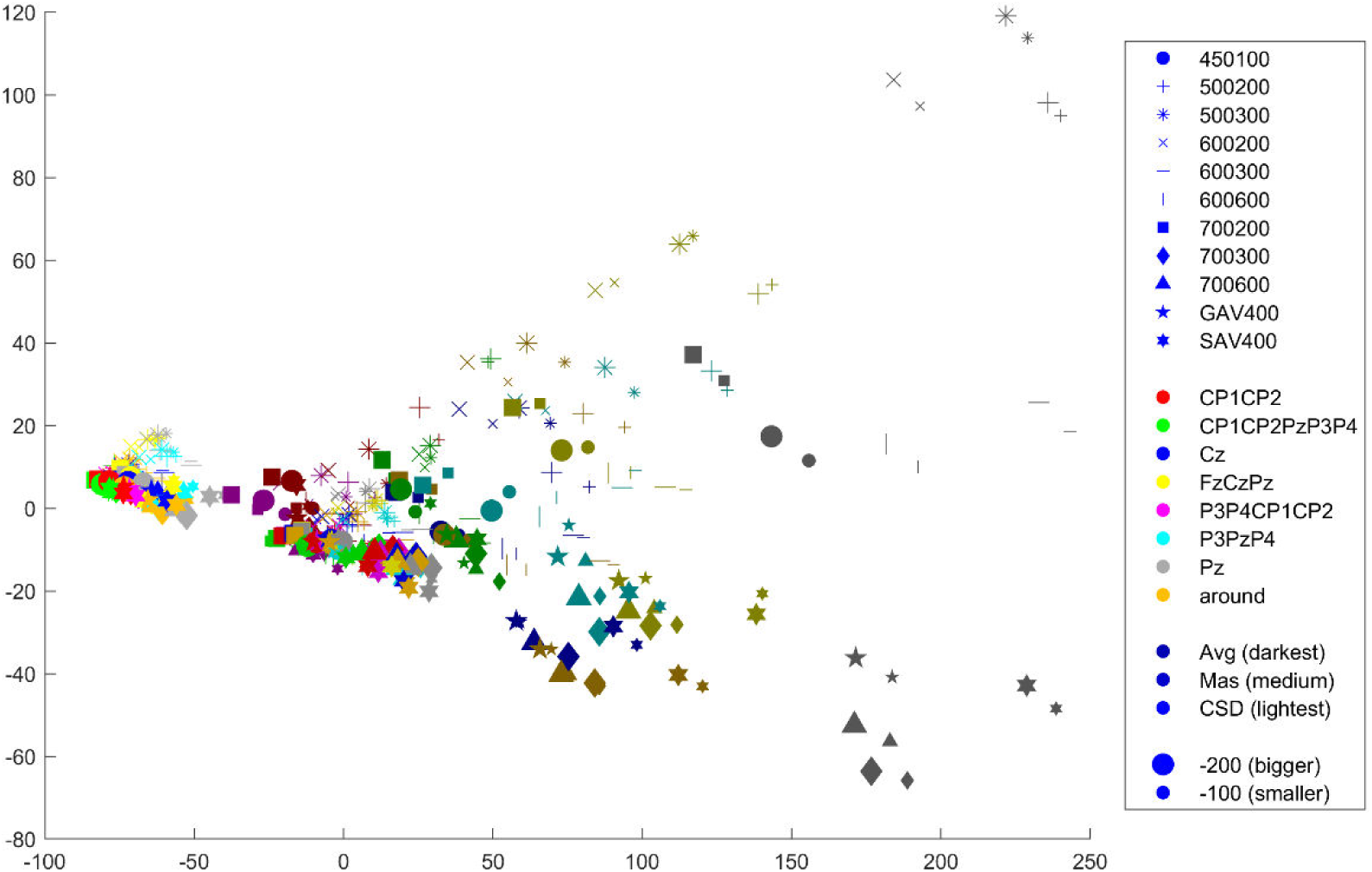
Classical multidimensional scaling (classical MDS)

**Figure 2:**
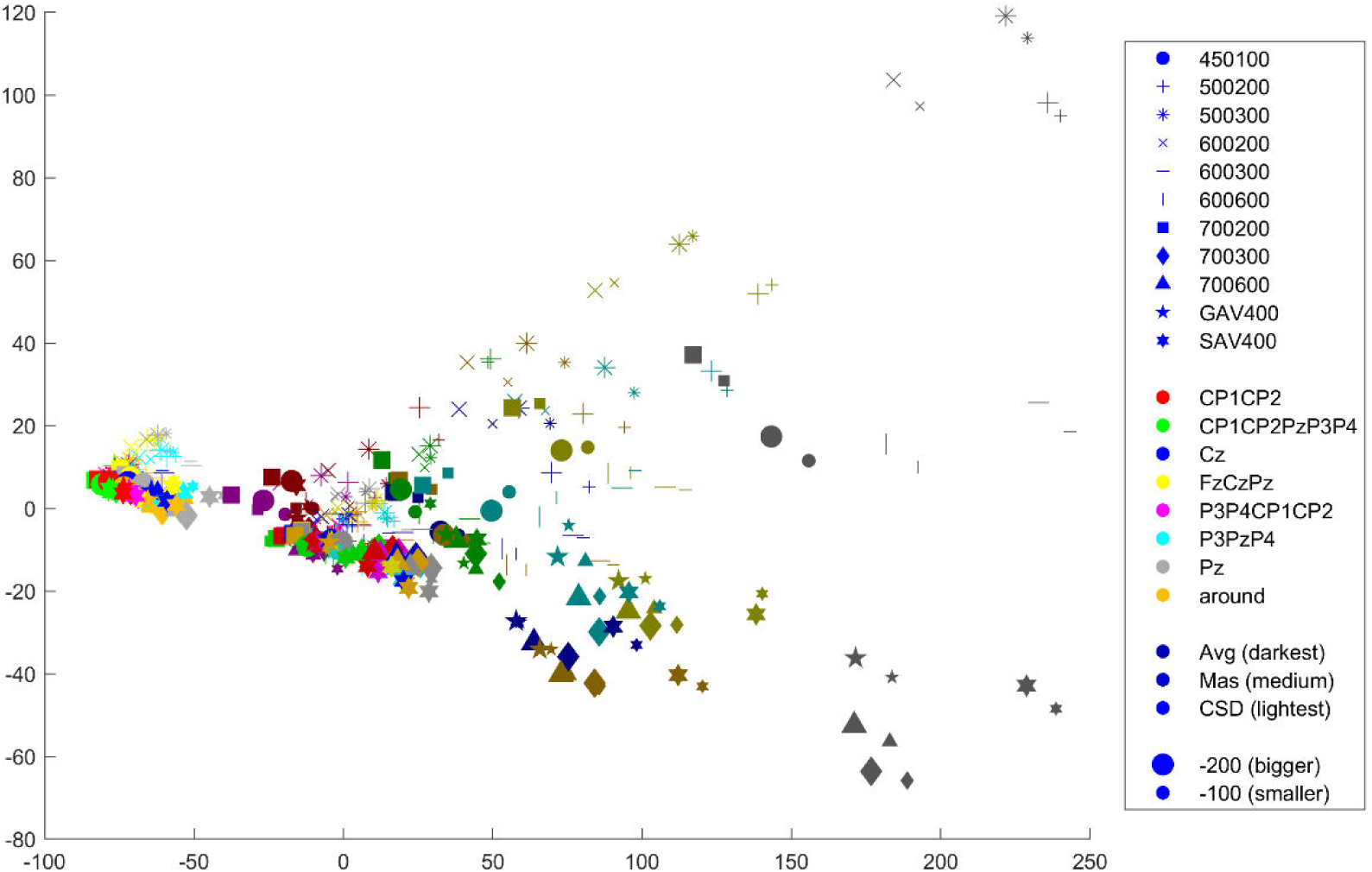
Principal Component Analysis (PCA)

**Figure 3:**
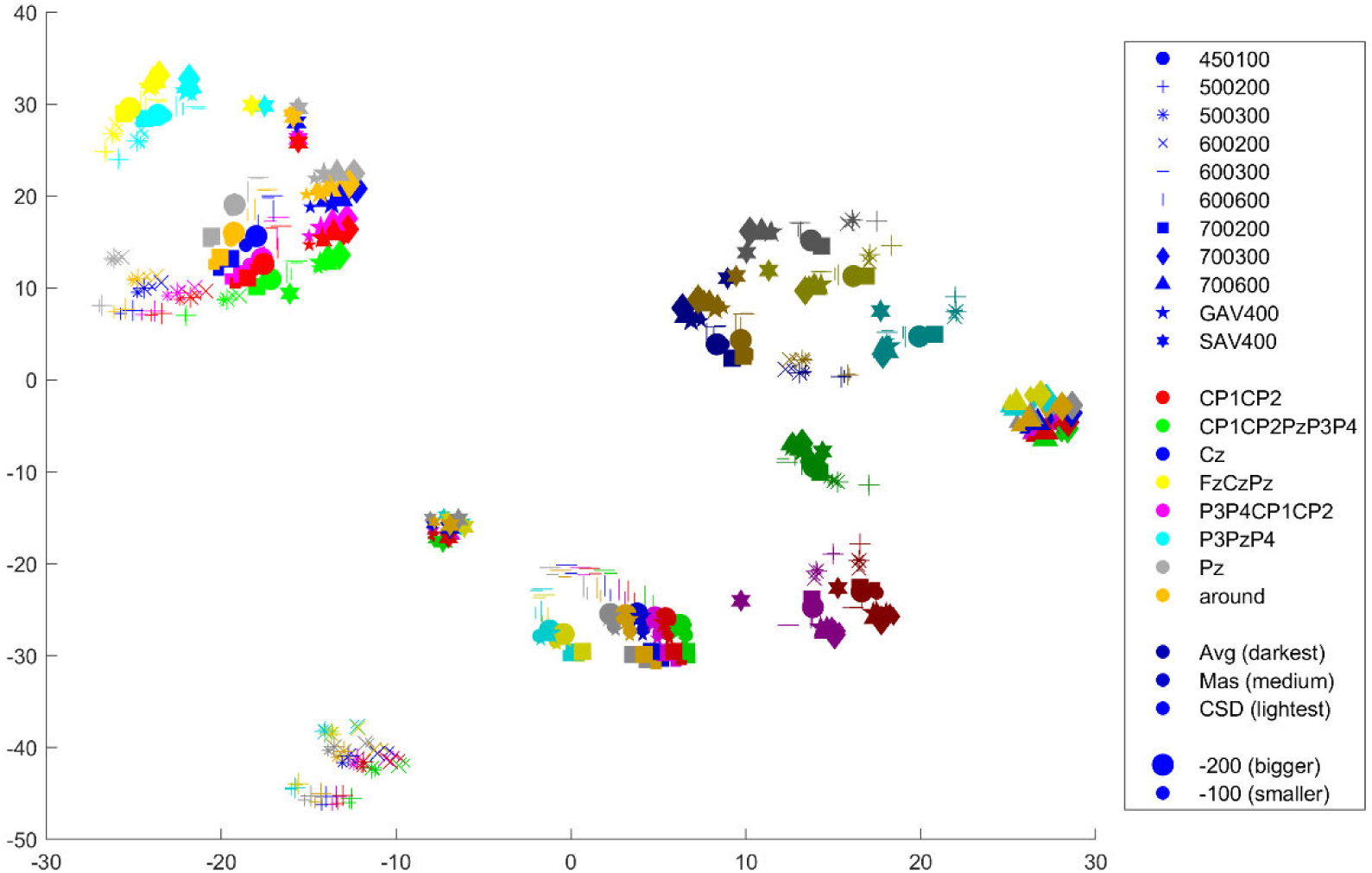
*t*-distributed Stochastic Neighbor Embedding (t-SNE), perplexity 30

**Figure 4:**
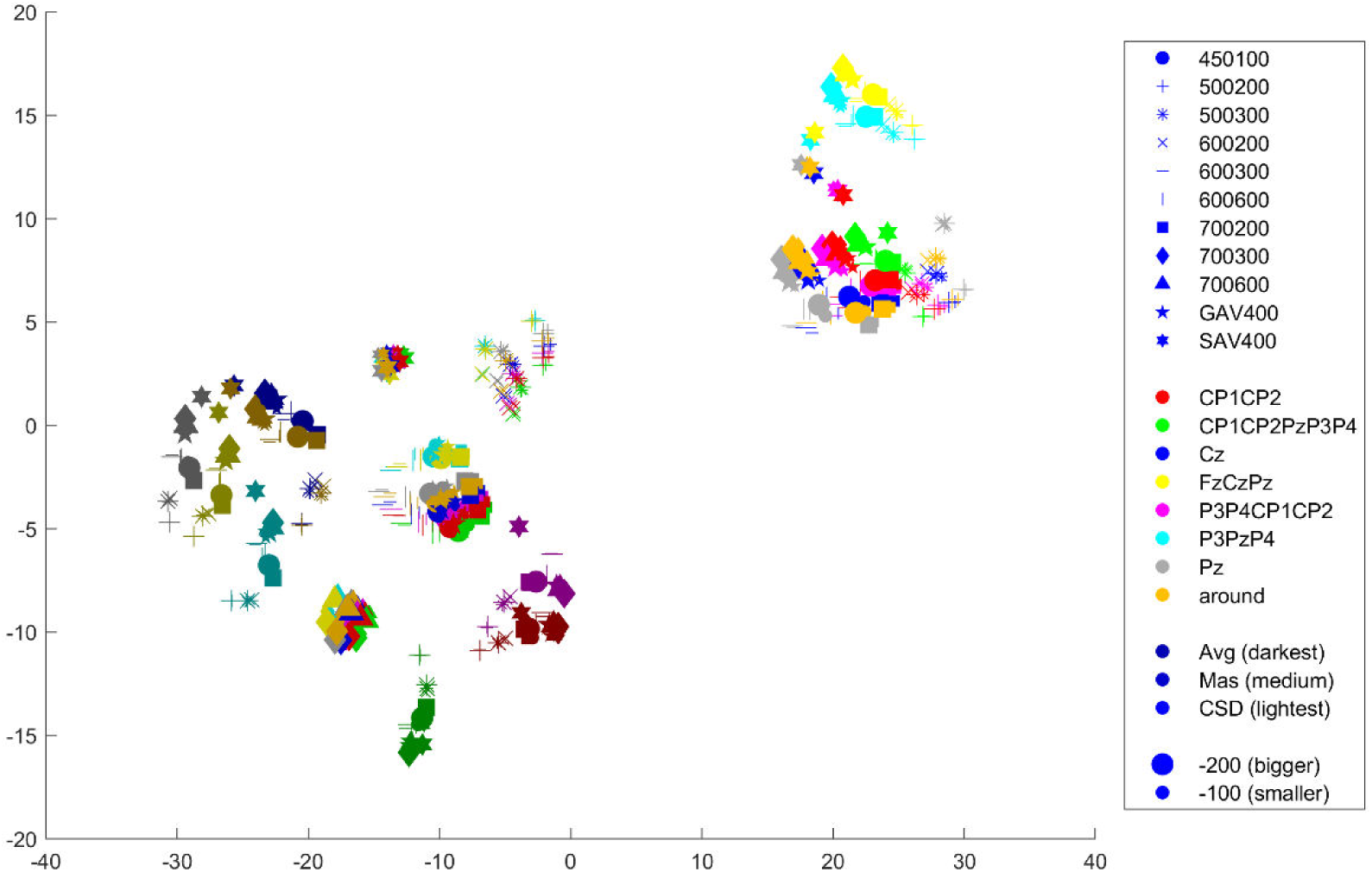
*t*-SNE, perplexity 50

**Figure 5:**
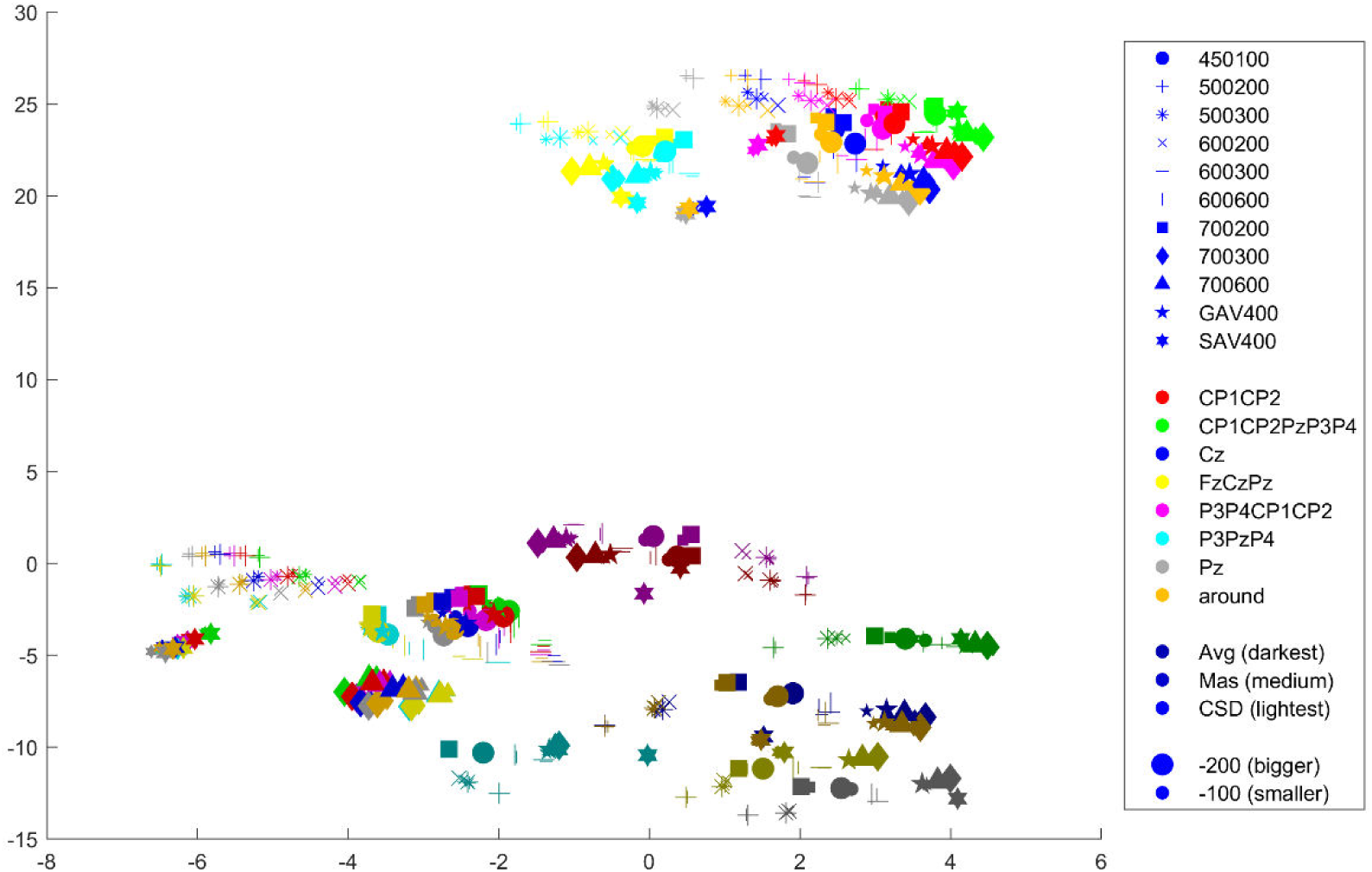
*t*-SNE, perplexity 100

**Figure 6:**
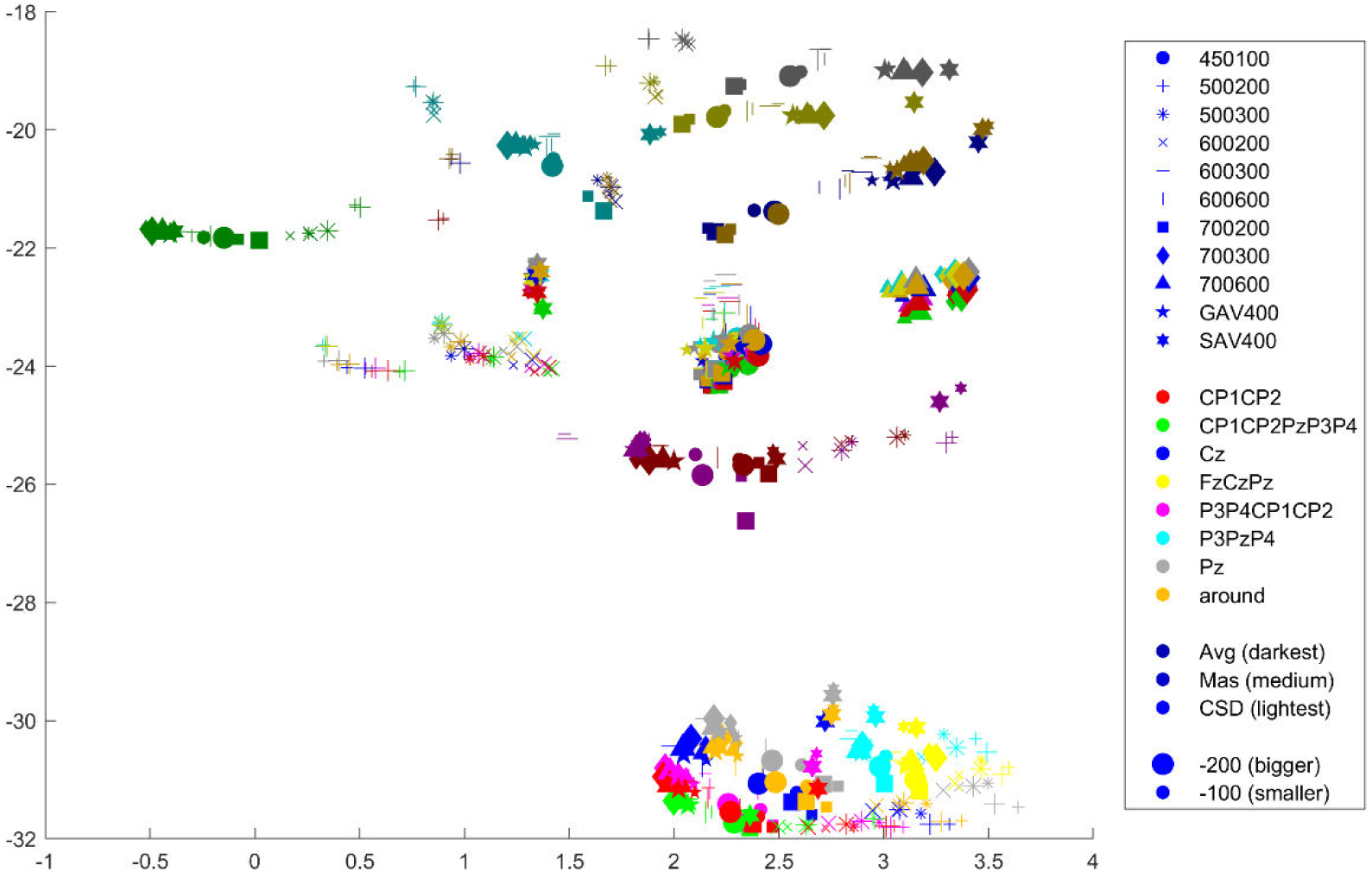
*t*-SNE, perplexity 190

**Figure 7:**
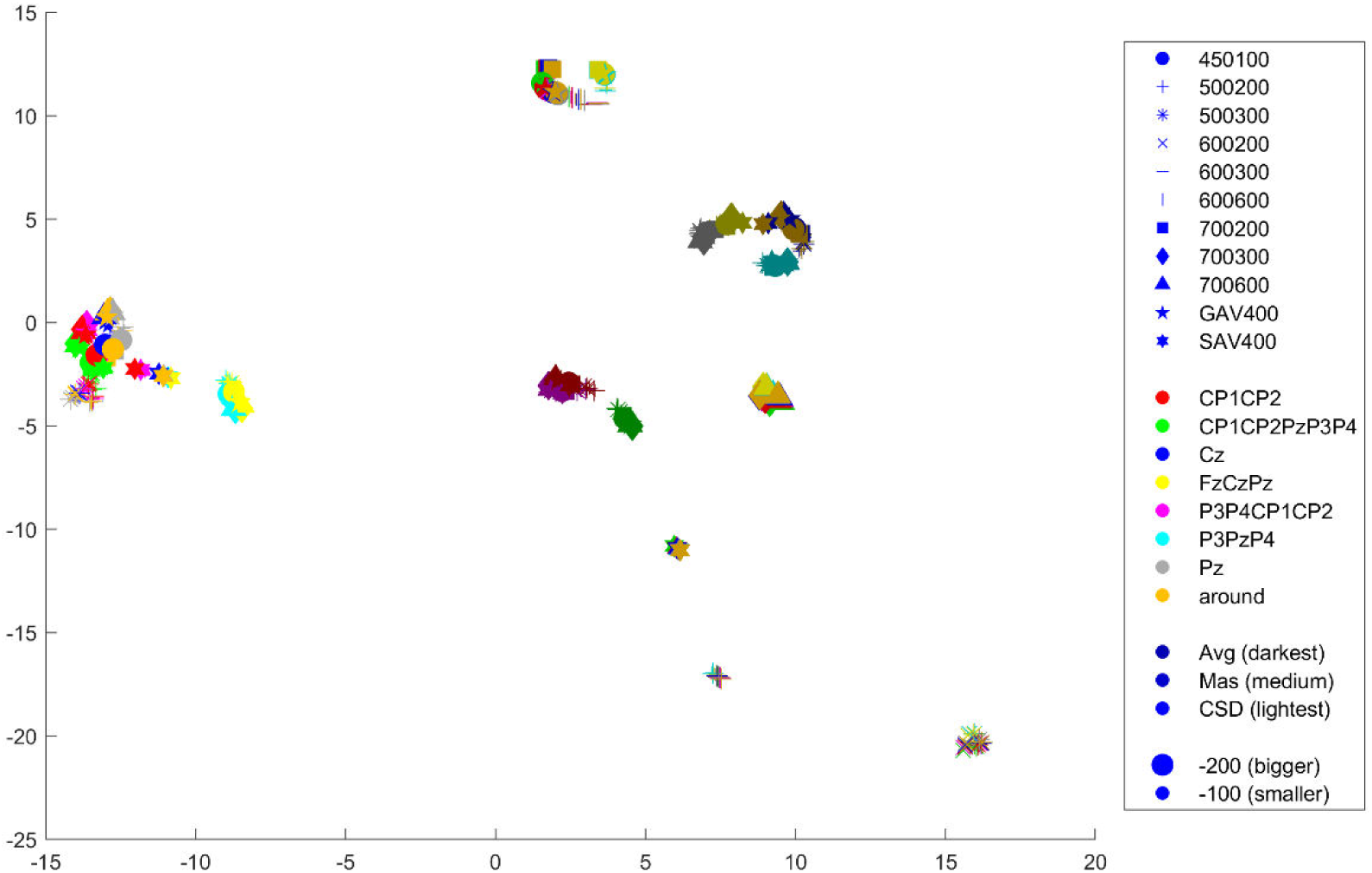
Uniform Manifold Approximation and Projection (UMAP), n_neighbours 15

**Figure 8:**
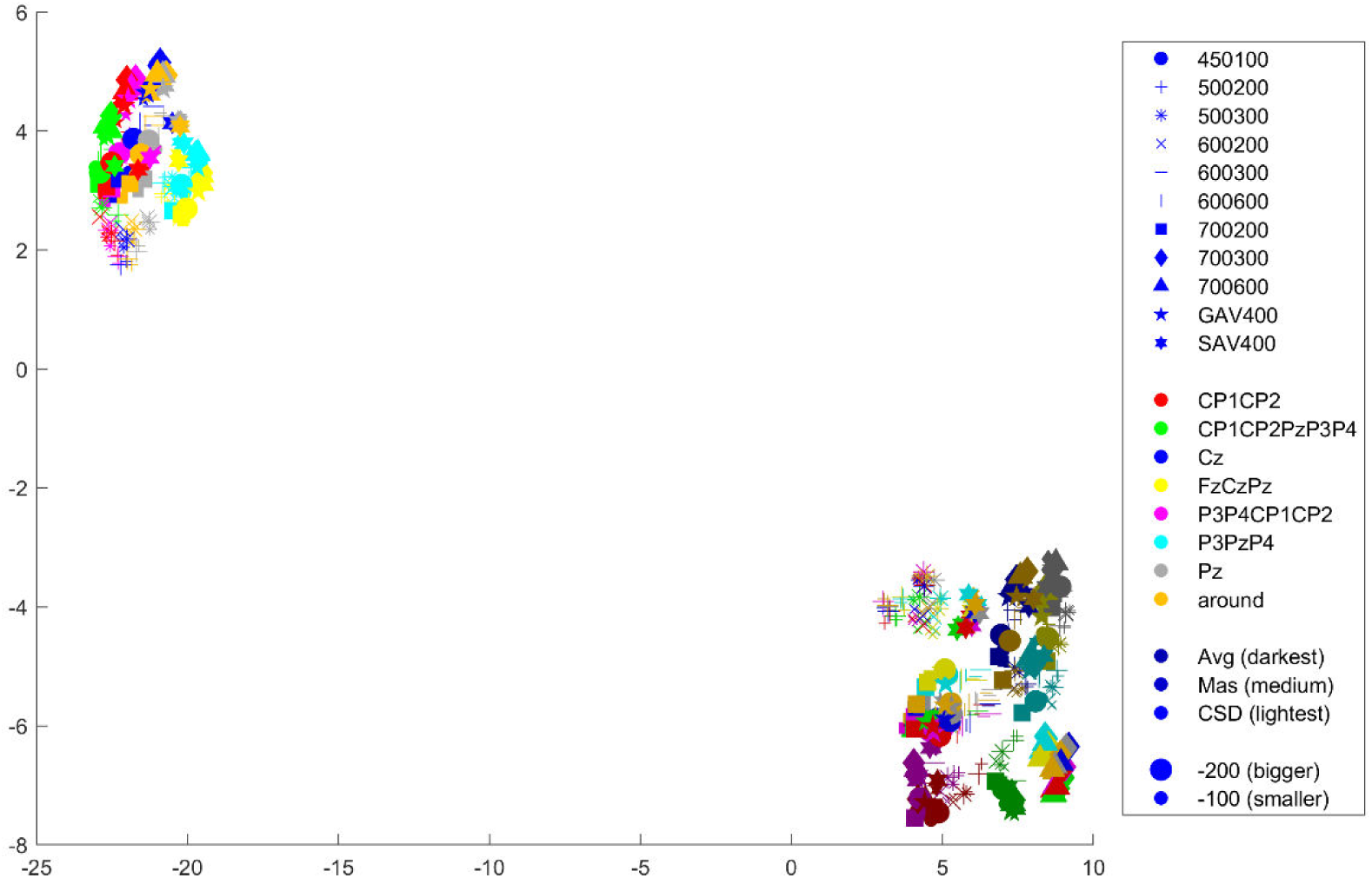
UMAP, n_neighbours 100

**Figure 9:**
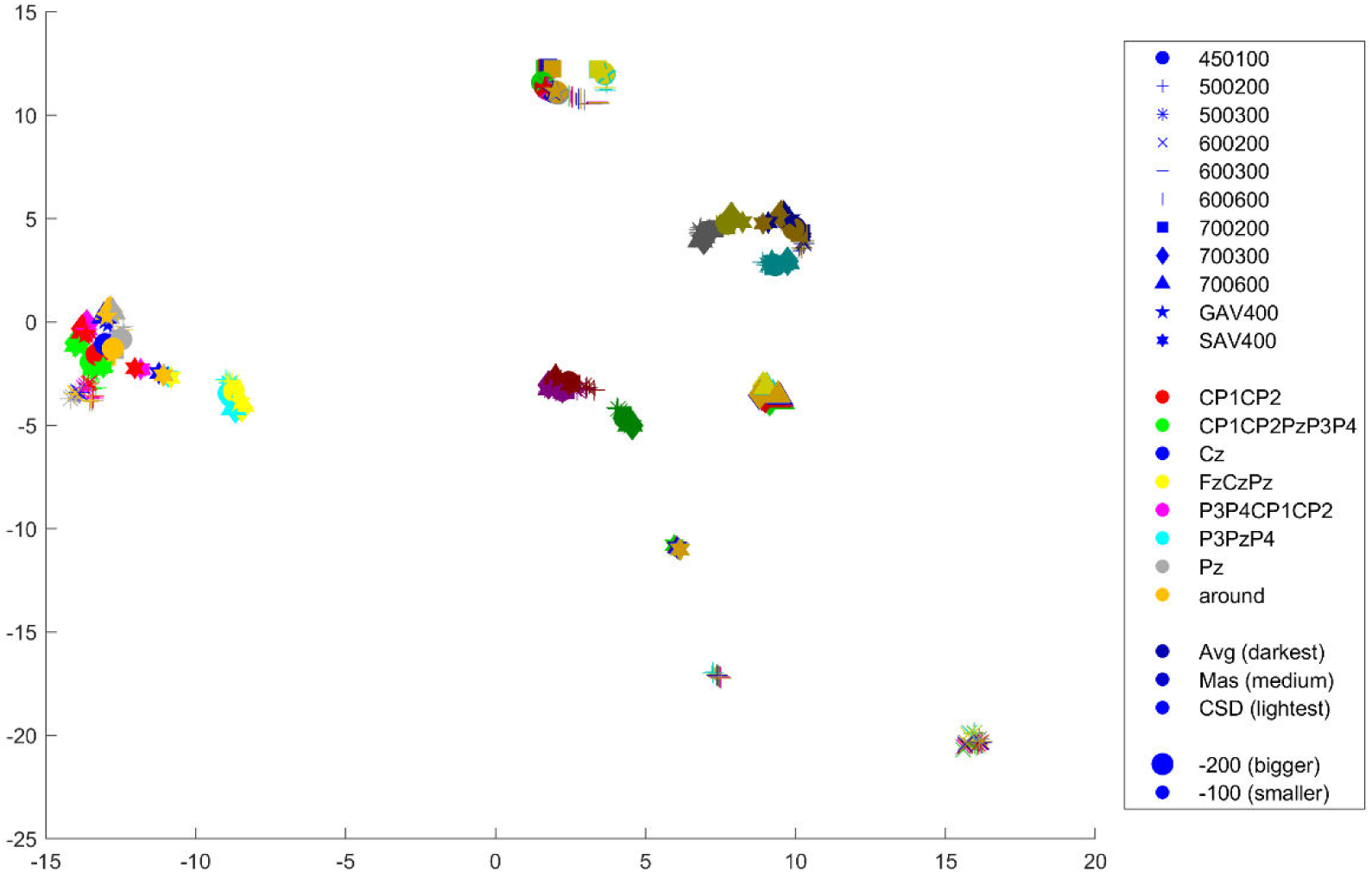
UMAP, minimum distance 0.1

**Figure 10:**
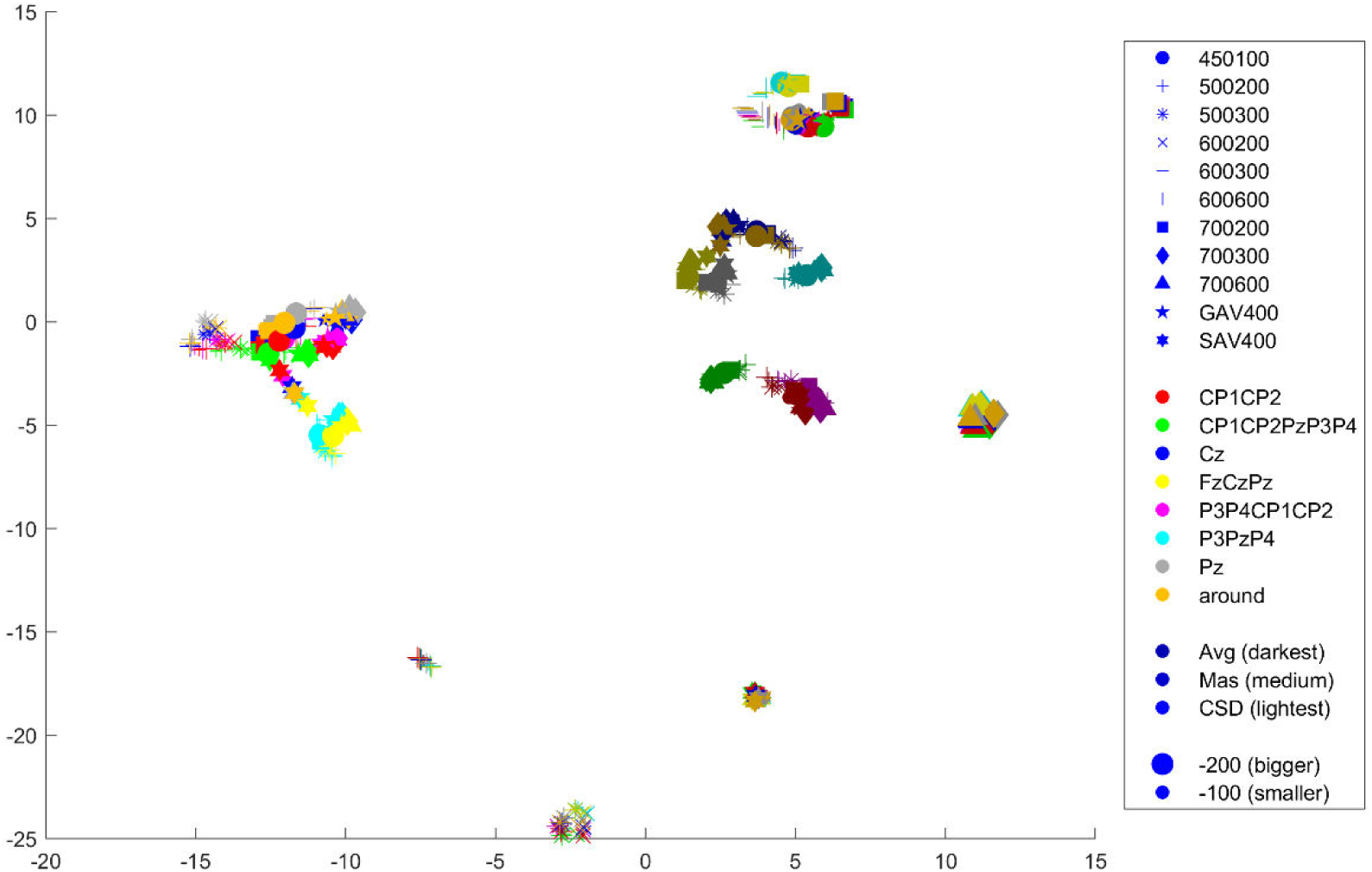
UMAP, minimum distance 0.3

**Figure 11:**
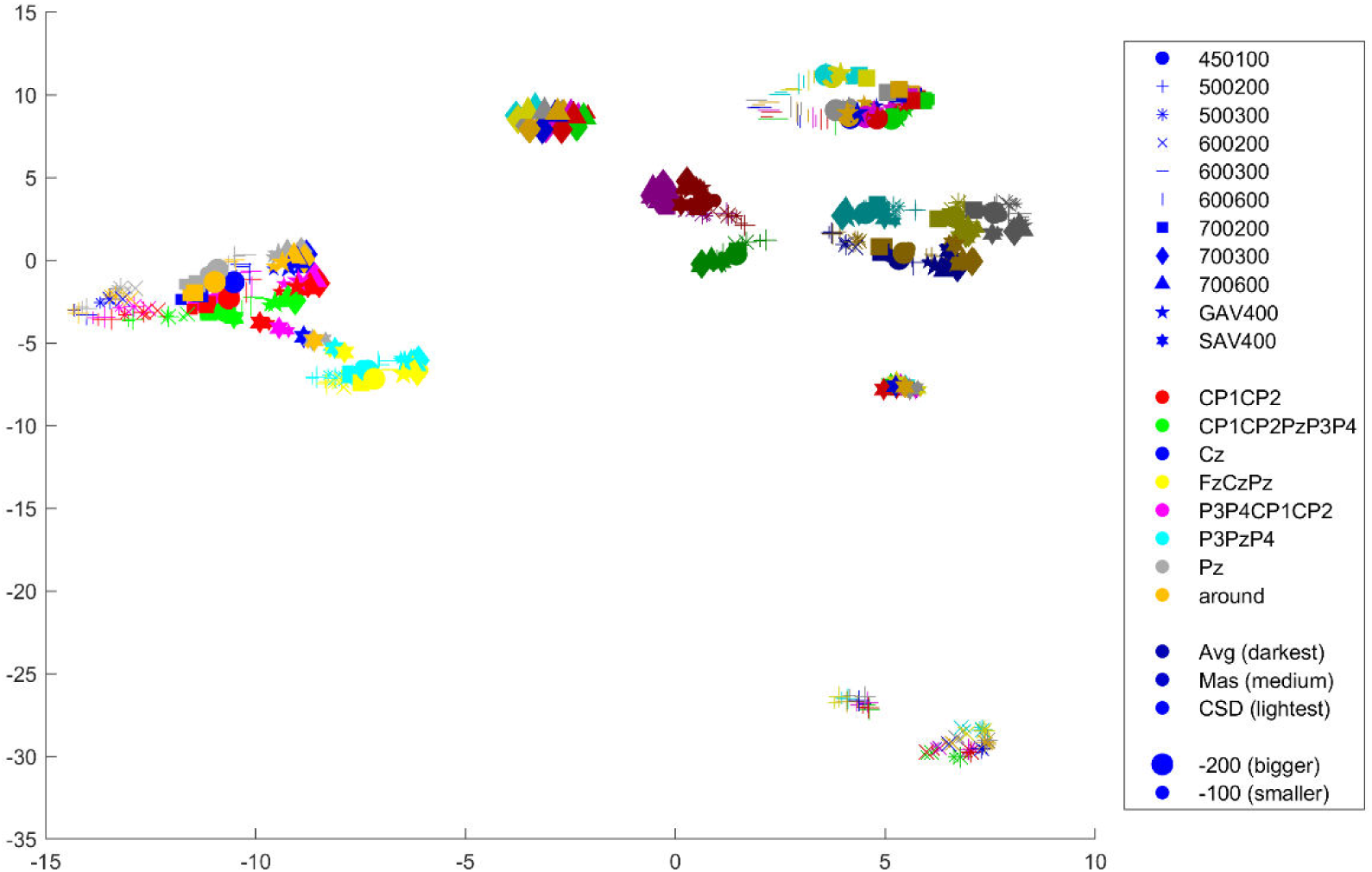
UMAP, minimum distance 0.5

Data matrices used for plotting were generated in script ‘Four_LPPMultiverse_DimensionReductionForActiveLearning.m’, and plots were created in script ‘Five_LPPMultiverse_VisualiseDimensionReduction.m’.

**All figures show the distribution of all 528 pipelines with respect to pairwise Euclidean distances in the LPP difference scores across the condition vector, in the two-dimensional space after embedding using the method in the plot title. *N* = 20 participants (the training sample taken from the full sample of 98 participants).**

## Supplementary 5 Outputs for the 53 pipeline (10%) and 79 pipeline (15%) samples

### Output 1: Median Predictive Accuracy and Kolmogorov-Smirnov Statistic

#### Sample Size = 53 pipelines (10% of full multiverse)

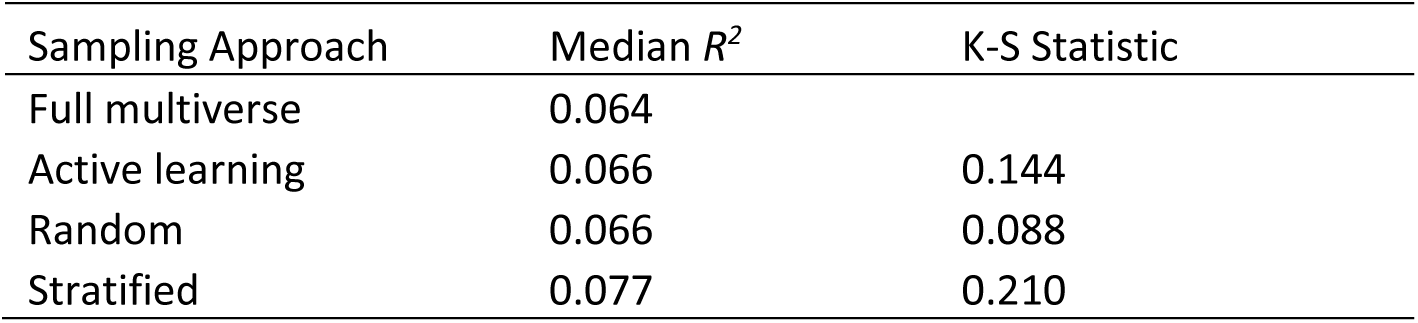

| Sampling Approach | Median $R^2$ | K-S Statistic |
| --- | --- | --- |
| Full multiverse | 0.064 |  |
| Active learning | 0.066 | 0.144 |
| Random | 0.066 | 0.088 |
| Stratified | 0.077 | 0.210 |

#### Sample Size = 79 pipelines (15% of full multiverse)

| Sampling Approach | Median $R^2$ | K-S Statistic |
| --- | --- | --- |
| Full multiverse | 0.064 |  |
| Active learning | 0.065 | 0.093 |
| Random | 0.051 | 0.154 |
| Stratified | 0.067 | 0.068 |

### Output 2: Predictive Accuracy of the Pipelines

#### Sample Size = 53 pipelines (10% of full multiverse)

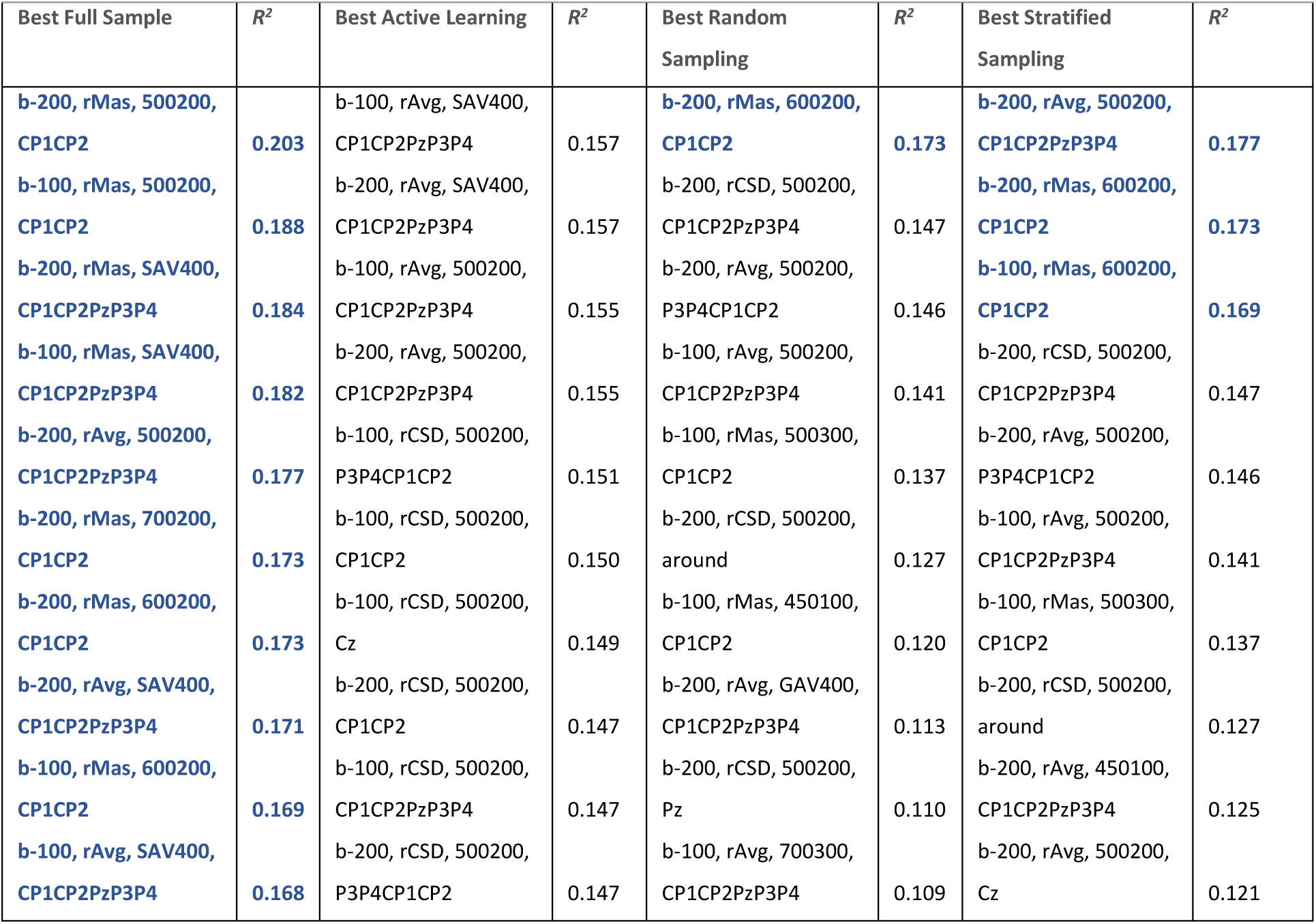

#### Sample Size = 79 pipelines (15% of full multiverse)

| Best Full Sample | $R^2$ | Best Active Learning | $R^2$ | Best Random Sampling | $R^2$ | Best Stratified Sampling | $R^2$ |
| --- | --- | --- | --- | --- | --- | --- | --- |
| b-200, rMas, 500200, CP1CP2 | 0.203 | b-100, rMas, SAV400, CP1CP2PzP3P4 | 0.160 | b-100, rCSD, 500200, P3P4CP1CP2 | 0.156 | b-200, rMas, 600200, CP1CP2 | 0.173 |
| b-100, rMas, 500200, CP1CP2 | 0.188 | b-100, rAvg, SAV400, CP1CP2PzP3P4 | 0.157 | b-100, rCSD, 500200, around | 0.152 | b-100, rCSD, 500200, CP1CP2 | 0.162 |
| b-200, rMas, SAV400, CP1CP2PzP3P4 | 0.184 | b-200, rAvg, SAV400, CP1CP2PzP3P4 | 0.157 | b-200, rAvg, 700200, CP1CP2PzP3P4 | 0.152 | b-200, rCSD, 500200, CP1CP2PzP3P4 | 0.147 |
| b-100, rMas, SAV400, CP1CP2PzP3P4 | 0.182 | b-100, rAvg, 500200, CP1CP2PzP3P4 | 0.156 | b-200, rMas, SAV400, CP1CP2 | 0.147 | b-200, rCSD, 500200, CP1CP2 | 0.142 |
| b-200, rAvg, 500200, CP1CP2PzP3P4 | 0.177 | b-200, rAvg, 500200, CP1CP2PzP3P4 | 0.156 | b-200, rMas, 700200, P3P4CP1CP2 | 0.137 | b-100, rAvg, 500200, CP1CP2 | 0.138 |
| b-200, rMas, 700200, CP1CP2 | 0.173 | b-100, rCSD, 500200, P3P4CP1CP2 | 0.152 | b-200, rMas, 600200, P3P4CP1CP2 | 0.131 | b-100, rMas, 500300, CP1CP2 | 0.137 |
| b-200, rMas, 600200, CP1CP2 | 0.173 | b-100, rCSD, 500200, CP1CP2 | 0.152 | b-200, rAvg, 700200, P3P4CP1CP2 | 0.116 | b-200, rCSD, 500200, Cz | 0.128 |
| b-200, rAvg, SAV400, CP1CP2PzP3P4 | 0.171 | b-100, rCSD, 500200, Cz | 0.150 | b-100, rAvg, 600200, CP1CP2PzP3P4 | 0.115 | b-200, rCSD, 500200, around | 0.127 |
| b-100, rMas, 600200, CP1CP2 | 0.169 | b-200, rMas, SAV400, CP1CP2PzP3P4 | 0.149 | b-100, rAvg, SAV400, P3PzP4 | 0.103 | b-200, rAvg, 500200, around | 0.123 |
| b-100, rAvg, SAV400, CP1CP2PzP3P4 | 0.168 | b-100, rCSD, 500200, CP1CP2PzP3P4 | 0.149 | b-200, rAvg, 700600, CP1CP2 | 0.099 | b-100, rAvg, 500200, P3P4CP1CP2 | 0.121 |

Blue = the Pipeline is included within the best pipelines of the full multiverse. b-100 and b-200 = baseline durations of 100 and 200 ms respectively, rAvg, rMas and rCSD = common average, linked mastoids and common source density reference schemes respectively.

### Output 3: Raincloud Plots of Predictive Accuracies

#### Sample Size = 53 pipelines (10% of full multiverse)

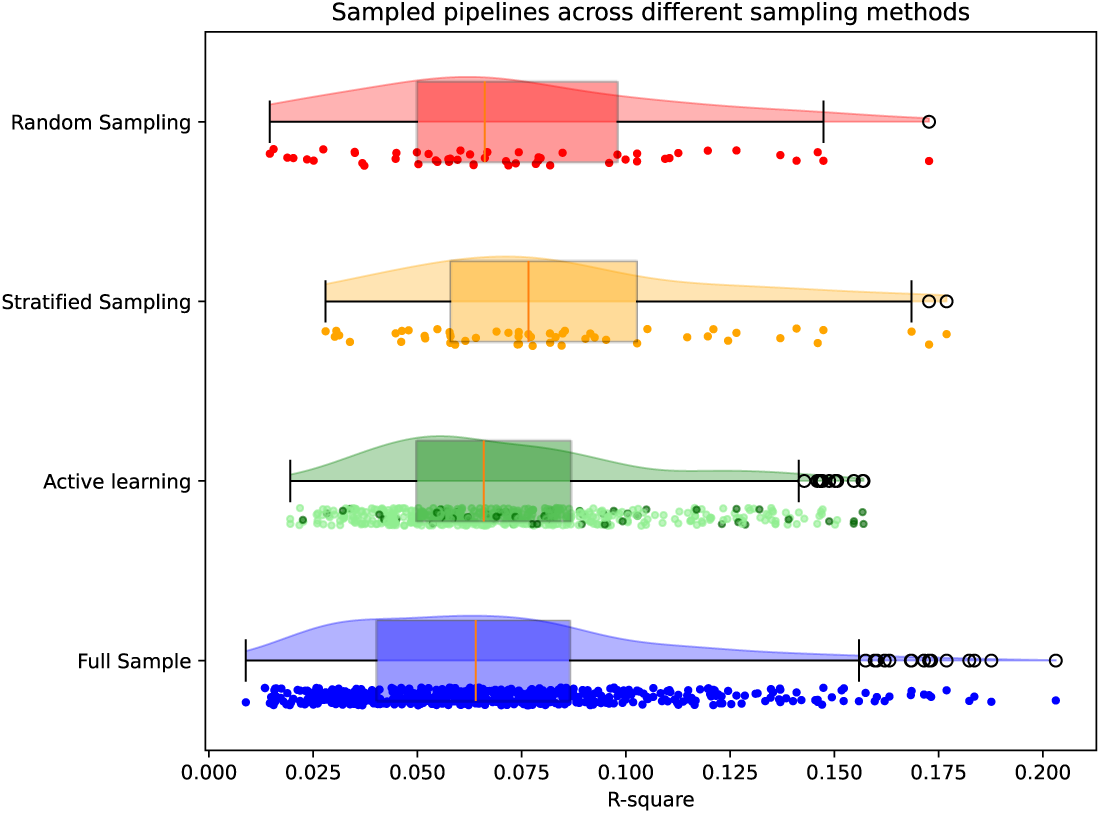

#### Sample Size = 79 pipelines (15% of full multiverse)

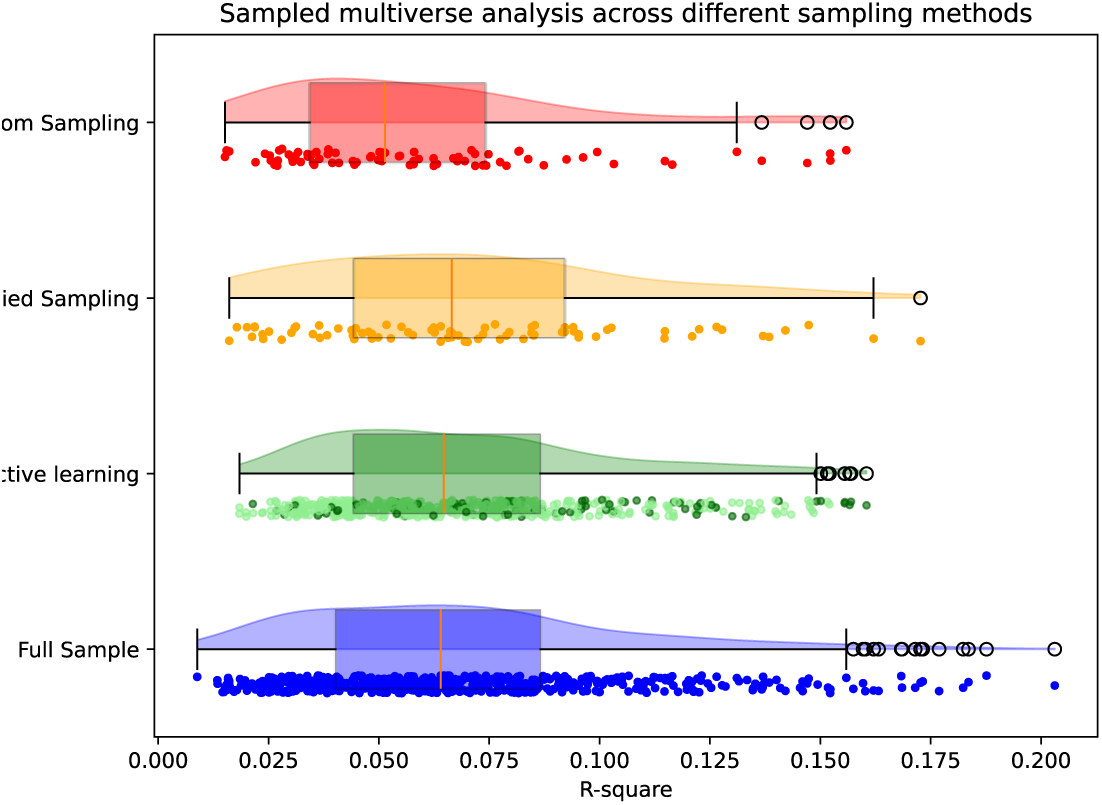

### Output 4: Scatter Plots of Spatial Distribution in the Low Dimensional Space

#### Sample Size = 53 pipelines (10% of full multiverse)

| Sample Size | Full Multiverse vs. Random<br>Sample | Full Multiverse vs.<br>Stratified Sample | Full Multiverse vs. Active<br>Learning Sample |
| --- | --- | --- | --- |
| 53 pipelines | 70.90 | 72.53 | 70.78 |

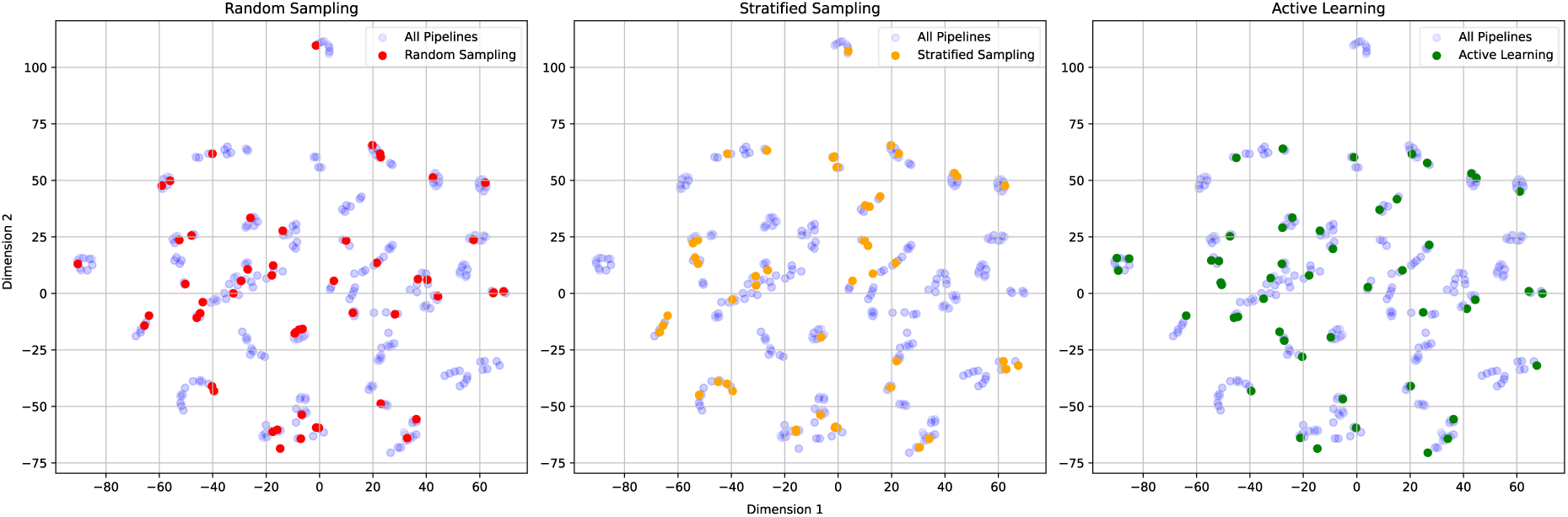

#### Sample Size = 79 pipelines (15% of full multiverse)

| Sample Size | Full Multiverse vs. Random<br>Sample | Full Multiverse vs.<br>Stratified Sample | Full Multiverse vs. Active<br>Learning Sample |
| --- | --- | --- | --- |
| 79 pipelines | 69.09 | 70.92 | 71.63 |

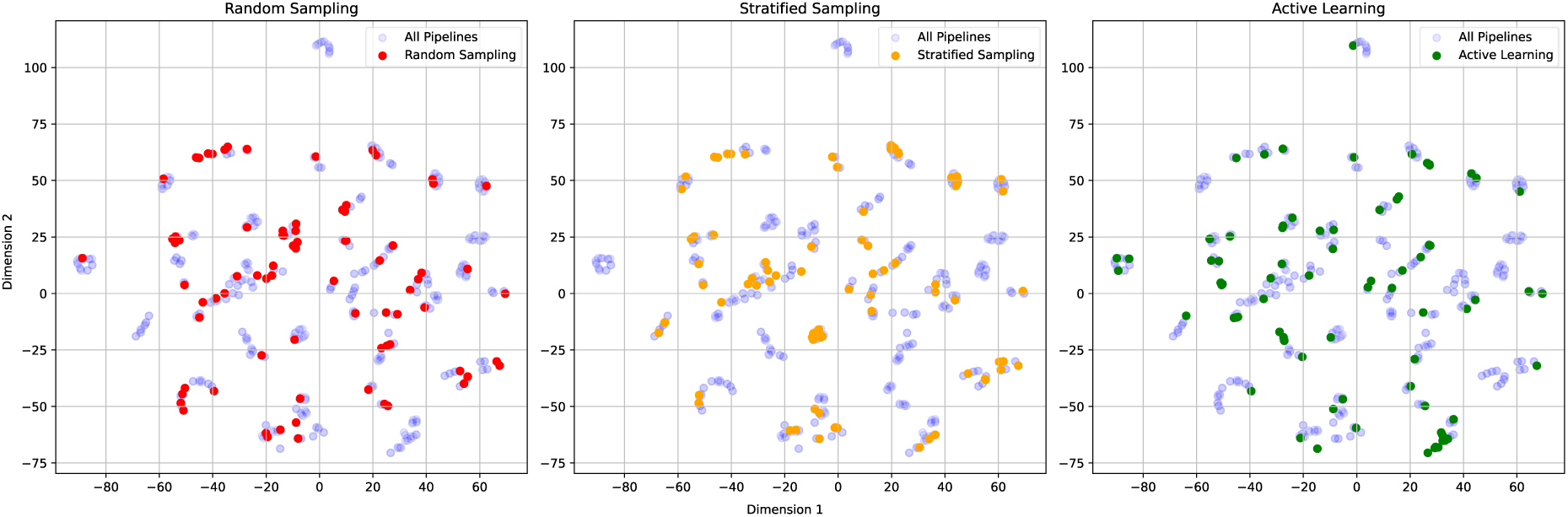

### Output 5: Specification Curves

#### Sample Size = 53 pipelines (10% of full multiverse)

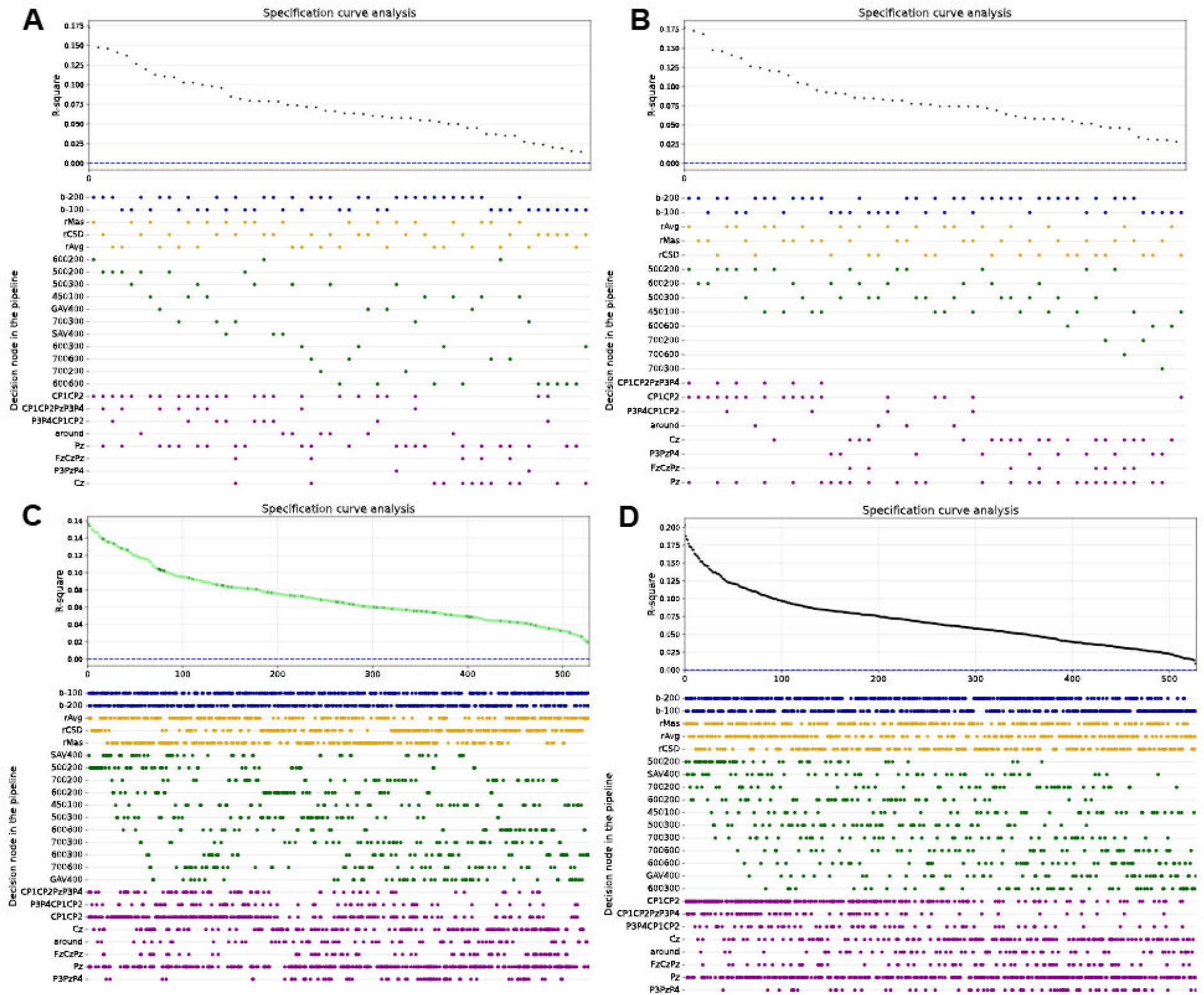

Specification curves displaying the variability in the *R^2^* across each sample, in vertical alignment with the respective pipeline options. Panel A = random sample, Panel B = stratified sample, Panel C = active learning sample, Panel D = full multiverse. Each colour in the lower specification panel corresponds to one decision node. Blue = baseline duration, yellow = reference scheme, green = time window, purple = electrode cluster. Each row within a colour represents a different option at that decision node. In the top panel of the active learning plot, dark green points denote pipelines that were directly sampled and light green points denote pipelines that were estimated.

#### Sample Size = 79 pipelines (15% of full multiverse)

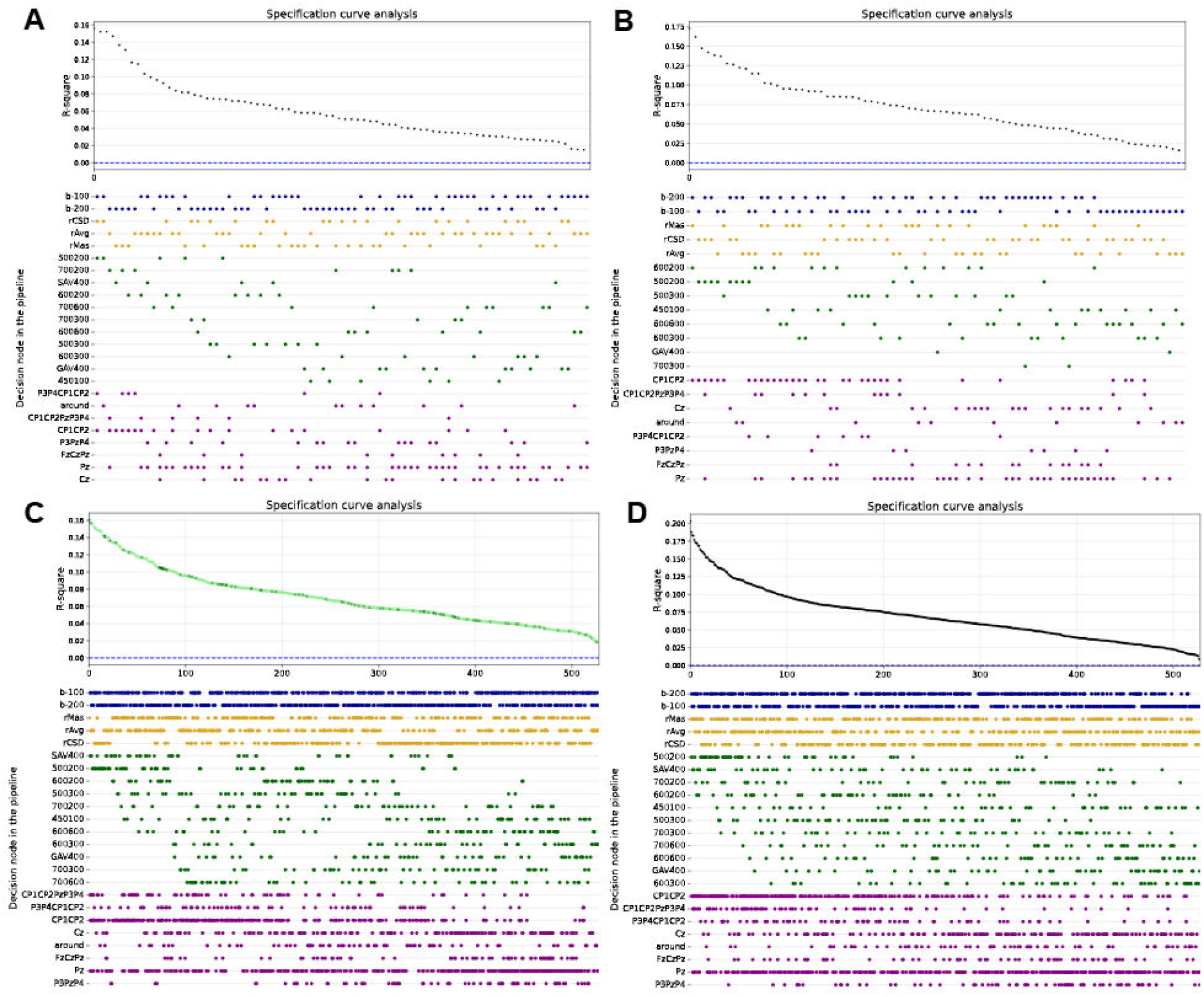

Specification curves displaying the variability in the *R^2^* across each sample, in vertical alignment with the respective pipeline options. Panel A = random sample, Panel B = stratified sample, Panel C = active learning sample, Panel D = full multiverse. Each colour in the lower specification panel corresponds to one decision node. Blue = baseline duration, yellow = reference scheme, green = time window, purple = electrode cluster. Each row within a colour represents a different option at that decision node. In the top panel of the active learning plot, dark green points denote pipelines that were directly sampled and light green points denote pipelines that were estimated.

### Baseline

The alternative baseline durations, which refer to the pre-stimulus time window in which the voltage is subtracted from the post-stimulus waveform, were (1) −100 ms to 0 ms and (2) −200 ms to 0 ms. Each of the alternative baseline durations were considered appropriate for the research design, as we expect no stimulus-related cognitive or behavior induced potentials in either of these time windows. As the baseline voltage is subtracted from the entire waveform, essentially making the post-stimulus period a difference score between the pre- and post-stimulus periods, any noise or task-related cortical activity in the baseline voltage will influence the post-stimulus amplitudes. The larger the baseline window, the greater the opportunity for noise within it, and the more likely it is contaminated with stimulus-related cortical activity. On the other hand, the larger the baseline window, the more likely that the averaging will decrease this influence.

### Offline reference

The alternative offline reference schemes were (1) linked mastoids, (2) common average reference (CAV), and (3) current source density (CSD). using the electrode coordinates. The vertex (Cz) was deemed unsuitable given that this electrode and those in its spatial proximity are electrodes of interest for quantifying the LPP. Different reference schemes may influence the amplitude of ERP components by altering the relative weighting of electrode signals. For example, depending on the density of electrode coverage around the region of the electrode(s) of interest, the amplitude of an ERP of interest may not be well represented in the CAV dataset.

### Time window

The alternative time windows were (1) 400-500ms, (2) 400-600ms, (3) 350-650ms, (4) 500-700ms, (5) 450-750ms, (6) 300-900ms, (7) 600-800ms, (8) 550-850ms, (9) 400-1000ms, (10) +/− 200ms around the grand average within 300-1000ms, (11) +/− 200ms around the single subject average within 300-1000ms. On the one hand, a larger time window is advantageous for capturing the broad LPP component despite individual differences in latency, which may be particularly poignant when the expression stimulus appears gradually over a dynamic video. On the other hand, a larger time window increases the likelihood that the LPP estimate may be distorted by other cognitively distinct components occurring during that span of time.

### Electrodes

The alternative electrodes of interest were (1) CP1, CP2, Pz, P3, P4; (2) P3, P4, CP1, CP2; (3) P3, Pz, P4; (4) Fz, Cz, Pz; (5) CP1, CP2; (6) Cz; (7) Pz; (8) four electrodes around the midline peak. It is possible that broader electrode clusters may dilute localized signals, whereas smaller, more targeted clusters may enhance them. On the other hand, broader electrode clusters may be more accommodating to between-person variability in signal propagation more so than smaller, more localized clusters.

